# Modelling Osteoarthritis pathogenesis through Mechanical Loading in an Osteochondral Unit-on-Chip

**DOI:** 10.1101/2023.08.29.555292

**Authors:** Andrea Mainardi, Anastasiya Börsch, Paola Occhetta, Robert Ivanek, Martin Ehrbar, Lisa Krattiger, Philipp Oertle, Marko Loparic, Ivan Martin, Marco Rasponi, Andrea Barbero

## Abstract

A cure for osteoarthritis (OA), the most prevalent musculoskeletal disease, remains an unmet need. Investigating the molecular and cellular processes leading to OA is challenged by the absence of human models that capture the complex interplay among different tissues in the joint under pathophysiological mechanical loading.

In this study, we have engineered an OsteoChondral Unit (OCU)-on-chip system where composite hyaline cartilage - mineralized osseous microtissue analogues are exposed to controlled, tissue-specific compression regimes akin to those of the OCU *in vivo*. Through single-cell transcriptomic analysis, we demonstrate the critical relevance of the mineralized layer in inducing chondrocyte subpopulations implicated in the progression of OA.

Upon exposure to hyperphysiological loading, the OCU-on-chip captures early phenotypic traits of OA pathogenesis, comprising alterations of subchondral mineral content and acquisition of previously described OA genetic signatures.

This system enabled to identify novel upstream drivers of OA metabolic changes, including mechanically induced ribosomal alterations, as well as associated molecular targets towards the development of disease-modifying OA therapies.

## Introduction

Osteoarthritis (OA) is a whole joint degenerative disease with no available pharmacological cure^1,2^. Pathological alterations include cartilage erosion but also, beside synovial inflammation^3^, subchondral bone modifications^4,5^; the entire osteochondral unit (OCU) is affected^1^. Cause-effect mechanisms of these processes and potential cross-talk among the different joint structures are nevertheless still not understood and the precise origin of the disorder remains unknown.

Although OA is recognised as a multifactorial pathology with molecular, genetic, and environmental influences, there is consensus concerning its correlation with mechanical risk factors such as trauma, obesity, or joint misalignment^6,7^, thus warranting further studies in this direction.

Involvement of multiple tissues and joints’ mechanically active environment introduce nevertheless a multiplicity of confounding factors in the effort to pinpoint the origin of the pathology.

Besides, while different chondrocytes subpopulations have recently been identified and demonstrated to be affected by OA^8,9^, we still need to comprehend how a specific transcription signature correlates with the cells’ response to pathological stimuli, and how these subpopulations are affected by cartilage/bone interactions, possibly mediating the progression of OA^10^. Innovative preclinical OA models where a sufficient level of complexity goes hand in hand with a strict control over the experimental environment could be invaluable in these matters.

We previously introduced a Cartilage-on-Chip (CoC) OA model demonstrating that a 30% hyperphysiological compression (HPC) leads to recapitulation of OA traits encompassing inflammation, a shift in cartilage homeostasis towards catabolism, and the onset of an hypertrophic phenotype^11^.

Understanding the root causes of OA requires, however, to scrutinize the interplay among different joint compartments, such as cartilage and subchondral bone^12^. Engineering an *in vitro* model that provides mechanical stimuli in the range of *in vivo* pathophysiological stimulation for both cartilage and bone is, nevertheless, not trivial. In native joints, the OCU response to loading is mediated by tissues with a complex hierarchical organization^13^ and with Young’s moduli ranging from a few MPa to GPa^14^. Given the higher mechanical properties of calcified cartilage^15^ and subchondral bone^14,16^, these tissues are reasonably subjected to strain levels which are at least one order of magnitude lower than those experienced by hyaline cartilage. No existing *in vitro* model addresses this issue.

In this work we aimed at devising an OCU-on-chip that recapitulates OA phenotype through tissue-specific mechanical loading so to suggest effective targets for disease modifying therapies.

To this end we developed a microscale platform where - through a new microfluidic concept, namely the Vertical capillary Burst Valve (VBV) - bi-phasic microconstructs with a spatial organization that mimics the OCU functional anatomy can be subjected to tissue specific, physiologically relevant, mechanical compression. With this technology, we achieved OCU-like structures constituted by a cartilaginous and a mineralized subchondral layer, and determined how the interplay between the two tissues influences their response to stimuli. Finally, we exploited our model and single cell RNA sequencing (scRNA-seq) to decipher how the co-culture between cartilage and bone impacts chondrocytes subpopulations variety and, additionally, provided a comprehensive assessment of the transcription machinery activated, in these subpopulations, by aberrant mechanical loading.

## Results and Discussion

### A device for mechanical compression of bi-layered 3D microtissues

To mimic the basic functions of the OCU we conceived a microscale device which can host two directly interfaced 3D cellular constructs, namely a cartilaginous tissue and a subchondral tissue, and subject them to tissue-specific mechanical compression (Fig. 1a).

**Fig. 1.**
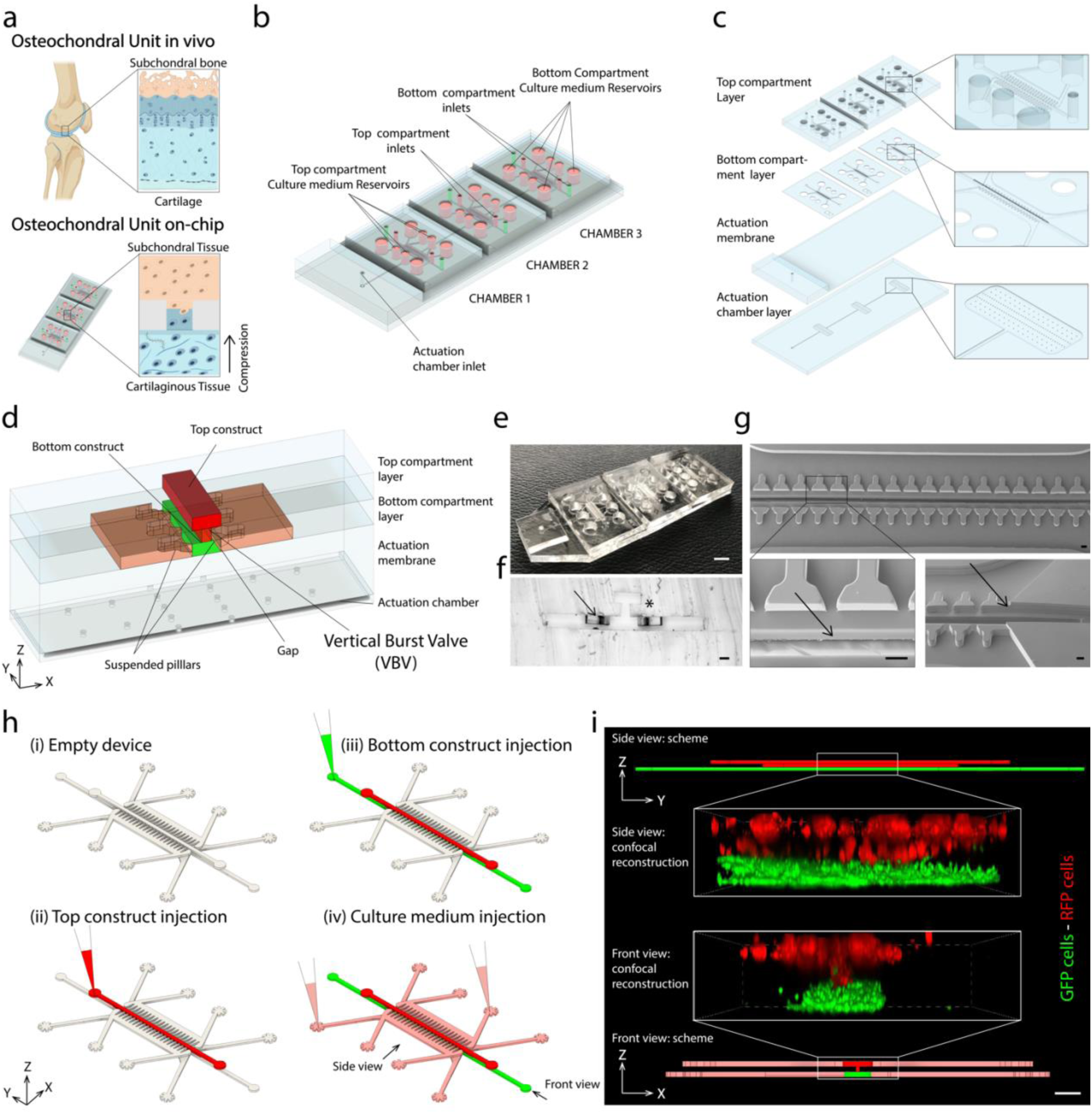
A microfluidic device providing bi-layer 3D constructs with tissue specific mechanical compression. **a**, Schematization of the knee OCU *in vivo* and of the OCU-on-chip which includes both a cartilaginous tissue and a subchondral tissue that have a direct interface and are subjected to tissue specific mechanical compression. **b,** Schematization of the PDMS device. Each platform is constituted by three culture chambers. **c,** Sketch of the 4 PDMS layers composing the device. A Thick PDMS slab (height 3 mm) is added on the actuation membrane to increase the retention of the tube providing pressure to the actuation compartment. **d,** Schematic detailing the inner structure of the device were the two 3D constructs (the top represented in red, the bottom in green), the gap between T shaped pillars and actuation membraned, and the VBV geometry can be appreciated (the top compartment is representative of the NDG configuration). **e,** Photograph of a fabricated device with the DG configuration. Scale bar, 4 mm. **f,** Photograph of a device section (NDG configuration), the arrow points to the suspended pillars, the asterisk indicates the VBV geometry. Scale bar, 100 𝜇m **g,** SEM images of the device bottom compartment observed from below. Arrows point to the necking connecting bottom and top compartment (left) and to the gab below the T shaped pillars (right). Scale bar, 100 𝜇m. **h,** Schematic of the device injection procedure. Either top or bottom compartments can be filled first, knowing that the cell laden hydrogel injected as former will also fill the necking area. **i,** Confocal microscopy reconstruction of cell laden constructs seeded in the two culture compartments. GFP labelled hACs are depicted in green, RFP labelled HUVECs are represented in red. The two tissues are in direct contact for 1:3 of their width (along the x direction) and for the whole tissue length (along the Y direction). Scale bar, 100 μm. 3D schematics are added at the top and at the bottom of the image. The necking area presents the only contact point between top and bottom compartment as can be observed in the XZ plane highlighting the tilted-H VBV geometry.

The microfluidic system comprises three independent functional units (i.e. chambers), each being constituted by a bottom compartment for the cartilaginous tissue, and a top compartment for the subchondral tissue. All functional units are connected to the same pneumatic actuation chamber adopted as a mechanism to apply mechanical compression through the deformation of an overlaying actuation membrane (Fig. 1b, c).

The device was realized assembling four layers as depicted in Fig. 1c. All components of the device were manufactured in polydimethylsiloxane (PDMS) following the procedure described in Supplementary Fig. 1.

The bottom compartment is based on a platform that we previously introduced to mechanically stimulate 3D cartilaginous microconstructs^11^. It is constituted of a central channel that hosts a 3D cell laden hydrogel (height 143 𝜇m, width 300 𝜇m, represented in green in Fig. 1d) divided by two rows of overhanging pillars from lateral culture medium channels (in orange in Fig. 1d). A gap is present between the bottom surface of the pillars and the top surface of the actuation membrane. Upon pressurization of the actuation chamber, the membrane bends upwards until it abuts against the pillars’ bottom surface. The ratio between the height of the pillars (100 𝜇m) and that of the gap (43 𝜇m) defines the entity of the compression. In this case, the device was designed to provide a precise 30% hyperphysiological compression (HPC) to the 3D tissue hosted in the bottom compartment. We previously demonstrated that such stimulus is sufficient to elicit an OA-like phenotype in cartilaginous microconstructs^11^. Notably, the pillars’ T-shaped cross section was chosen to minimize posts’ outward bending upon microconstruct compression due to the increase in generated pressure. The distance between two adjacent pillars (30 𝜇m) was selected to confine the 3D construct during injection and polymerization phases, beyond limiting its lateral expansion during compression.

Differently from our single-culture formerly presented platforms^11,17^, the device in this study was tailored to obtain a bi-layer construct resembling the OCU where hyaline cartilage is interfaced with subchondral bone via the calcified cartilage, one of the structures more involved during the progression of OA^418^. To this end, an additional cell culture top compartment was introduced in the platform. The top compartment (in red in Fig. 1d) is constituted either by a simple channel to host the subchondral tissue (height 150 𝜇m, width 300 𝜇m, Fig. 1d) or by a central channel connected in different ways to culture medium channels and reservoirs (Extended Data Fig. 1 a-e). Three versions were introduced: a non-diffusive geometry (NDG), a lowly diffusive geometry (LDG), and a diffusive geometry (DG) depending on the presence (or absence), and the entity, of an interface area between the central channel and lateral culture medium channels.

Top and bottom compartments are linked through a necking (width 100 𝜇m, height 150 𝜇m) in the culture chamber geometry, which is part of the bottom compartment PDMS layer. The assembly constituted by top and bottom compartments’ central channels, which has the cross section of a tilted H, constitutes a capillary burst valve (CBV) rotated of 90°. The complete structure can be observed in Fig. 1d, where the NDG configuration is represented. CBVs are strictions within microfluidic channels providing a pinning interface for a forward proceeding fluid^19–21^. Classically, CBVs were designed to stop an advancing front from going further on, left or right^21,22^. (Supplementary Fig. 2a). Owing to the limited role played by volumetric forces at the microscale (e.g. gravity) we hypothesizes that, flipping by 90° such CBV configuration, it was feasible to obtain a Vertical capillary Burst Valve (VBV) designed for stopping an advancing fluid from moving upward or downward (i.e. along the Z axis in Supplementary Fig. 2b).

Integrating the VBV with the above-described compression mechanism allows to apply a mechanical stimulus to a bilayer construct. The necking reduces the interface area between the cartilaginous and the subchondral tissue to one third (its width being 100 𝜇m) but it also creates a discontinuity in the strain field of the two constructs. The compression of the cartilaginous construct is determined by the actuation membrane stroke, while the subchondral constructs deformation is limited to very low strains by the PDMS structures. Such a strain distribution reflects the OCU deformation during gait where a high compressive strain of the cartilage surface corresponds to very limited deformations of the underlying calcified cartilage and subchondral bone^23^. A picture of a fabricated device is represented in Fig. 1e, while Fig. 1f depicts a device section with the VBV geometry. Details of the bottom compartment from below can also be viewed in the scanning electron microscope (SEM) images in Fig. 1g. In all cases a high adherence of the fabricated device to the nominal geometry was achieved.

The adopted fabrication methodology (Fig. 1c, Supplementary Fig. 1) requires a precise and accurate alignment of the PDMS layers. Various dimensions of the top compartment central channel were evaluated to maximise the ease and precision of the alignment during fabrication (Supplementary materials, Supplementary Fig. 3). In the final version both bottom and top compartments have a width of 300 𝜇m.

The functionality of the VBV concept and of the platform in general were validated. Firstly, we showed that a bi-layer tissue can be obtained with high reproducibility by simply injecting two cell laden hydrogels in the top and bottom compartments and maintaining them confined to their respective sections (Fig. 1h). More than 80% of devices (n> 100 devices) could successfully be injected. This efficacy is in agreement with the 90% injection efficiency of our CoC device considering that both hydrogel injections have this success rate. Fig. 1i shows a confocal microscopy reconstruction of red fluorescent protein (RFP) expressing human umbilical vein endothelial cells (HUVECs) and green fluorescent protein (GFP) expressing human articular chondrocytes (hACs) embedded in an enzymatically cross-linkable and degradable 2% w/v polyethyleneglycole (PEG) based hydrogel^24^ and seeded, respectively, in top and bottom compartments. Both images in the YZ and in the XZ planes highlighting the direct contact between cells in the two constructs.

Secondly, the pressure necessary to achieve contact between actuation membrane and T-shaped pillars’ bottom surface, i.e 0.4 Atm, was determined experimentally as previously described^11,25^ (Supplementary materials, Supplementary Fig. 4).

Finally, following the procedure outlined in Extended Data Fig. 1f, we determined the diffusion kinetic between the bottom and top compartments of solutes with a molecular weight range of known morphogens^25^. The assessment was performed considering DG and LDG configurations while this was not possible with the NDG that does not have medium channels in the top compartment (Extended Data Fig. 1g). Notably, the LDG (combined with a low cartilaginous construct permeability reached after 14 days of hACs constructs maturation) limits the inter-compartments diffusion of molecules with a molecular weight in the range of common growth factors (Extended Data Fig. 1h, i). Within a limited time span, it is consequently possible to distinguish from which compartment a given molecule was secreted.

These data demonstrate the successful fabrication of the envisioned platform and the functionality of the VBV concept. Our multi-layer fabrication approach, furthermore, allows to tailor the different modules for specific purposes thus making the strategy that we devised for the OCU on-chip model suitable for other applications.

### Providing 3D microconstructs with compartment specific compression levels: numerical model

To test whether the proposed device could effectively provide constructs hosted in top and bottom compartments with well-defined and discrete compression levels, indicative of those experienced by OCU tissues *in vivo*^23^, we estimated their strain field through a Finite Element (FE) numerical model.

Fig. 2a depicts the device minimal repetitive unit. The Region Of Interest (ROI) control volume which was used in computations is represented by the shaded area.

**Fig. 2.**
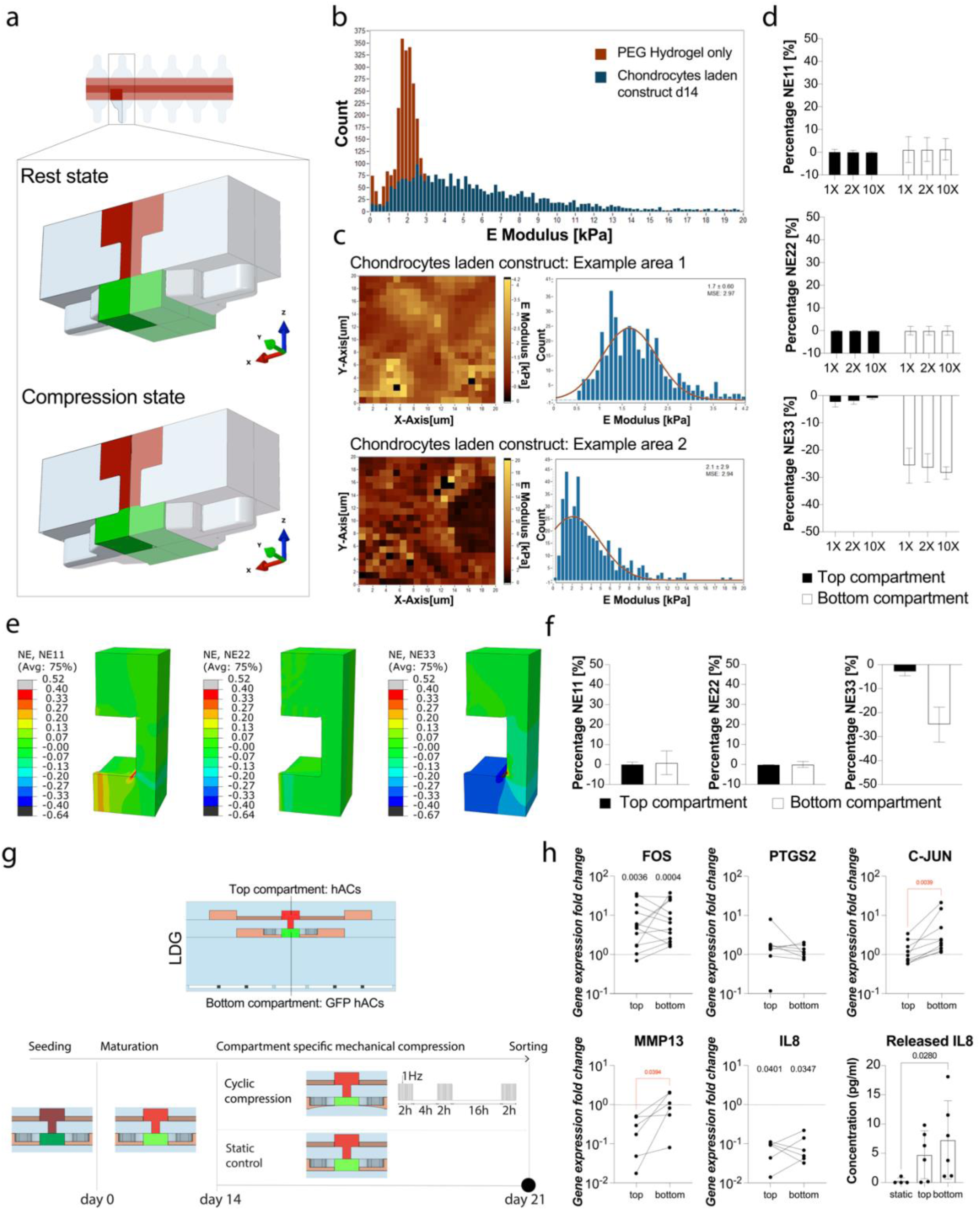
The OCU-on-chip provides compartment specific compression levels: strain field estimation and biological validation. **a**, Schematization of the 3D model adopted for FE simulation (NDG configuration). A top view of the constructs and of the T-shaped pillars is represented in the upper figure; the black rectangle encircles the minimal repetitive unit element while the shaded area represents the Region of interest (ROI) adopted in simulations. Boundary conditions were adopted to reduce the minimal repetitive unit to the selected ROI. 3D FE reconstructions of the minimal repetitive unit in rest state and upon compression are represented at the bottom. The gel filling the top compartment is represented in red, the one filling the bottom compartment is represented in green. **b,** Cartilaginous constructs and PEG hydrogel mechanical properties as determined by IT-AFM. E modulus frequency histograms are referred to empty PEG hydrogels (in dark red) or to PEG hydrogels laden with hACs cultured statically for 14 days within microfluidic devices. (Cartilaginous constructs: n= 3 biologically independent samples from 1 donor, PEG hydrogels only: n=3 independent samples). **c,** Cartilaginous constructs properties varied depending on the indentation location within constructs. Two indentation maps referring to different positions are reported as an example. Local E values are colour coded; frequency histograms of E values are reported on the right. **d,** Compartment specific compression levels are robust with respect to constructs mechanical properties variations as determined by FE estimated nominal strains (NE) along the principal direction (NE11, NE22, and NE33 respectively for strains aligned with the X, Y, and Z axes) as a function of the top compartment E modulus. 1X, 2X, and 10X, refer to the modulus adopted for the hydrogel in the top compartment with respect to the one of the hydrogel in the bottom compartment (i.e. 1.9 kPa as experimentally determined by IT-AFM). The top construct was considered comprehensive of the volume in correspondence of the VBV necking. Values are reported as mean ± s.d. **e, f,** The device applies compartment specific compression levels to bi-layer cartilaginous constructs. Both top and bottom compartments were considered as filled with cartilaginous constructs with E modulus of 3.66. kPa, as determined experimentally by IT-AFM. The strain distribution is reported through the NE evaluated along the principal directions. Lateral and longitudinal expansions are limited by the device geometry. A clear difference in the compression levels (i.e. NE33) of top and bottom compartments is visible from the contour plot as well as from the global NE values reported in **f (**as mean ± s.d.). **g,** The elicitation of different mechano-dependent gene expression modulations in top and bottom compartments was verified experimental as delineated in the schematic. Top and bottom compartments (LDG configuration) were seeded with hACs and GFP hACs respectively (or *vice versa*). Immature constructs (represented as shaded in the schematic) were cultured statically for 14 days before application of the loading regimen indicated in the panel for further 7 days. Static devices were used as controls. At day 21 constructs were digested, hACs were sorted based on GFP expression, and cells from top and bottom compartment analysed separately though RT-qPCR. **h,** Mechanical compression elicits a different response in cartilaginous constructs in top and bottom compartments as assessed evaluating compartments gene expression and IL8 release. Gene expression was quantified through RT-qPCR (n≥9 biologically independent samples from n=3 donors, for FOS, PTGS2 and C-JUN, n=6 biologically independent samples from n=2 donors for MMP13 and IL8). When comparing cells from top and bottom constructs, statistical significance was determined by paired t-test for normal populations and by Wilcoxon test for non-gaussian populations ((adjusted) p-values < 0.05 are indicated in red). When comparisons were performed with respect to static controls (whose expression levels are indicated by the horizontal lines in black) statistical significance was determined by Kruskal-Wallis test with Dunn’s test for multiple comparisons ((adjusted) p-values < 0.05 are indicated in black). All genes expression was referred to GAPDH expression and values were normalized for the expression of the sorted static controls for each donor. Values are reported as mean + s.d. Gene expression levels referring to top and bottom constructs coming from the same chamber are connected by black lines. IL8 release was determined by ELISA (n=6 biologically independent samples from n=2 donors, Values are reported as mean ± s.d.). Statistics by Kruskal-Wallis test with Dunn’s multiple comparison test. For all graphs, populations’ normality was assumed if both Shapiro-Wilk and Kolmogorov-Smirnov tests resulted positive.

PDMS, the material of the device elements i.e. pillars and walls, was described as an hyperelastic solid with a Mooney-Rivlin strain energy function^11,26^.

Throughout the study, both top and bottom compartments were seeded with cells encased in an enzymatically cross-linkable and degradable 2% w/v PEG-based hydrogel^24^ (which we used to establish our previously published CoC model^11^). Both hydrogels and cartilage exhibit a largely non-linear and strain rate dependent behaviour ^27,28^. A Biphasic PoroElastic (BPE) constitutive relation, which accurately predicts the strain field of these materials^28,29^, was thus adopted to model both PEG hydrogels and cartilaginous constructs.

Preliminary numerical simulations (Supplementary Fig. 5 a-e) were performed considering both top and bottom compartments as simply filled with only the PEG based hydrogel. Subsequent computations were executed using the actual mechanical properties of mature cartilaginous constructs obtained after 14 days of hACs chondrogenic differentiation in the device (Fig. 2b, c).

Poisson’s ratio and specific permeability, assumed respectively equal to 0.33^30^ and 3×10^-4^ mm s^-127^ in all cases, were derived from literature data. The Young’s modulus (E) of the 2% w/v PEG gel (1.9 ± 0.53 kPa) and that of mature cartilaginous constructs (3.66 ± 4.16 kPa) were estimated through Indentation-Type Atomic Force Microscopy (IT-AFM) (Fig. 2b,c). After 14 days of static culture, beside a higher average E value, mature cartilaginous tissues exhibited a wider E distribution (Fig. 2b) with areas characterized by different mechanical properties two representative regions are reported in Fig. 2c). These observations indicated the presence of hACs secreted extra-cellular matrix.

Different plane views of the adopted ROI geometry and mesh are depicted in Supplementary Fig. 5a. The overall strain field was evaluated through Nominal strains (NE) along the principal directions X (i.e. NE11), Y (i.e. NE22), and Z (i.e. NE33). A mesh sensitivity analysis was conducted to ensure the consistency of the results (Supplementary Fig.5b).

Initial simulations (Supplementary Fig. 5c-e) were aimed, given the introduction for the top compartment of three different geometries (i.e. NDG, LDG, and DG), at evaluating the strain field in all cases (both top and bottom compartments were considered as filled with empty PEG gels, E=1.9 kPa).

In all three cases, NE33 values computed for bottom and top constructs were respectively -25% and -2.5% on average, demonstrating the achievement of a consistent and compartment specific mechanical stimuli. Moreover, all geometries exhibited very limited longitudinal and lateral expansion in response to the imposed compression with NE11 and NE22 resulting negligible in both top and bottom constructs and in all geometric configurations. Lateral expansion peaks appeared exclusively in the narrow areas between the T shaped pillars in the bottom compartment and between the pillars of the top compartment in the DG configuration.

Notably, limiting the lateral expansion (i.e. achieving confined compression) allows a finer control over the applied strain field but is also associated with cartilage interstitial fluid outflow, which was linked to chondrocytes mechanotransduction activation due to transient hypo-osmotic stresses secondary to mechanical loading^31^.

Subsequently, a conceptual experiment was performed (using the NDG configuration) to further characterize the device. The bottom compartment was considered as filled with the 2% PEG gel (i.e. E=1.9 kPa), while the top compartment with constructs with a modulus of 1, 2, or 10 times that of the 2% PEG gel (Fig. 2d). Interestingly, limited variations in the strain field of the two constructs were detected, indicating the robustness of the device with respect to possible variations in the mechanical properties of hosted hydrogels/constructs. Finally, a more refined FE-based assessment of the strain field of mature cartilaginous constructs (E= 3.66 kPa) was implemented. In this case we adopted the LDG configuration (which was then employed in the experimental validation of these results). As visible from NE11, NE22, and NE33 contour plots in Fig. 2e, computed strains were homogeneous and close to null in the X and Y directions, while well-defined and compartment specific in the Z direction. The average NE33 resulted equal to -2.9% ± 1.6% and -25% ± 7% in top and bottom constructs respectively (Fig. 2f).

*In vivo*, cartilage strains depend on the performed activity^32^, its duration, the location within the body^33^, and the specific cartilaginous layer depth. Cartilage compression levels were indeed demonstrated to vary deeply within tissue layers. An overall compression of -5% results in strain levels of -35% in the superficial zone, -5% in the transitional zone, and -1% in the deep zone/calcified cartilage^23^. Moreover, strain across the OCU are influenced by the fact that the estimated macroscopic Young’s moduli of cartilage and subchondral bone differ of almost four orders of magnitude (0.8-1.3 MPa for cartilage^16,34^, 11-14 GPa for subchondral bone ^14^) with calcified cartilage functioning as a transition zone ^15^. These values are also affected by OA^14^

*In vitro*, at the microscale, devices are often constituted by PDMS, which is characterized by E values of 1.32-2.97 MPa^35^, in which cellular tissues are obtained by the incorporation of cells in soft hydrogels, thus not permitting the achievement of bone-like E values (i.e. GPa).

Strain values obtained with FE simulations indicated that the present device can circumvent these limitations. Estimated compression levels are well in accordance with desired ideal targets, reaching values of cartilage HPC, which was correlated to the onset of OA both *in vivo*^7,36,37^ and in our previous COC model^11^, while maintaining the low strain experienced by subchondral tissues^23,32^.

Notably, cartilage and subchondral bone *in vivo* are subjected to a wide variety of stimuli comprehensive of compression but also shear stresses^38^. The full recapitulation of these stimuli in such a bi-compartmental model is highly complex and would introduce multiple confounding factors. To better control the experimental environment trying to dissect the still unclear mechanisms by which, for instance, chondrocytes respond to loading, we focused exclusively on confined compression.

### Providing 3D microconstructs with compartment specific compression levels: experimental validation

We next experimentally validated numerical results proving that cartilaginous constructs hosted in top and bottom compartments exhibit different cyclic loading responses.

To this aim, GFP hACs and non-labelled hACs were embedded in 2% w/v PEG hydrogels and respectively seeded in bottom and top compartments. Cells were then cultured statically for 14 days and subsequently exposed to cyclic mechanical compression. A schematic of the experimental timeline is reported in Fig. 2g.

Notably, after 14 days of static maturation constructs assumed a hyaline cartilage-like gene expression signature (i.e. the increased expression of collagen type II to collagen type I ratio (*COL2A1/COL1A1*), aggrecan (*ACAN*), and lubricin or proteoglycan 4 (*PRG4*), but also of the wingless-type MMTV integration site (Wnt) antagonist Frizzled-related protein (*FRZB*) and Bone Morphogenetic Protein (BMP) antagonist Gremlin-1 (*GREM1*), recently characterised as hypertrophy brakes^39,40^, and a positive trend in the expression of the cartilage development associated growth differentiation factor 5 (*GDF5*)^41^, Supplementary Fig. 6a). We previously demonstrated these characteristics to be necessary to obtain clinically relevant load responses in our CoC model^11^. Here we verified their replication in the bi-compartmental device proving, furthermore, that chondrocytes formed compact cartilaginous tissues in both device compartment, but maintained well defined bi-layer structures (Supplementary Fig. 6b, c).

At the end of the chondrogenic culture, hACs based constructs were enzymatically digested and sorted by flow cytometry discriminating on GFP positivity. The purity of the populations after sorting was verified checking GFP expression levels by RT-qPCR and through direct observation of GFP expression in re-plated sorted populations (Supplementary Fig. 7).

The two hACs populations were separately analysed to quantify the expression of Fos Proto Oncogene (*FOS*), Prostaglandin-endoperoxide synthase 2 (*PTGS2*), and Jun proto-oncogene, AP-1 transcription factor subunit (*C-JUN*), i.e. a set of genes associated with mechanotransduction signalling via Transient Receptor Potential Cation Channel Subfamily V Member 4 (TRPV4)^31^. Additionally, we analysed the expression of matrix metalloproteinase 13 (*MMP13*), and of the pro-inflammatory cytokine interleukin 8 (*IL8*), which are both highly expressed in OA cartilage and associated with cartilage breakdown^42–44^.

Notably, the expression of none of these genes differed in top and bottom compartments of static controls (Supplementary Fig. 8a) excluding a possible role of the geometry of these structures in modulating the expression profile of hosted cells.

Following cyclic compression, a statistically significant upregulation of *FOS*, whose inhibition has been correlated to cartilage protection in mice models of OA^45,46^, was registered for hACs isolated from both compartments with respect to static controls (*p*-values in black in Fig. 2h). This result suggests that constructs in both compartments are affected by the cyclic loading. Additionally, the expression of the transcription factor *C-JUN*, which is involved in joint cells fate specification^47^, and that of the OA-related collagen degrading enzyme *MMP13*^42^, were significantly increased in hACs that were cultured in the bottom compartment with respect to those cultured in the top one (in pairwise analyses of samples from the same device, *p*-values in red). These selective modulations indicate the achievement of differential compression levels and associated distinctive mechanotransduction-mediated cellular responses within the bi-layered constructs. The expression of *PTGS2* and *IL8* did not show differences between compartments and were also not modulated (*PTGS2*), or were downregulated (*IL8*) with respect to static controls. This is probably due to constructs digestion and sorting (static controls were also subjected to these stress inducing steps before comparisons with other groups), which caused an overall increase of *FOS*, *PTGS2*, *C-JUN*, and *IL8* expression (Supplementary Fig. 8b), thus possibly masking some gene modulations due to cyclic compression.

The onset of an inflammatory phenotype was thus assessed measuring the release of the IL8 protein in the culture medium. When HPC was applied, a statistically significant higher concentration of this inflammatory cytokine was measured only in the culture medium collected from the bottom compartment (Fig. 2h). Notably, the different trends in the cytokine’s concentration could be measured given that the LDG limits solutes diffusion between compartments.

In summary, reported experimental results and FE estimates constitute a proof of concept that the proposed device provides superimposed constructs with discrete tissue-specific compression levels consistent with those experienced by OCU tissues *in vivo*. Additional investigations will nonetheless be necessary to show a correlation between applied compression levels and a quantitative modulation in the expression of reported genes, in particular *FOS*, *PTGS2*, and *C-JUN* whose modulation following mechanical stimulation was only obtained with the application of a ∼10% compression ^31^. Another aspect to consider is that different mechanotransduction mechanisms e.g. the activation of PIEZO1 and PIEZO2 channels, were also reported to be affected by over-physiological compression levels ^48,49^. Ulterior differences might hence be detected investigating alternative pathways.

### Subchondral layer’s mechanical properties affect hACs response to loading

OA has been associated with changes in the structure^5^, the mechanical properties^50^, and the entity of mineralization of calcified cartilage and subchondral bone^1,51^. Hyper-mineralized calcified cartilage was found in hips from OA patients^51^, possibly altering loading patterns and accelerating cartilage abrasion. Localized stiffening of areas immediately adjacent to the tidemark were correlated to increased stresses in cartilage deep layers ^52,53^. Changes in subchondral bone architecture were demonstrated to alter stress distribution in articular cartilage leading to transforming growth factor beta (TGF𝛽) activation, loss of hACs homeostasis, and impairment of the tissue metabolic activity^54^. Whether these subchondral alterations precede and cause cartilage deteriorations remains however to be elucidated.

We therefore performed a further set of experiments to prove that the proposed device could be used to examine, *in vitro*, how alterations in the mechanical properties of subchondral layers affect cartilaginous constructs’ response to loading.

The adopted experimental setup is depicted in Extended Data Fig. 2a. The bottom compartment was seeded with hACs laden PEG hydrogels, while the top compartment (in the NDG configuration) with PEG hydrogels loaded with polystyrene beads (diameter 10 μm) with volumetric fractions of 0.1% (indicated as 0.1% B) or 5% (indicated as 5% B), respectively. Images of the hydrogels in the top compartment are visible in Extended Data Fig. 2b. The stiff plastic beads (that have an E modulus of 3250 MPa^55^) locally alter the hydrogel mechanical properties proportionally to their volumetric fraction as assessed by IT-AFM (Extended Data Fig. 2c, d).

Construct were cultured statically for 14 days to achieve mature cartilaginous tissues and subsequently subjected to cyclic loading for 7 days.

Cartilaginous constructs response to loading depended on the local E modulus of the hydrogel in the top compartment. A higher volumetric fraction of beads resulted in hACs showing a higher gene expression of the degradative enzyme *MMP13* (in a non-statistically significant way) and of the inflammatory cytokine *IL8* (in a statistically significant manner, both with respect to static controls and comparing the 0.1% B and the 5% B substrates), as shown in Extended Data Fig. 2e. Similarly, hyperphysiological loading induced and increase in the release of proMMP13, IL8, and IL6, with the concentration of IL8 and IL6 being significantly higher for the 5% B substrate (Extended Data Fig. 2f), while interleukin 1 beta (IL1 𝛽) and tumour necrosis factor alpha (TNF𝛼) were not detectable (data not shown).

These outcomes indicate that the introduced device can effectively be used to study how the mechanical properties of subchondral layers affect hACs load responses in a highly controlled environment. Our data also suggest that even subtle changes in subchondral bone properties or architecture, beside playing a role in late OA stages as previously demonstrated^54^, might trigger, or at least exacerbate, early degenerative events in the cartilage layer.

Of note, the usage of a completely artificial (i.e. cell-free) substrate, as our polystyrene loaded PEG hydrogels, limits the implications of these results in the assessment of the cellular crosstalk between cartilage and subchondral bone (e.g. we cannot investigate what caused subchondral mechanical alterations in the first place). Further investigations on these regards have thus been conducted in the following paragraphs.

### Establishment of a subchondral mineralised tissue on-chip (SMToC)

For a complete OCU on-chip model we required two components: the above introduced and validated technological platform, and a cell-based biphasic construct with a cartilaginous like-layer and a subchondral bone-like layer.

We formerly demonstrated the capacity of hACs to differentiate into hyaline-like microconstructs in our CoC model^11^.

Here we set to establish a 3D single culture model of subchondral (bone-like) mineralized tissue on-chip (SMToC). To this end, we made use of human bone marrow derived mesenchymal stromal cells’ (bmMSCs) intrinsic commitment to hypertrophic differentiation^56^. Indeed, these cells are able to undergo a program recapitulating embryonic endochondral ossification *in vitro* with the eventual formation of bone *in vivo*^57^.

As a first step, we adopted our CoC^11^ platform which is intended for the culture of single constructs only. The newly introduced SMToC was however meant to be coupled with our hyaline cartilage model. Given this aim, we optimized an osteochondral culture medium formulation (OCM) that could drive the formation of mineralized subchondral tissues starting from bmMSCs, while not affecting hACs chondrogenic differentiation.

bmMSCs isolated from healthy donors were embedded in the PEG-based hydrogel formulation described earlier and injected in microfluidic devices. Cells were cultured statically for 14 days in either a serum free chondrogenic medium^58^ (CHM, see methods section), or in OCM (Fig. 3a). This latter medium was derived from CHM with the addition of beta-glycerophosphate (𝛽-GP) which serves as a source of inorganic phosphate when hydrolysed by alkaline phosphatase (ALP), and was previously reported to be involved in hydroxyapatite(HA) deposition and bone formation^59^.

**Fig. 3.**
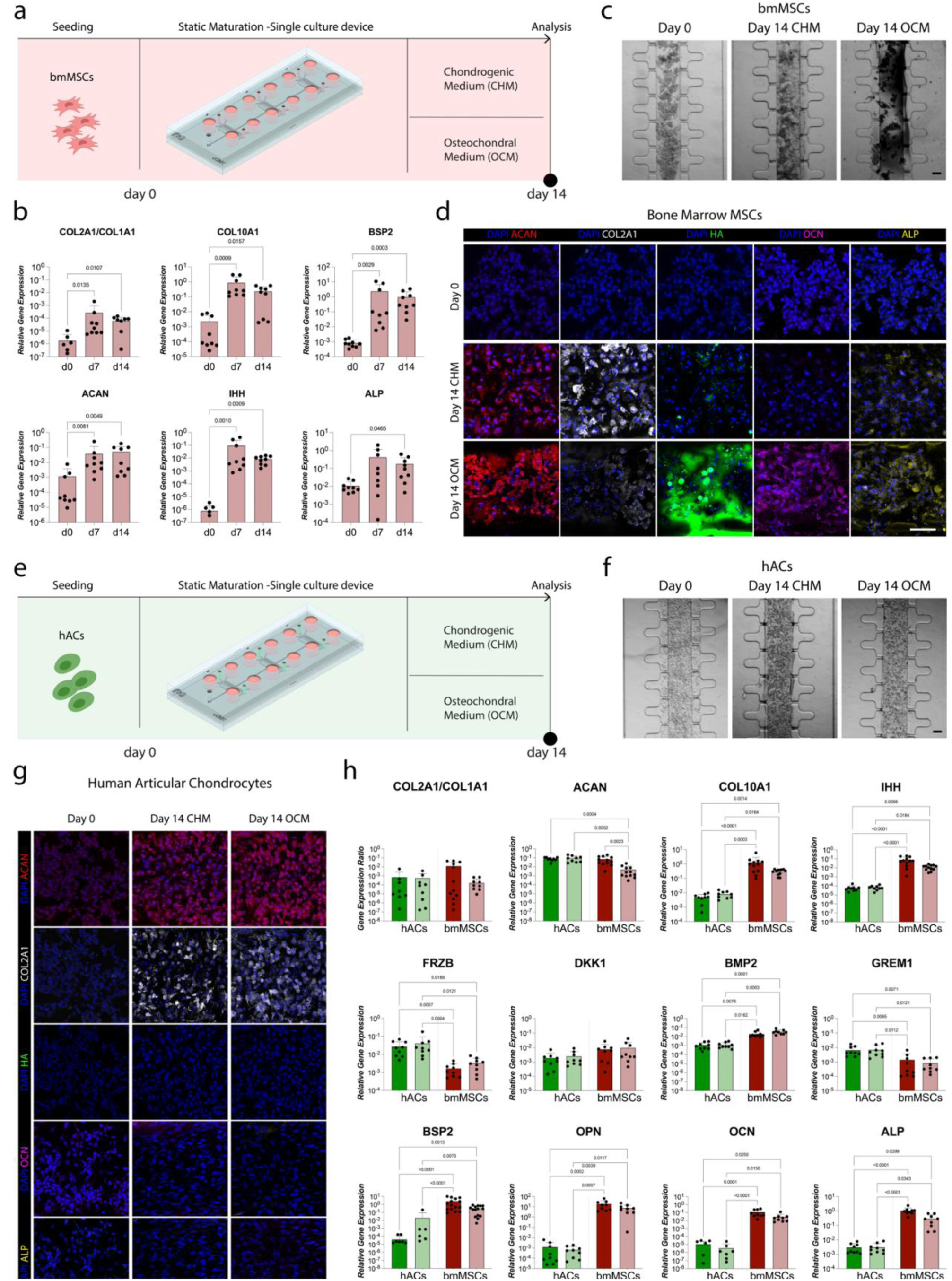
Achievement of subchondral-like mineralized tissues on-chip. **a**, Schematic indicating the experimental procedure used for bmMSCs differentiation. bmMSCs were seeded our previously introduced CoC device^11^ and cultured statically for 14 days in either in CHM or OCM **b**, SMToCs mature over time. Gene expression was quantified through RT-qPCR (n=9 biologically independent samples from n=3 donors were analysed for each condition and each cellular type). Statistical significance was determined by Kruskal-Wallis test with Dunn’s post hoc test for multiple comparisons. All genes expression was referred to GAPDH expression. Values are reported as mean + s.d. **c**, Brightfield pictures of bmMSCs constructs at day 0 and after 14 days of static maturation. Construct appeared black upon exposure to OCM, consistently with the deposition of calcium salts. (n≥30 biologically independent samples from n≥5 donors for each cellular type were considered in analyses). Scale bar, 100 μm. **d**, Immunofluorescence images bmMSCs constructs cultured in CHM or OCM for 14 days. Day 0 constructs were adopted as negative controls (n≥.1 biologically independent samples from n≥3 donors were considered in analyses). The OCM consistently led to bmMSCs depositing hydroxyapatite (HA). Hyaline cartilage markers ACAN and COL2A1 are represented in red and white respectively. HA is represented in green. The bone markers ALP and OCN are represented in in yellow and magenta. In all images DAPI (represented in blue) was used as nuclear counterstaining. Scale bar, 100 µm. **e**, Schematic indicating the experimental procedure used to assess if the OCM impaired hACs chondrogenic differentiation capability. hACs were seeded in our previously introduced CoC device^11^and cultured statically for 14 days in either CHM or OCM. **f**, Brightfield pictures of hACs constructs at day 0 and after 14 days of static maturation. Constructs appeared similar after culture in CHM and OCM. Scale bar, 100 μm. **g**, Immunofluorescence images hACs constructs cultured in CHM or OCM for 14 days. Day 0 constructs were adopted as negative controls (n≥.1 biologically independent samples from n≥3 donors were considered in analyses). The OCM formulation does not alter hACs chondrogenic potential. Hyaline cartilage markers ACAN and COL2A1 are represented in red and white respectively. HA is represented in green. The bone markers ALP and OCN are represented in in yellow and magenta. In all images DAPI (represented in blue) was used as nuclear counterstaining. Scale bar, 100 µm. **h**, Assessment of medium formulation and cell type effects on chondrogenic and osteogenic gene expression signature. Gene expression was quantified through RT-qPCR. (n≥9 biologically independent samples from n≥3 donors were analysed for each condition and each cellular type). Statistical significance was determined by Kruskal-Wallis test with Dunn’s post hoc test for multiple comparisons. All genes expression was referred to GAPDH expression. Values are reported as mean + s.d. Passing from HCM to OCM did not imply alteration in hACs and bmMSCs gene expression which retained distinct and defined profiles. For all graphs, populations’ normality was assumed if both Shapiro-Wilk and Kolmogorov-Smirnov tests resulted positive. (Adjusted) P values < 0.05 are reported on the graph.

After 14 days of static maturation in OCM, bmMSCs exhibited a gene expression signature indicative of endochondral ossification (Fig. 3b). Constructs had an increased expression of markers of chondro-differentiation (i.e. *COL2A1/COL1A1* ratio and *ACAN*), of assumption of hypertrophic traits (i.e. collagen type X (COL10A1)^60,61^ and Indian hedgehog (IHH)^62^), and of mineralization processes (i.e. *ALP* and bone sialoprotein 2 (*BSP2*^63,64^)).

These results were confirmed at the ECM level: tissues generated in both CHM and OCM were positive for COL2A1 and ACAN, but only the latter ones showed dark deposits in brightfield images (Fig. 3c, d) corresponding to areas positive for HA (Fig. 3d), and produced the bone-specific protein osteocalcin (OCN^65,66^). ALP was accumulated at low levels in CHM cultured bmMSCs (in accordance to the intrinsic capacity of these cells to acquire (pre)hypertrophic traits in response to pro-chondrogenic factors ^56^) and at high levels in bmMSCs cultured in OCM (Fig. 3d).

To exclude a possible negative effect of 𝛽-GP on hACs chondrogenesis, the same experiments were repeated with hACs comparing the extent of chondrocytes’ differentiation capacity in CHM and OCM (Fig. 3e).

hACs based constructs cultured in CHM and OCM exhibited similar appearances in brightfield images (Fig. 3f). Moreover, regardless of the adopted medium formulation, these cells formed constructs positive for COL2A1 and ACAN, and negative for ALP, OCN and HA (Fig. 3g). While with both media the cartilage-forming capacity by hACs was highly donor dependent in accordance with previous literature^67^, (Supplementary Fig. 9), these results indicated that the addition of 𝛽-GP does not impair the chondrogenic potential of these cells. We then analysed a more comprehensive set of genes assessing the effect of OCM (versus CHM) on both hACs and bmMSCs(Fig. 3h).

In hACs, the OCM (with respect to the standard chondrogenic medium), did not induce changes in none of the investigated chondrogenic markers (i.e. *COL2A1/COL1A* ratio and *ACAN*); of hypertrophy markers (*COL10A1* and *IHH*); and of BMP and Wnt pathways related markers (i.e. *FRZB*^3968^, dikkopf 1 homologue (*DKK1*^69^), *BMP2*^70,71^, and *GREM1*^68^).

Instead, in referral to bmMSCs, switching from CHM to OCM caused a decrease in the *COL2A1/COL1A1* ratio (non-statistically significant) and of the expression of *ACAN* (statistically significant), results that were in accordance with differentiation towards a mineralized tissue.

Clear statistically significant differences in gene expression were registered between cell types. hACs and bmMSCs maintained well distinct cellular phenotypes correlating with embryonal development where hyaline and transient cartilage are selectively exposed to Wnt and BMP signalling^72^. Specifically, hACs exhibited significantly higher levels of the Wnt and BMP antagonists *FRZB* and *GREM1*, whose decreased expression was correlated with the onset of OA^39,68^, and lower levels of *BMP2* indicating that the OCM does not induce hypertrophic differentiation.

The expressions of the bone related markers *ALP*^73^, osteopontin (*OPN*^74,75^), and bone sialoprotein 2 (*BSP2*^63,64^), was consistently higher in bmMSCs with respect to hACs, while it was not affected by OCM exposure.

Overall, these results indicate that, with our OCM formulation, we could obtain, respectively starting from hACs and bmMSCs, the formation hyaline-like tissues and of mineralized constructs. The adopted medium composition seemed therefore optimal for subsequent steps towards the achievement of a complete OCU-on-chip model.

To increase the robustness of above described results the effect of adding 𝛽-GP to the CHM formulation was also evaluated in standard 3D cellular aggregates (i.e. pellets) culture.

The OCM formulation did not alter hACs phenotype. hACs pellets cultured in OCM contained similar quantities of glycosaminoglycans (GAG) to those cultured in CHM, remained negative for alizarin red staining of calcium deposit and for COL10A1 and, in a highly donor dependent way, did not alter their COL2A1 deposition (Extended Data Fig. 3a). The overall chondrogenic quality of the pellets was also unchanged as determined through an automated evaluation of the modified Bern score^76^ (Extended Data Fig. 3b).

Analogously to what observed on-chip, instead, the addition of 𝛽-GP induced bmMSCs to mineralize, despite not altering their modified Bern score (Extended Data Fig. 3c, d). Notably, bmMSCs-derived pellets deposit COL2A1 but are markedly positive for COL10A1 even in CHM, confirming the previously reported^56^ commitment of these cells to the acquisition of a hypertrophic phenotype.

Similar conclusions were also obtained considering GAG/DNA quantifications and gene expression analyses (Extended Data Fig. 3 e-h).

We therefore demonstrated that a mineralized subchondral tissue can be obtained on-chip through our OCM formulation and that said formulation does not induce mineralization or hypertrophy of hACs. Confirming these results with a different model, moreover, reinforces the rational to exclude that possible effects observed co-culturing hACs and bmMSCs are not medium-dependent and can be ascribed to a crosstalk between the cellular populations.

### Establishment of a biphasic osteochondral unit (OCU)-on-chip

Next, merging the above-mentioned 3D layers, we aimed at obtaining a mature osteochondral-like construct, i.e. an OCU-on-chip model with directly interfaced hyaline-like cartilage and mineralized subchondral tissues. The VBV-featuring platform that we introduced (Fig. 1a) allows to provide biphasic constructs with patho-physiologically relevant tissue specific mechanical stimuli. The vertically stacked tissue disposition is however suboptimal for direct microscopy-based observation of the two constructs. To assess the formation of osteochondral-like constructs starting from hACs and bmMSCs a different device where the two tissues are co-cultured with a side-by-side horizontal disposition was introduced.

The device was based on our CoC^11^ model having two rows of suspended T-shaped pillars that separate the cellular constructs from culture medium channels. In this case, however, there are two hydrogel compartments divided by a row of hexagonal pillars designed to maximize the interface are between the constructs. Fig. 4a depicts the device and the experimental setup.

**Fig. 4.**
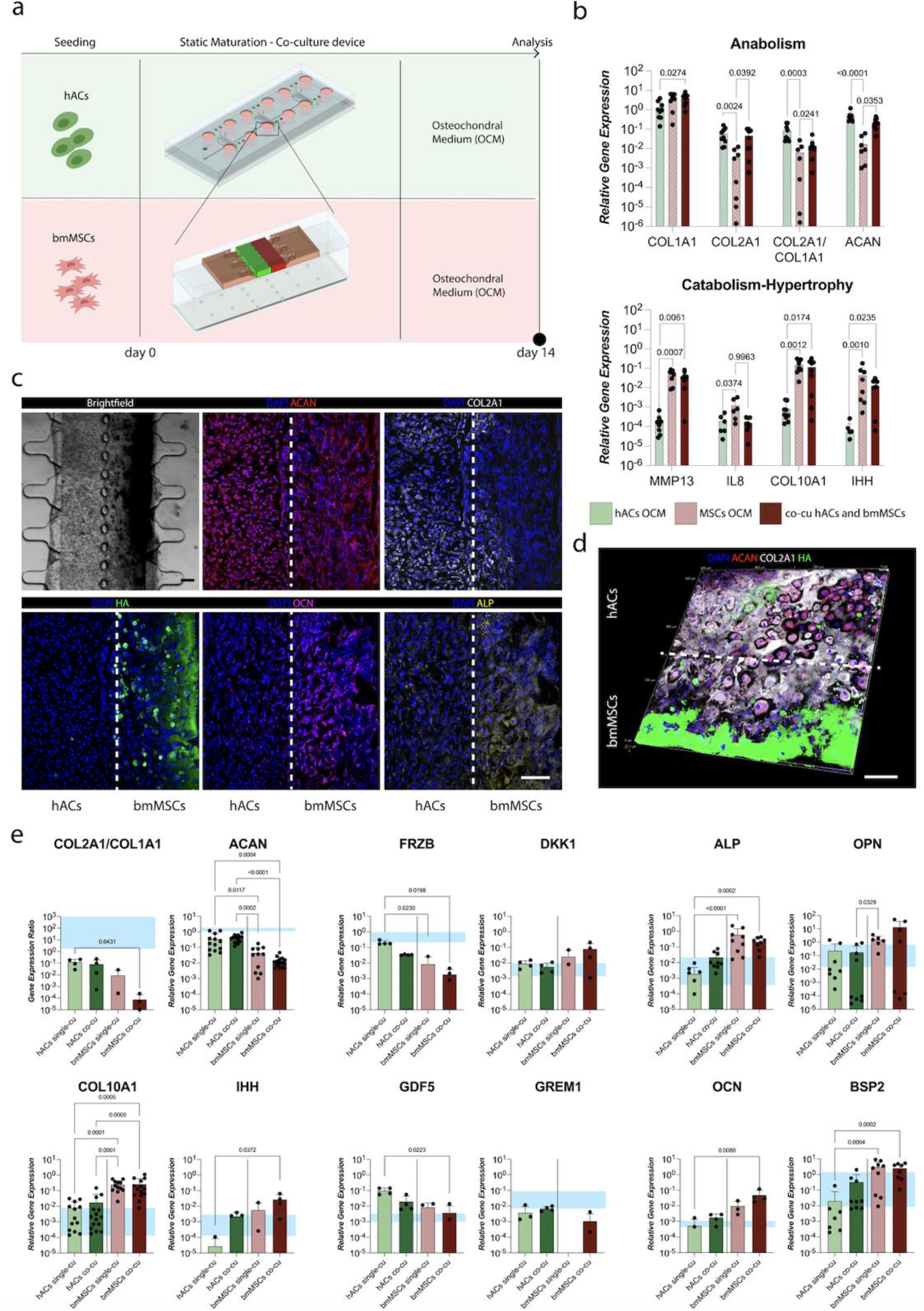
Achievement of bi-phasic osteochondral constructs on-chip. **a**, Schematic of the experimental procedure. hACs and bmMSCs were co-cultured statically for 14 days in OCM. A device capable of hosting two 3D constructs side by side was introduced for the purpose. The inner geometry of the device inspired to our published COC platform is represented in the inset. The hACs construct is represented in green, the bmMSCs one in red. Constructs are separated by suspended hexagonal shaped pillars designed to maximise the interface area between them. Such a horizontal disposition does not allow to provide a construct dependent compression level but it permits an easier visualization of both tissues and of their interface and was therefore adopted in the maturation phase. **b**, Gene expression of hACs and bmMSCs in single culture and of co-cultures after 14 days of static maturation in OCM. The co-cu expression refers to the two constructs analysed together; values follow within the range delimited from the expression of hACs and MCs in single culture. Gene expression was quantified through RT-qPCR (n=3 biologically independent samples from n=3 donors/experiments). Statistical significance was determined by Kruskal-Wallis test with Dunn’s post hoc test for multiple comparisons. Populations normality was assessed though Shapiro-Wilk and Kolmogorov-Smirnov tests. (adjusted) P < 0.05 are reported on the graph. All genes expression was referred to GAPDH expression. Values are reported as mean + s.d. **c**, **d** Immunofluorescence images of osteochondral constructs after 14 days of static maturation. hACs are located on the left, bmMSCs on the right (n=2 biologically independent samples from n≥3 donors/experiments were considered in analyses). Hyaline cartilage markers ACAN and COL2A1 are represented in red and white respectively. HA is represented in green. The bone marker ALP and OCN are represented in yellow and magenta respectively. In all images DAPI (represented in blue) was used as nuclear counterstaining. Scale bar, 100 µm. The hexagonal pillar line separating the constructs is represented by the dotted line. Both constructs were positive for cartilage markers (the staining positivity depending on the single donor, Supplementary Fig. 11), while HA, ALP and OC were largely confined to the subchondral construct. Differently to hACs cultured in single culture, HA positive cells appeared also in the hACs compartment. Overall biphasic osteochondral constructs could be achieved consistently. Moreover, as highlighted in the 3D confocal reconstruction in **d,** constructs tended to be polarized with abundant HA deposition at the border and a decreasing gradient approaching the hACs compartment. **e**, Assessment of hACs and bmMSCs gene expression in co-culture. Osteochondral construct composed of GFP hACs and bmMSCs were obtained after 14 days of static maturation. Constructs were digested enzymatically and cell populations sorted to be analysed separately. Derived hACs and bmMSCs gene expression was compared to that of the same cells cultured in single culture. Results were also compared to the gene expression of human knee cartilage samples whose range of expression (with values indicati^39^ng the expression from cartilage superficial zone to cartilage deep zone) is reported by the cyan band on the graphs. hACs and bmMSCs gene expression remained well differentiated originating distinct tissues. The expression of some bone markers in hACs seems to increase (in a non-statistically significant manner) as a consequence of the co-culture but it remains within physiological levels. Differentiated bmMSCs express levels of bone tissue markers and hypertrophy markers which are higher than those of human knee cartilage at the interface with subchondral bone. Gene expression was quantified through RT-qPCR (n=4 biologically independent samples from n=2 bmMSCs donor and n=1 hACs donor i.e. GFP positive hACs). Statistical significance was determined by Kruskal-Wallis test with Dunn’s post hoc test for multiple comparisons. Populations normality was assessed though Shapiro-Wilk and Kolmogorov-Smirnov tests. (adjusted) P < 0.05 are reported on the graph. All genes expression was referred to GAPDH expression. Values are reported as mean + s.d.

A first characterization was performed assessing the gene expression of whole OCUs-on-chip and comparing it to that of single culture controls (Fig. 4b). Notably, we confirmed that the gene expression of OCU-on-chip constructs is not affected by the tissues disposition (i.e. horizontal or vertical, Supplementary Fig. 10), thus supporting the use of a simplified device configuration.

OCU constructs expressed *COL2A1/COL1A1* and *ACAN* levels similar to those of cartilaginous tissues and significantly higher ones than those of mineralized counterparts. They also had *COL10A1* and *IHH* expression similar to those of the SMToC constructs alone. These data simply reflect that OCU constructs are composed of both cell types. Interestingly, OCUs and bmMSCs showed high *MMP13* levels, but while bmMSCs also had higher *IL8* expression this was not the case for OCU constructs. This might indicate co-culture dependent non inflammatory remodelling processes.

We then made use of the side-by-side disposition obtaining a direct observation of both tissues and of their interface. Clear differences between hACs-derived and bmMSCs-derived layers were evident from brightfield images (Fig. 4c, Supplementary Fig. 11): dark mineralized areas appeared exclusively in the bmMSCs compartment. Immunofluorescence images confirmed then the formation of osteochondral-like constructs with a cartilaginous layer (derived from hACs) positive for ACAN (whose positivity remained also in the bmMSCs layer) and COL2A1, and a mineralized layer (derived from bmMSCs) positive for HA, ALP, and OCN (Fig. 4c). A certain donor to donor variability could be seen in the quality of OCU constructs obtained in the device, but this was in line with the characteristic donor-dependency of hACs and bmMSCs (Supplementary Fig 11).

Notably, while hACs in single culture remained completely negative for the HA staining (Fig. 3d, Supplementary Fig. 9), in co-cultures HA positive spots were also detectable in the cartilaginous layer (Fig. 4c, d, Supplementary Fig. 11).

This observation could be explained by an active cross-talk of the two populations that induces hACs to transdifferentiated, or by migration of cells from ones to the other compartments. Cells expressing HA in the cartilage layer could be bmMSCs coming from the bone layer.

Further evidence of a mutual influence between the two compartments was given by mineralization patterns in bmMSCs-based constructs. These were homogeneous and symmetrical in single culture (Fig. 3d, Supplementary Fig. 9), but a gradient appeared in co-cultures with areas strongly positive for HA being limited to the construct’s region further away from the hACs layer (Fig. 4d, Supplementary Fig. 11).

Analogous results were obtained using the VBV featuring device where constructs were sectioned vertically to properly observe the interface (Extended Data Fig. 4 a, b).

These data collectively indicate that we successfully obtained OCU-like tissues on-chip and point towards a possible two-way cross-talk between hACs and bmMSCs. A deeper understanding of these interplay seems to require however a distinction between cellular subpopulations.

To further dissect possible effects of the co-culture, the osteochondral differentiation experiment was repeated using GFP positive chondrocytes so that, at the end of the culture, constructs could be digested and GFP positive hACs separated from bmMSCs. Populations were analysed separately and compared with single culture counterparts. Both OCU constructs with a horizontal and a vertical disposition were considered (Fig 4e, Extended Data Fig. 4c).

Constructs gene expression levels were also compared to those of samples of human knee cartilage (represented by the cyan bands on the graph, Fig. 4e) obtained from patients undergoing knee replacement. The whole range of gene expression levels in cartilage was determined comparing the expression of cartilage in superficial and deep zones (Supplementary Fig. 12).

In hACs, the co-culture led to upward trends in the expression of the hypertrophic marker *IHH* and of the bone markers *ALP*, *OCN*, and *BSP2*. All hACs gene expressions remained however within at physiological ranges (with the exception of the *COL2A1/COL1A1* ratio which was still lower than values of the joint specimens). bmMSCs-based mineralized constructs expressed higher levels of hypertrophy and bone markers than those of cartilage deep zones thus indicating the achievement of tissues more resembling of subchondral calcified cartilage and bone.

Despite differences observed through immunofluorescence images, no statistically significant changes in gene expression were detected between co-cultured hACs and bmMSCs and single culture counterparts. Pronounced differences remained instead between the two cell types.

These findings collectively indicate that mature osteochondral-like constructs with clear layers separation could be obtained on-chip but also that further investigations are warranted.

One thing to consider is that RT-qPCR analyses are highly influenced by abundant cells. To better understand differences between single and co-cultures, such as the nature of HA positive cell within cartilaginous constructs, more refined methodologies (as those described in subsequent sections) are necessary.

### Cyclic hyperphysiological loading of OCU-on-chip constructs drives cartilage anabolism and subchondral tissue mineralization

OA is characterized by changes in the whole OCU which receiving and dissipating the stress associated with movement and loading is continuously biomechanically challenged ^1,77^. Different studies point to a complex interplay between the two tissues upon mechanical stimulation. Subchondral bone structural abnormalities have been correlated to increased peak stresses in articular cartilage resulting in altered TGFβ signalling and disruption of chondrocytes metabolism^54^. At the same time, abnormal mechanical loading due to an anterior cruciate ligament transection (ACLT) was demonstrated to activate TGFβ1 in subchondral bone in a mouse model of OA^78^.

We set to discriminate which traits of the OA phenotype are elicited in OCU tissues by hyperphysiological compartment-specific loading.

Firstly, we assessed if hyperphysiological stimulation had an effect on the deposition of calcium phosphate crystals. Changes in subchondral bone mineral content have been reported in various (early) OA animal models^79^ and in OA patients^1,74^. Moreover, OA was associated with chondrocalcinosis, that is to say calcification of hyaline cartilage^74^, and hyper-calcification of calcified cartilage at the interface between cartilage and bone^51^. Some authors even posited calcium particles to function as stress concentration points driving (and preceding) cartilage degeneration^4,50,51,53^. A schematic of the experimental setup is depicted in Fig. 5a. Upon mechanical stimulation, we observed a significant increase in constructs’ HA content (Fig. 5b, c). HA deposition was mainly located in the mineralized subchondral tissue, where HA deposits appeared also to be distributed in the extracellular space. The same increase was detected considering calcium salts released in the culture medium and deposited in medium channels and reservoirs (Fig. 5d) consistently with the detection of calcium crystals in the synovial fluid of OA patients^80^. A feature of interest is that, while the major differences in HA deposition were measured in the subchondral layer, this tissue is exposed to an almost null compression level in the device (Fig. 2 e, f); it is the cartilage layer which is subjected to a ∼30% hyperphysiological compression. These data highlight therefore a synergistic response of OCU tissues to aberrant stimuli.

**Fig. 5.**
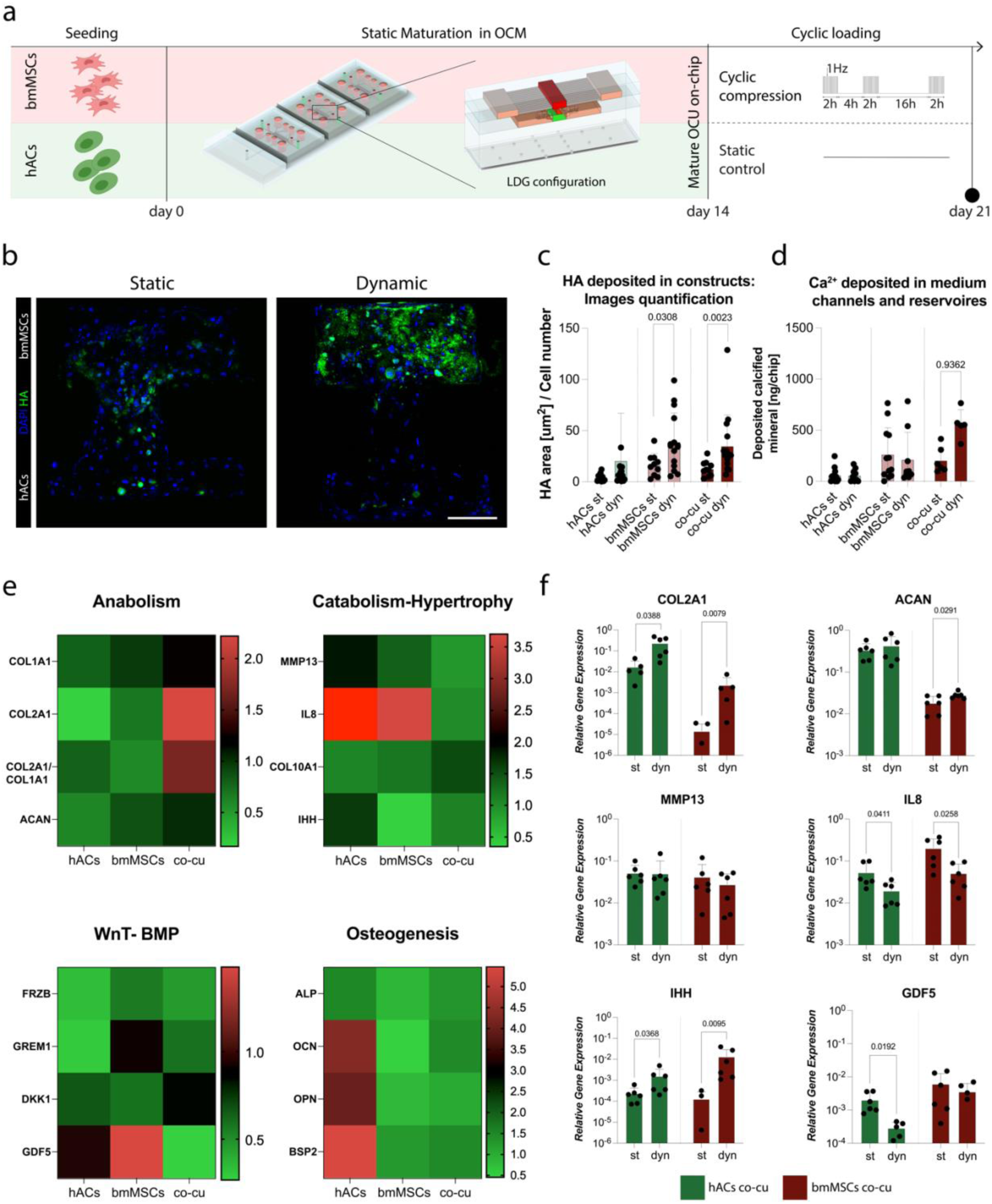
Hyperphysiological cyclical loading elicits early-OA like responses in OCUs-on-chip. **a**, Schematic indicating the experimental plan. hACs and bmMSCs were seeded in VBV-featuring microfluidic devices (with the LDG configuration) and co-cultured statically for 14 days in OCM thus achieving mature OCU-on-chip constructs . Constructs where then exposed to the loading regimen indicated in the panel for the following 7 days. Static constructs were used as controls. The same experiment was performed with hACs and bmMSCs cultured separately (i.e.in single culture), in the CoC device. **b-d**, Cyclic mechanical loading induces an increase in subchondral tissues mineral content. **b**, Immunofluorescence images of representative sections of static (left) and cyclically loaded (right) osteochondral constructs at day 21. hACs based cartilaginous constructs are located at the bottom, bmMSCs based mineralized tissues at the top (n≥10 sections from n=2 hACs and bmMSCs donors/experiments were considered in analyses). HA is represented in green. DAPI, which is represented in blue, was used as nuclear counterstaining. Scale bar, 100 µm. **c**, Quantifications of HA stainings in osteochondral constructs sections. Image analysis was performed using QuPath^89^. HA positive areas were selected using a threshold-based pixel classifier. HA positive areas were normalized for the number of cells in the section. Top areas were considered comprehensive of the tilted-H VBV necking area. Statistical significance between static and cyclically loaded constructs was determined by two tailed unpaired t-test for normal population and by Man-Whitney test for non-gaussian populations. (n≥10 sections from n=2 hACs and bmMSCs donors/experiments were considered in analyses). Values are reported as mean ± s.d. **d**, Quantification of calcium deposits in culture medium channels and reservoirs as determined by alizarin red stainings. Devices were opened to remove the cellular constructs and calcium deposits in the culture medium channels and reservoirs stained with Alizarin Red. Deposits were then dissolved and quantified (n≥5 biologically independent samples from n=2 experiments were considered in analyses). . Statistical significance between static and cyclically loaded samples was determined by two tailed unpaired t-test for normal population and by Man-Whitney test for non-gaussian populations. Values are reported as mean ± s.d. **e**, Comparison of the effect of cyclic mechanical loading on hACs and bmMSCs single cultures, and osteochondral constructs. Single cultures were performed in our previously published COC device; osteochondral constructs were cultured in the VBV device (LDG). Heatmaps represent the gene expression fold change with respect to static controls. In each case the gene expression value was normalized static control expression of the correspondent cell population (n=9 biologically independent samples from n=3 hACs donors and n=3 bmMSCs donors were considered). **f**, Cyclical loading appears to cause an early-OA response in OCU-on-chip constructs’ layers. Osteochondral construct composed of GFP hACs and bmMSCs were obtained after 14 days of static maturation and subjected to cyclical loading for further 7 days. Constructs cultured statically for 21 days were used as controls. To analyse separately the two cell populations constructs were digested enzymatically and cell populations sorted. Gene expression was quantified through RT-qPCR (n=6 biologically independent samples from n=2 bmMSCs donor and n=1 hACs donor i.e. GFP positive hACs). Statistical significance was determined, comparing cells from the same compartments in static and cyclically loaded devices, with two tailed unpaired t-test for normal populations and though Man-Whitney test for non-gaussian populations. All genes expression was referred to GAPDH expression. Values are reported as mean + s.d. In all cases populations normality was assessed though Shapiro-Wilk and Kolmogorov-Smirnov tests. P values ≤ 0.05 are reported on the graphs.

We then better characterised the OCU-on-chip response in terms of gene expression and compared it to that of single cultures of hACs-based cartilaginous construct and of bmMSCs-based mineralized tissues.

All tissues were cultured statically in OCM for 14 days and then subjected to cyclic mechanical loading for 7 days. Mechanical compression was applied with the pattern described in Fig. 5a using our CoC platform for hACs and bmMSCs single cultures and the VBV device for OCU constructs. Dexamethasone, which has an anti-inflammatory and anti-degradative effect^81^ was removed from the OCM during mechanical stimulation. In this phase the gene expression of the OCU-on-chip was analysed as a whole (i.e. without separating cartilaginous and subchondral layers).

Fig. 5e depicts a heatmap of the gene expression fold change of cyclically loaded constructs with respect to model specific static controls (non-normalized gene expression levels are reported in Supplementary Fig. 13). OCU constructs and single culture counterparts exhibited different behaviours.

hACs based cartilaginous construct responded to hyperphysiological loading as previously reported in our CoC model. We detected a decrease in the expression of anabolic markers, such as *COL2A1* and *ACAN*, and the assumption of an inflammatory (marked by *IL8*), degradative (by *MMP13*) and hypertrophic (by *COL10A1* and *IHH*) phenotype (Fig. 5e, Supplementary Fig. 13). They also showed a reduction in the expression of the BMP and Wnt antagonists *FRZB*, *GREM1*, and *DKK1*, similarly to what reported for OA cartilage in patients^39,68^.

We could also observe an increase in the expression of the osteoblasts markers *OCN*^65,82^ and *BSP2*^63^, but not of *OPN* and *ALP*. Of note, *OPN*^75^ and *BSP2* belong to the small integrin-binding ligand N-linked glycoproteins (SIBLING) gene family, the integrin dependent signal cascade being one of the key modalities of chondrocytes mechanotransduction^43^ with implication in OA^63^.

Cyclical HPC had instead limited effects on bmMSCs constructs (Fig. 5e, Supplementary Fig. 13). The only statistically significant change was represented by a decrease in the expression of *IHH*. Remarkably, a deficiency in periosteal stem-cell-derived IHH was associated to impaired endochondral ossification processes^83^. These differences in hACs and bmMSCs responses to HPC might be due to dissimilarities in constructs ECM composition and/or mechanotransduction pathways activation. While further investigations are needed, these results indicated that CoC and SMToC models exhibit characteristic responses to loading and reinforce the need to study OCU’s tissue specific response.

Whole OCU tissues demonstrated a distinctive response to loading different from the simple average of hACs and bmMSCs in single culture (Fig. 5e). Specifically, OCU-on-chip exhibited an upregulation of anabolic markers (e.g. the *COL2A1/COL1A1* ratio) while not increasing *MMP13* and *IL8* expression. Furthermore, at day 21, OCU tissues expressed higher levels of *COL2A1, ACAN*, and *MMP13* than single culture counterparts, both in static culture and under cyclical loading. This is possibly indicative of the onset of remodelling processes. Of particular significance might also be the downregulation of the chondrogenic stem cell marker growth differentiation Factor 5 (GDF5) which happened only in the OCU model subjected to HPC (Supplementary Fig. 13). GDF5 expression characterizes synovial progenitor populations involved osteochondral defect repair mechanisms^84,85^ and has been implicated in cartilage repair^86^. However discordant studies point to an increase in GDF5 expression in OA cartilage with respect to healthy hip cartilage samples^87^ or to a reduction of its expression (both in human samples and mice models of OA)^86^

To dissect whether different responses were elicited in OCU layers we used GFP hACs so that, at the end of the culture period, constructs could be digested, sorted, and the two tissues analysed separately (Fig. 5f).

When comparing static and dynamic cultures, both hACs and bmMSCs showed an increase in the expression of *COL2A1*, and bmMSCs also of *ACAN*. Both tissues express also lower levels of *IL8* with respect to static controls. While *IL8* downregulation might be a bias of the sorting (Supplementary Fig. 8), the results are in accordance with bulk RT-qPCR done on whole constructs (which were not digested nor separated though FACS). Finally, HPC led to an increase in the expression of the hypertrophic marker IHH in both tissues and to an hACs specific decrease in *GDF5* expression.

These observations might be reflective of an early OA response. Upregulation of *COL2A1* was previously observed in a mechanically induced OA animal model^88^ and, while OA is characterized by an overall cartilage erosion, an increase in anabolic processes has been reported^77^. The enhanced matrix deposition is however coupled with the onset of hypertrophy and depletion of progenitor markers that might be the reason for subsequent degeneration.

Our data collectively point to the importance of considering both layers constituting the natural OCU when studying joints’ cell responses to aberrant loading. The presented OCU-on-chip seems able to recapitulate the key events of early OA steps and could be a valuable tool in this direction.

### hACs sub-populations with distinct phenotypes characterize loaded cartilaginous and osteochondral on-chip models

OA pathogenesis was related to the lineage progression patterns of chondrocytes with distinctive roles and molecular markers^8,9,90^. Moreover, promising cartilage repair strategies seem to be rooted in the recruitment of specific joint resident progenitor populations^91,92^.

In this framework we aimed to better characterize the diversity of hACs population present in our OCU-on-chip model by performing single cell RNA sequencing (scRNA-seq). Gene expression analysis was applied to hACs from single culture cartilaginous constructs (i.e. the CoC model) and from OCU-on-chip constructs, and exposed (or not) to cyclical mechanical loading (Fig. 6a);

**Fig. 6.**
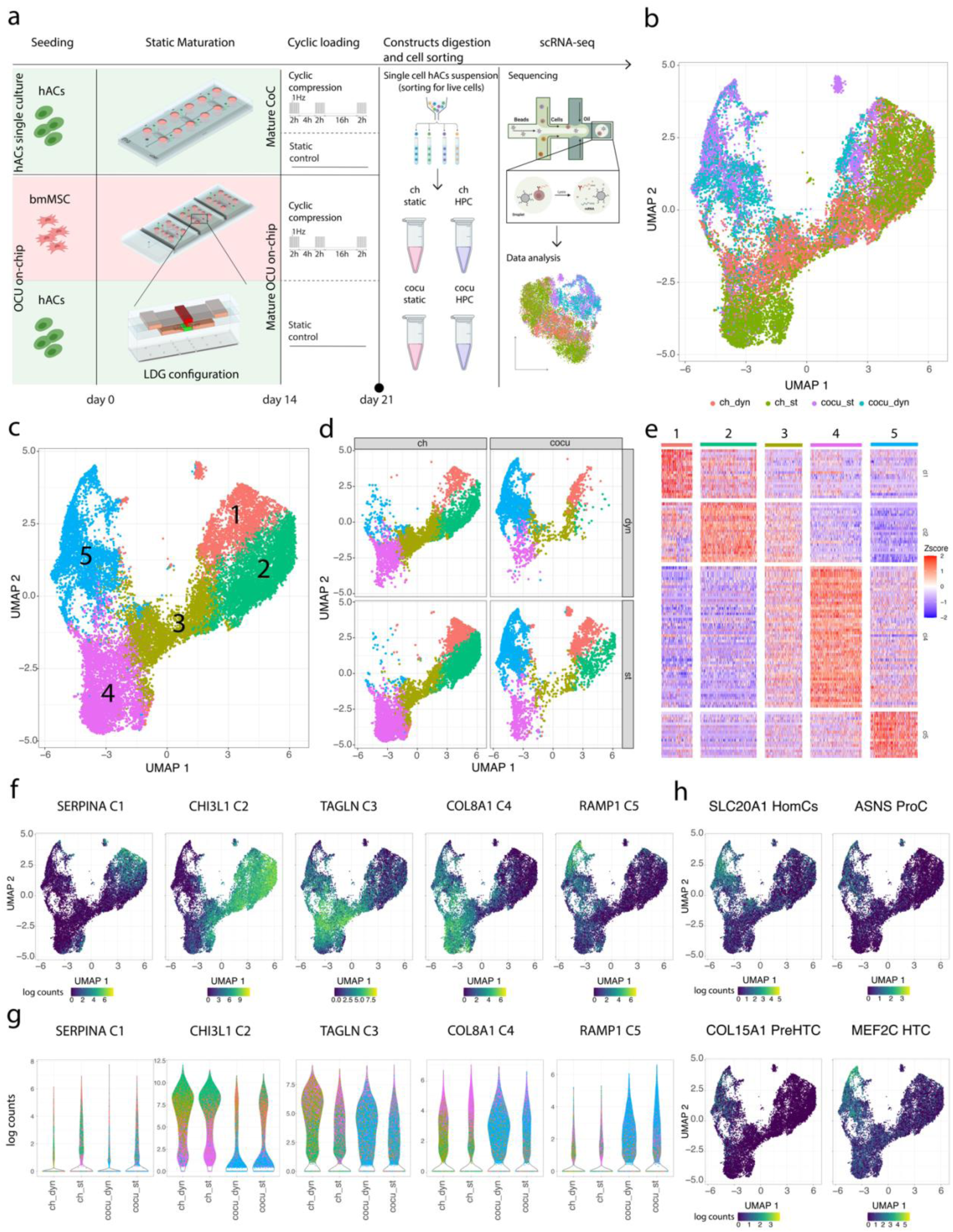
scRNA-seq highlights co-culture and HPC-correlated cellular heterogeneity in hACs populations. **a**, Schematic indicating the experimental plan. Cells were retrieved from cartilaginous and osteochondral constructs trough enzymatic digestion and populations enriched for live cells through sorting. scRNA-seq was performed on hACs only. **b**, UMAP plot of single-cell transcriptomes coloured according to hACs stimulation state (i.e. static vs HPC) and culture conditions (i.e. single vs co-culture). **c,** UMAP cell-clusters representation of scRNA-seq data. The five identified clusters are represented in different colours. **d,** scRNA-seq data UMAP cell-cluster representation divided by stimulation state (rows) and culture conditions (columns). Cells are coloured according to the cluster to which they are assigned adopting the colour scheme of panel **c**. **e,** Heatmap reporting the Z-score of differentially expressed genes in clusters defined in panel **c**. While specific cluster markers could be defined for clusters 1, 2, 4,and 5, cluster 3 had an intermediate state between cluster 2 and 4 without characteristic marker genes. Genes are represented in rows, single cells in columns. **f,** Feature Plots of selected cell-cluster markers. **g,** Violin plots of the above indicated cell-clusters markers. Data points referring to single cells are coloured according to the clusters to which they belong using the colour scheme reported in panel **d**. **h**, Feature plots of selected marker genes for cluster 5 that were previously defined as hACs sub-population markers in samples from OA patients^8^. hACs populations are indicated in the titles.

A total of 24579 cells from 4 different human donors passed quality control steps and underwent further analyses.

Five transcriptionally distinct cell clusters (Fig. 6b-e, Supplementary Fig. 14a, b) were distinguished performing Uniform Manifold Approximation and Projection (UMAP). Cluster marker genes were subjected to gene ontology (GO) analysis using the STRING database^93^.

A clear dependence of the clusters on the culture conditions was apparent (Fig. 6d): hACs from OCU-on-chip, were mainly found in cluster 1 and 5, while cells from the CoC model were assigned mainly to clusters 2, 3 and 4. Markedly, cluster 5 was almost exclusively constituted of cells from the co-culture, while co-culture (slightly) and dynamic loading (majorly) together seemed to have a depleting effect on cluster 2.

Cluster 1 was characterized by expression of *SERPINA1*, *SERPINA5*, *SERPINE2* and *TNFAIP6* which have been reported among the most modulated genes in OA cartilage (as assessed considering patients from the RAAK study^94^). Cluster 2 was characterized by a high expression of *SOD3* and *MAFB*, together with *CHI3L1* and *CHI3L2* which, being expressed also by other clusters, were described to distinguish a chondrocytes subpopulation that decreases in abundance during OA progression^8^ but seems more present in OA samples if compared with healthy cartilage^9^.

Cluster 3, primarily constituted by hACs cultured in single culture and subjected to HPC, was characterized by only 3 markers, namely *COL1A1*, *COL3A1* and *TAGLN*. None of these markers was however exclusive of cluster 3 which seems constituted by cells with an intermediate differentiation state between those of clusters 2 and 4. Cluster 4, characterized by GO terms such as Extracellular Matrix Organization, Collagen Fibril Organization, and Supramolecular Fibril Organization (Supplementary Fig. 14b), was distinguished by markers such as *COL1A1*, *COL1A2*, *COL8A1*, and *FBN1*.

Finally, the co-culture dependent cluster 5 was categorised by *UNC5B*, *RAMP1* and the calcium binding protein *S100A13*, which are associated with GO terms such as Inorganic Diphosphate Transport, System Development, and Multicellular Organismal process. A complete list of all cluster markers can be found in Supplementary File 1 (Tab 1 – Tab 5).

In accordance with previous studies on scRNA-seq of chondrocytes^8–10^, even if various subpopulation (or clusters) could be highlighted, chondrocytes subpopulations seem nevertheless to represent smoothly differentiating cells rather than completely distinct populations. As reported in the heatmap of Fig. 6e (where each column represents one of the cells assigned to a specific cluster and each raw represents a gene), the gene expression of marker genes varies smoothly from cluster 1 to 5. Other authors performed pseudo-space trajectory analyses of chondrocytes from OA patients suggesting a possible correlation between highlighted cell subpopulations and OA progression^8,9^. An immediate transposition of these results to *in vitro* models using *ex vivo* cultured chondrocytes is however not trivial.

Feature plots of selected cluster markers are reported in Fig.6f, violin plots in Fig. 6g. These latter further highlight the dependence of selected markers on co-culture and HPC: *TAGLN* expression which was associated with a preHTCs phenotype^9^, seems to increase with HPC; *CHI3L1* expression was typical of single culture hACs; *COL8A1* and *RAMP1* expressions correlated with both co-culture and mechanical stimulation. To better characterize the general effects of culture type and mechanical loading on hACs, samples were subjected to principal component analysis (PCA) (Extended Data Fig. 5, Supplementary File 2).

Of note, during analyses, gene expression was corrected for the effect of individual donors. This is visible in Extended Data Fig.5a where cells from different donors are randomly dispersed in the PC1-PC2 space.

PCA did not split cells in separate clusters pointing at the similar nature of hACs across conditions. This result is in accordance with an *in vivo* model of OA, where anterior cruciate ligament damage did not result in distinct cell clusters 7 days post injury^10^.

Nevertheless, PC2 showed a clear correlation with the culture modality (i.e. single vs co-culture, Extended Data Fig. 5b), whereas PC3 was indicative of mechanical loading (Extended Data Fig. 5c). The meaning of PC1 was instead less clear, but it seems related to transitions between cell types or state. These results further remark the necessity to distinguish the behaviour of separate hACs populations.

To better understand which cellular processes were related to gene expression across conditions, we defined the top 100 genes contributing the most, either positively or negatively, to PC1 and PC2, and subjected them to GO analysis using the STRING database^93,95^

Corresponding significantly enriched Biological Processes (BPs) and protein-protein interaction networks are reported in Extended Data Fig. 5d, e. Top BPs enriched by genes aligned with PC1 included Skeletal System Development, Ossification, and Cartilage Development (PC1 negative), or Collagen Fibril Organization, Regulation of Cell Migration, and Regulation of Cell Population Proliferation (PC1 Positive). BPs enriched by genes aligned with PC2 included Defence Response, Response to Stress, and Inflammatory Response (PC2 negative), or Cell Cycle, Mitotic Cell Cycle, and Cell division (PC2 Positive). Of note, the top 100 genes contributing to PC2 formed a highly interconnected protein-protein interaction network globally associated with proliferative processes (Extended Data Fig. 5f). One feature of interest was that the PC1-negative PC2 positive area characterized by proliferative and developmental processes was preferentially occupied by hACs from the OCU-on-chip model. This results suggests that by co-culturing hACs with bmMSCs in osteochondral constructs, the former acquire a phenotype more similar to that of chondrocytes in the growth plate during development^96,97^ whose processes are also recapitulated in OA^77^.

These data collectively point to the presence of hACs subpopulations with substantial differences in single cell transcriptomes but also to a close relationship between these different populations. Given the influence of HPC and co-culture, our data also further highlight the necessity of more comprehensive OCU models in the study of OA pathogenesis.

### OCU-on-chip culture better preserve chondrocytes subpopulations with implications in OA pathogenesis

Previously published transcriptome-wide scRNA-seq analyses of cartilage samples from patients undergoing knee arthroplasty for OA led to the classification of chondrocytes into seven molecularly defined subpopulations^8^: Effector chondrocytes (ECs), regulatory chondrocytes (RegCs), proliferative chondrocytes (ProCs), pre-hypertrophic chondrocytes (preHTCs), hypertrophic chondrocytes (HTCs), homeostatic chondrocytes (HomCs), and fibrocartilage chondrocytes (FCs).

To identify analogies between the clusters identified in our CoC and OCU-on-chip models and subpopulations found in OA patients, hACs were re-analysed according to the above mentioned classification^8^.

Specifically, we cross-referenced genes defining our five clusters with those adopted in previous studies^8,9^ (Fig. 6h, Supplementary Fig. 15, Supplementary File 1, Tab 6), and performed a hierarchical clustering of our samples based on previously determined marker genes (Extended Data Fig.6a). These analyses collectively revealed that cluster 2, 3, and 4 (largely constituted by hACs from the CoC model) were broadly associated with PreHTCs and FCs phenotypes, cluster 1 also included markers of RegCs, and cells of cluster 5 (rich in cells form OCU-on-chip constructs) possessed features of HomCs, ProCs, preHTCs, and HTCs.

These results were confirmed by feature plots of selected cell types markers genes which showed different expression by cells of defined clusters (Extended Data Fig. 6b). Nevertheless, while some markers were circumscribed to specific clusters (e.g. HMOX to cluster 1 or COL10A1 to cluster 5) other where ubiquitously expressed (e.g. JUN).

As highlighted by other scRNA-seq studies^8–10^, hACs cannot be easily classified by a restricted set of indicators and subpopulation markers are expressed across cell populations.

To obtain a clearer picture of how hACs cell subtypes were distributed, UMAPS representations were coloured according to a score that defines the fitting of assigning a given cell to a precise cell type (Extended Data Fig. 6c; higher scores are represented in red; Scores were based on the complete list of markers of that population as available from previous studies^8^).

In general, the highest scores for ECs, RegCs, ProCs, HomCs, and HTCs were found at the top of the UMAP representation, where the majority of hACs retrieved from OCU-on-chip of clusters 1 and 5 is located.

A large portion of the cells was assigned a high score for FCs. This phenotype has been associated with terminally differentiated hACs in OA^8,9^ but also to dedifferentiated chondrocytes obtained as a result of their handling *in vitro*^98^. hACs’ tend in fact to dedifferentiate during their 2D expansion and to only partially re-acquire a native phenotype during their subsequent in 3D cultures^98^. hACs used in this study were expanded in 2D up to passage 3, and, particularly in single culture, showed the elongated appearance typical of hACs expanded on plastic surfaces.

Alternative cell sources that would not require expansion *in vitro* are however not available. As highlighted by other authors^56^ and in previous sections, bmMSCs are committed to hypertrophic differentiation (Fig. 3, Extended Data Fig. 3), while hACs from OA patients undergoing joint replacement, even if harvested from unaffected joint regions, behave differently than healthy hACs^99^.

Scores were then combined to annotate cells i.e. to assig them to a subpopulation. Notwithstanding the above-mentioned limitations (and albeit an extremely low number of hACs were assigned to the ECs and ProCs cell types), the OCU-on-chip exposed to HPC was the only experimental group to account for all chondrocytes types identified in OA cartilage samples (Extended Data Fig. 6e).

Together, these data indicate that, while the proposed OCU-on-chip is still affected by the necessity to expand hACs in 2D, hACs retrieved from osteochondral constructs retain (or re-acquire) a more complex variety of populations; the co-culture appears necessary to achieve *in vitro* constructs that preserve, at least partially, the endogenous transcriptional heterogeneity of chondrocytes in OA samples.

These data seem also to indicate that HA positive cells appearing in the cartilaginous compartment of OCU-on-chip constructs (Fig. Fig 4c, d, Fig. 5b, and Supplementary Fig. 11) are HTCs rather than migrating bmMSCs. Cells annotated as HTCs, who are classically associated to mineralization processes^60^, seem to be a consequence of the co-culture.

Beside introducing a taxonomy of OA chondrocytes, Ji et al.^8^ provided a characterization of the transcriptome of cells at different disease stages. We performed hierarchical clustering based on the top 50 genes that they identified as indicative of healthy cartilage/early OA and on the 50 genes indicative of advanced OA grades. As mentioned above, OCU-on-chip derived clusters 1 and 5 are enriched for preHTC and HTC populations, hypertrophy being one of the signatures of OA ^60,61^. They however expressed higher levels of genes associated to milder OA states (Supplementary Fig. 16a, b). This result also confirms our RT-qPCR data where co-cultured samples had an overall more anabolic gene signature (Fig. 5, Supplementary Fig. 13). It is interesting however to notice that regardless the cluster, and of the culture conditions, HPC led to an increase in the expression of *COL5A1* and *DKK3* which have been associated to more advanced OA states in patients cartilage^8^ (Supplementary Fig. 16c).

Other authors constructed a complementary scRNA-seq-based classification of hACs subpopulations based on healthy cartilaginous samples (even if with a very restricted cohort)^9^. Specifically, on top of RegCs, HomCs, and PreHTCs, two different FCs populations (i.e. FCs-1 and FC-s2), and two Cartilage Progenitor Cells (CPCs) populations were defined. Similar to what described above, we compared our clusters with this second classification.

CPCs populations markers are highly expressed in cluster 5, which together with cluster 1, also expresses preHTCs markers (Extended Data Fig. 6e, f). Notably, CPCs markers, among which *BIRC5*, *CDC20* and *CENPF* were expressed by a well-defined set of cells (Extended Data Fig 6f, Supplementary Fig. 16).

CPCs population were found to be depleted in samples from OA patients^9^ but the cell cluster expressing them is constituted by both static controls and cells subjected to HPC. These results highlight a bivalent effect of the co-culture that, on one side enriches hACs for late OA associated populations such as preHTCs and HTC while, on the other, allows to obtain (or maintain) cells with a more progenitor-like transcriptome. Of note the CPC population in our model was also positive for the proliferation markers *MKI67* and *E2F1*^100^ (Supplementary Fig. 16). CPCs have been associated with self-renewal and proliferative properties^101^ but more investigations will be necessary to more deeply characterize these cells.

### OCU-on-chip responses to aberrant mechanical loading highlight ribosome dysfunctions as a possible trigger of OA pathogenesis

We then made use of scRNA-seq data to acquire an unbiased assessment of pathways affected by aberrant mechanical stimuli. Specifically, we determined differentially expressed (DE) genes between cyclically compressed samples and static controls, and discriminated between hACs exposed to HPC in single culture and in the OCU-on-chip model.

DE genes (Fig. 7a) were obtained aggregating scRNA-seq data in *in silico* bulk samples. 193 genes were modulated both in single culture and in co-culture, 359 genes were modulated only in single culture, and 357 genes only in co-culture (Fig. 7b, Supplementary File 3). Additionally, different KEGG pathways were enriched in the two cases (Supplementary Fig. 18) with upregulated ones following HPC in co-culture including Cell Cycle, cGMP-PKG Signalling^70^, and Proteasome. A cross talk between hACs and subchondral layers leads therefore to a different response to mechanical stimuli.

**Fig. 7.**
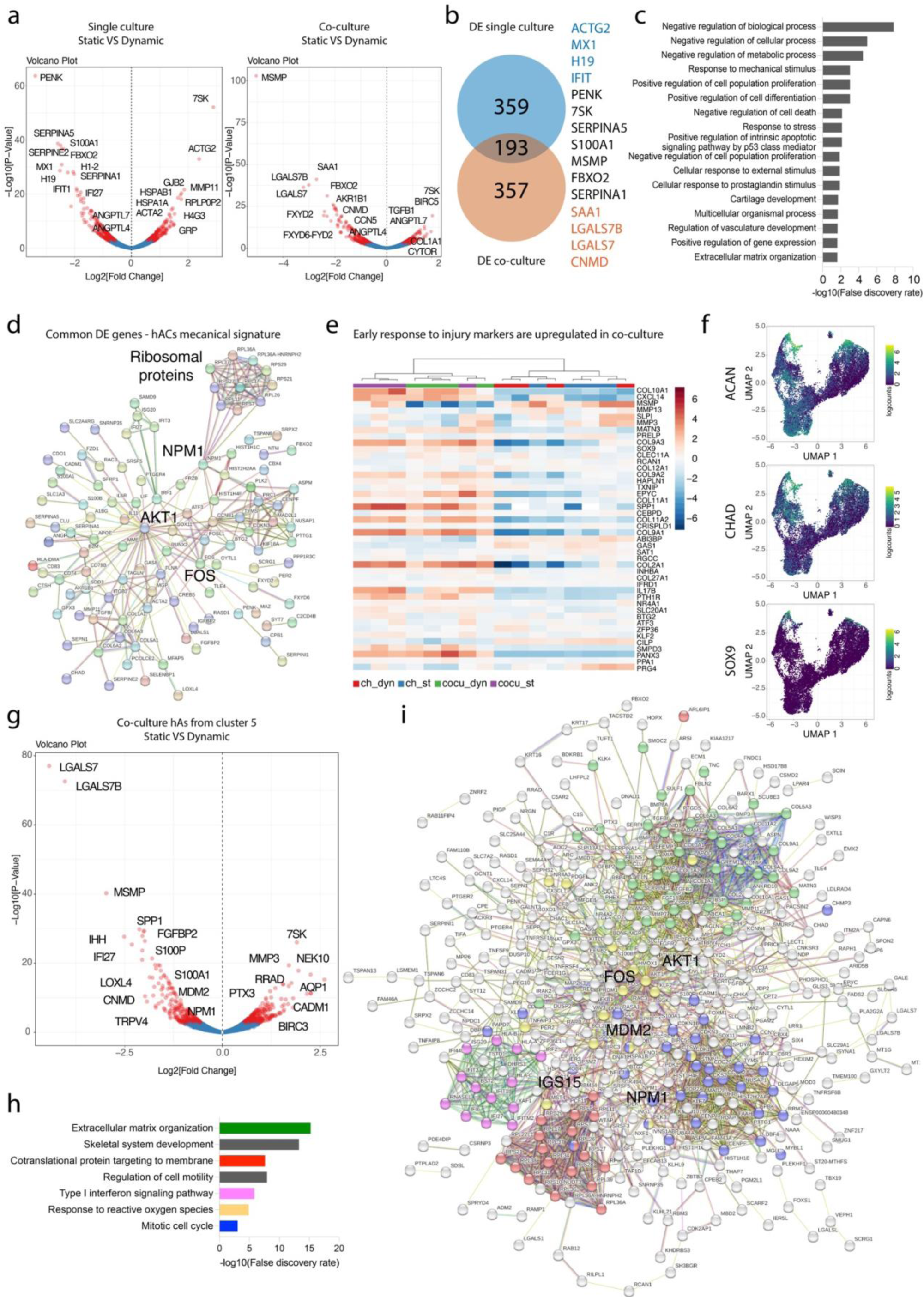
HPC dependent DE genes in OCU-on-chip: pathways and interactions underlying hACs mechanotransduction. **a**, Volcano plot of DE genes obtained comparing static samples and samples subjected to HPC considering *in silico* bulk samples from single culture (left) and co-culture (right) samples. DE genes with adjusted P value < 0.05 are indicated in red. **b**, Common and not common differentially expressed genes in single and co-culture. A few representative genes are indicated on the right. The number of common DE genes is indicated by the overlapping between the two circles. **c**, GO terms (cellular processes) related to genes that are DE in both single and co-culture (GO terms were obtained through STRING). **d**, Protein-protein interaction network relative to common DE genes. Networks were obtained through STRING. Non connected nodes were excluded from visualization. Branches in the network are coloured according to the interaction evidence **.e**, Samples hierarchical clustering based on DE genes 7 days after ACL damage in a mouse model of OA^10^. **f**, Feature plots of representative markers adopted to distinguish chondrocytes in a mouse model of OA^10^. **g**, Volcano plot of DE genes obtained comparing static samples and samples subjected to HPC. DE expression was acquired considering *in silico* bulk samples obtained summing the expression of all cells belonging to cluster 5 and retrieved from co-cultures. DE genes with adj. P value < 0.05 are indicated in red. **h**, GO terms (cellular processes) related to DE genes in cluster 5 comparing static and HPC samples and considering only cells from co-cultures. GO terms were obtained through STRING. **i**, STRING protein-protein interaction network relative to Cluster 5 DE genes. Nodes relative to the GO terms highlighted in panel h are represented with the same colour code. Non connected nodes were excluded from visualization. Branches in the network are coloured according to the interaction evidence.

Starting from these considerations we set, at first, to identify fundamental regulators of hACs response to pathological mechanical loading focusing on genes that were modulated by HPC both in single and in co-culture. Enriched BPs (Fig. 7c) comprised Negative regulation of Biological Processes, Response to Mechanical Stimulus, Cartilage Development, and Extracellular matrix Organization, but also Positive Regulation of Intrinsic Apoptotic Signalling Pathway by p53 class mediator. HPC affects therefore chondrocytes at a metabolic level, acting on matrix production, but also at a higher level directly associated with proliferation and apoptosis^102^.

Common modulated genes were used to visualize possible protein-protein interactions (Fig. 7d). The most connected hub in the network was represented by AKT Serine/Threonine Kinase 1 (AKT1), which was associated to cartilage calcification during endochondral ossification^103^, and to bone and joint specification in embryonal development^104^.

Other DE genes comprised known markers of matrix degradation and deposition (e.g. *COL1A1*,*COL5A1*,*COL6A2*); genes associated to the onset of hypertrophy (i.e. *FRZB*^39^); *TGFB1*, whose signalling was demonstrated to be dysregulated in cartilaginous areas corresponding to altered architecture of subchondral bone^54^; *SERPINE*s that were among the most modulated genes highlighted in the RAAK study^94^; and *FOS*, which was previously associated to mechanotransduction in chondrocytes^31^.

A feature of particular interest is that, among those affected by compression, we found numerous genes coding for ribosomal proteins (e.g. *RPS29*, *RPS7*, *RPL26*, *RPL37*). *RPS29* was previously demonstrated to be expressed at higher levels in OA patients^8^. OA has been linked to alterations in ribosome biogenesis, ribosomal RNA transcription, and preferential protein translation^105^. While OA is classically viewed as the result of an imbalance between anabolic and catabolic processes^106^,however, limited attention was given to ribosomes, which are responsible for protein translation from gene transcripts. A recent view of the pathology fine-tunes the anabolism/catabolism paradigm characterising OA as an acquired ribosomopathy^105^. Our data corroborate this interpretation establishing moreover a previously unknown link between ribosome alterations and aberrant mechanical stimuli.

Of note, the majority of modulated genes coding for ribosomal proteins was connected to nuclephosmin (*NPM1*, Fig. 7d). In the field of cancer biology, NPM1 is known to be implicated in processes such as mitotic spindle assembly, chromatin remodelling, DNA repair, embryogenesis and ribosome synthesis^107,108^. To the best of our knowledge NPM1 was not previously associated to OA; it was however reported to be modulated by mechanical stimulation (specifically stretching) in tendon derived Stem cells^109^. Acting upstream of ribosome biogenesis, NPM1 could be a promising occasion for therapeutic intervention.

Subsequently, we aimed at better understanding differences in DE genes between CoC and OCU-on-chip models.

Given that the response to HPC of OCU-on-chip seemed more in line with findings relative to early OA phenotype (Fig.5), we performed hierarchical clustering of the samples based on a set of genes whose expression was altered seven days post intervention in an anterior cruciate ligament (ACL) injury OA mouse model^10^. hACs coming from single culture cartilaginous construct and those coming from osteochondral constructs clearly clustered separately. OCU-on-chip derived hACs expressed higher levels of early OA markers (Fig. 7e). Furthermore, classic cartilaginous markers that were adopted to select mice chondrocytes for analysis after ACL injury (i.e. *ACAN*, *CHAD* and *SOX9*) were more expressed in clusters 1 and 5 which are prevalently constituted by hACs from the OCU-on-chip (Fig. 7f). The cartilaginous layer in OCU-on-chip, consequently better recapitulates the transcriptional profile of joint chondrocytes and molecular changes happening in the first disease states.

In light of these results, we analysed DE genes more in depth, concentrating on hACs from co-culture samples only, and dissecting the differences in each cluster (Fig 7g, Extended Data Fig.7a).

hACs from the different clusters exhibited distinctive responses to loading. Only an exiguous number of DE genes was common to all clusters; these included *SPP1*, *MMP7*, and *COMP*, together with R*PL36A-HNRNPH2* and *RPS7* two ribosomal proteins (Extended Data Fig. 7b).

Due to the dependence of cluster 5 on the co-culture, we specifically focused on this population. Of note, studying cells belonging to cluster 5 and exposed to HPC, more than a thousand genes were modulated comparing single culture and co-culture. These included markers of ossification and hypertrophy such as *COL10A1*, *IBSP* and *SNORC* (Extended Data Fig. 7c) but also numerous indicators of cell division (Extended Data fig. 7d, e). The transcriptional heterogeneity maintained in our OCU-on-chip might therefore be necessary to investigate phenomena observed in clinical samples such as chondrocytes hypertrophy^60,110^ and multiplication^111^.

Analysing the genes modulated by HPC in cluster 5 (and in co-culture cells only) revealed a highly complex picture (Fig. 7g-i).

Interestingly, downregulated genes in cluster 5 included transient receptor potential vanilloid 4 (TRPV4), a ion channel that transduces mechanical loading of articular cartilage via the generation of intracellular calcium ion transients^112,113^. *TRPV4*, demonstrated to affect age related OA^113^ and previously considered as a therapeutic target for OA^114^, was not modulated in other clusters nor it was previously known to be regulated at a transcriptional level following injury or mechanical stimulation.

Among genes with the highest fold change we also found *MSMP,* which was downregulated as in the above mentioned mouse model of early OA^10^, and *LGALS7* (together with its paralog *LGALS7B*) a galectin implicated in cell-cell and cell-matrix interactions on top of cell differentiation and migration^115^ which was also downregulated.

Looking at clusters of connected proteins, 4 predominant drivers could be highlighted: Extracellular Matrix Organization, Cotranslational Proteins Targeting to Membrane (i.e. ribosomal proteins), Type I Interferon Signalling, and Mitotic Cell Cycle (Fig. 7h, i). Proteins interaction were moreover centred around *AKT1* and *FOS*, correlated, among other pathways (Supplementary File 3), to Response to Reactive Oxygen Species.

As a last consideration we hence focused on hubs (i.e. points of connection) of these macro areas in the protein-protein interaction network (Fig. 7i).

Hubs in the Extracellular Organization pathway included expected OA markers such as *MMP2*, and *3*, *COL1A1* and *COL10A1*, *TGFB1* and *3*, *VEGFB*, and *SPP1*; type I interferon related proteins and ribosomal proteins were tied to ISG15 an ubiquitin cross reactive protein associated with rheumatoid arthritis(RA)^116^; cell cycle proteins largely connected to the transcriptional factor *FOXM1*, and to the kinase inhibitor *CDKN3*s, beside the CPCs marker *BIRC5*.

Interestingly, all these three protein clusters were linked to *MDM2* an E3 ubiquitin-protein ligase that mediates the ubiquitination of p53 and was associated to RA^117^. Downregulation of *MDM2* could lead to increased apoptosis. Moreover, the protein is known to interact with the ribosome biogenesis-related *NPM1*^108107^, one of the central HPC reactive gene hubs highlighted in Fig. 7d.

These data collectively introduce new fundamental knowledge of the processes started by mechanical stimulation of hACs. They also indicate that transcriptional alterations following HPC, and supposedly OA, at least following traumatic events or due to joint misalignment, go far beyond classically considered metabolic^106^ and inflammatory^118,119^ pathways.

Ulterior mechanistic studies will be necessary to properly elucidate the cause-effect relationship of the highlighted pathways. Nevertheless, through the use of our OCU-on-chip model we identified different candidate triggers of OA pathogenesis, particularly in connection to ribosome biogenesis. Alteration of the expression of genes coding for ribosomal proteins was common to both single culture and co-culture, was shared among different cell clusters, and represented a set of majorly connected proteins looking at the OCU-on-chip specific cluster 5. Our OCU-model also highlighted possible upstream regulators of these processes, such as *MDM2* and *NPM1* which, extensively studied in cancer research^108^, could represent new venues for the treatment of OA.

## Conclusions and Outlook

In this work we introduced an innovative microscale technological platform and a biological OCU model; we then employed them synergistically to study what the mechanobiology contributions to the etiogenesis of osteoarthritis (OA) could be. We propose, a new vision of the pathology where the classical paradigm of OA as the result of catabolism and anabolisms imbalance is only the downstream effect of alterations in ribosomes biogenesis^105^ which are elicited by application of a mechanical stimuli.

Numerous therapeutic options that have been proposed for OA concentrated solely on the low-grade inflammatory state of the disease^119^. While this approach has given some results for rheumatoid arthritis, the failure in translating it to OA^120^ indicates that we are not addressing the underlying disease cause. Our results reinforce the connection between OA and mechanical risk factors emphasising a fundamental role of mechanotransduction for processes spanning from ribosome biogenesis to cell cycle and apoptosis.

Some considerations on the study have, nevertheless, to be made, and some limitations highlighted. Coupling the newly introduced VBV concept with a pneumatic compartment^11,25^ we recapitulated the functional anatomy and the compression levels experienced by cartilage and bone in articulations. The multi-layer fabrication technology allows furthermore to customize the features of top and bottom compartments. Our approach could therefore be generalized to applications beyond the osteochondral interface.

We successfully obtained bi-phasic tissues transitioning from a hyaline cartilage-like layer to a mineralize subchondral tissue. To do so we used a developmentally inspired endochondral ossification process where a cartilage structure prefigures the presence of long bones^121,122^. bmMSCs based tissues retained therefore some cartilage traits despite their mineralization and the high expression of bone markers. A more comprehensive bone model would incorporate osteoblasts, osteoclasts, but also vasculature, given the remodelling processes and cartilage vascular invasion typical of OA^61^. Such features might be the centre of further studies.

Of note, the LDG configuration was designed so that it is feasible to inject the hydrogel in the top compartment at a later time point with respect to the one in the bottom compartment. hACs can therefore mature into cartilaginous constructs before the incorporation of a second layer (e.g. with vasculature and osteoclasts) thus de-coupling respective maturation phases that can be optimized separately.

Our OCU-on-chip model remains however an over simplification of the joint environment. Our mechanical stimuli are limited to compression while cartilage in the knees is subjected to a complex range of motions including compression but also shear and rotation^33,123^.

Being aware of these limitations, our model features however a superior level of complexity with respect to existing systems and can be used to understand the role of subchondral bone in the development of OA both form a mechanical standing point and to study inter tissue cross talk.

On this regard our scRNA-seq analyses revealed a profound effect of the subchondral layer on hACs. The co culture seems necessary to better preserve *in vitro* hACs subpopulations found in OA cartilage^8^, maintain progenitor-like cells^9^, and to obtain a gene expression more reminiscent of early OA^10^. Of note, among the differentially expressed genes we found angiopoietin like 4 and 7 (*ANGPTL4* and *ANGPTL7*). An ANGPTL3 derivative, supposedly acting on cartilage progenitor, is presently in a phase II clinical trial for the treatment of OA^92^. Our OCU-on-chip could therefore be an ideal candidate to investigate the mode of action of such OA treatments.

Finally, it is important to notice that, while we focused only on given DE genes in our analyses, our data represent a broad overview of the mechanotransduction machinery of hACs and could highlight different promising targets for a pharmacological therapy of OA.

## Methods

### Devices design and fabrication

Microscale systems were realized in PDMS (Sylgard 184; Dow Corning), polymerized with defined casts at a 10:1 weight ratio of base to curing agent. All PDMS parts were cured for at least 2 hours at 65°C. Final devices were produced bonding four different layers (Fig. 1c, Supplementary Fig. 1).

Unpatterned actuation membranes were achieved pouring defined quantities of PDMS into a Petry dish (Sigma Aldrich; 100 mm x 15 mm polystyrene Petri dishes) containing a clean silicon wafer so to obtain a final thickness of 800 µm. This thickness was demonstrated to allow for the actuation pressure (i.e. the pressure necessary to obtain contact between the membrane and compression chambers pillars bottom surface, Supplementary materials, Supplementary Fig. 4) to depend exclusively on the bending stiffness of the membrane itself, thus preventing the effective compression level achieved during culture to vary as a result of tissue maturation. Polymerization was achieved on a levelled surface.

All other layers were obtained by replica moulding of appropriately designed master moulds. Layers schemes were realized through Computer Aided Design (CAD) Software (AutoCad 2020, Autodesk). Master moulds were fabricated in a cleanroom environment (Class ISO6) using multilayer direct laser writing (Heidelberg MLA100) of SU-8 2035 or SU-8 2100 (MicroChem) onto 102 mm silicon wafer substrates.

Different designs were realized for the top compartment i.e. NDG, LDG, and DG; final features had a nominal height of 150 µm, while their width depended on the design version. Schematics of the different features are reported in Extended Data Fig.1. All top compartment geometries were realized in different versions, with a central channel width of 300 µm, 400 µm, and 500 µm. After an assessment of the precision and an optimization of the alignment of top and bottom compartments (Supplementary data, Supplementary Fig. 3) devices with top compartments central channel of 300 µm were used for following experiments.

Features in the bottom compartment had a cross section of 1800 µm (width) x 293 µm (height) and consisted in three different layers. The three SU-8 layers were realized sequentially: gap layer (height: 43 µm), pillars layer (height 100 µm) and necking layer (height 150 µm). Pillars in the compression chamber layer, positioned 30 µm apart, were designed with a T cross section, each arm of the shape being 300 µm long and 100 µm wide. The T tails facing the culture medium channels were rounded to avoid air entrapment during the medium filling phase. Pillars extremities were tapered to obtain a wider contact angle and better confine the hydrogel formulation in the central channel during injection.

The actuation chamber layer was designed so that three actuation chambers could be connected with the same actuation inlet (Fig. 1b, c). Actuations chambers had a cross section of 3397 µm (width) x 50 µm (height). Six rows of round pillars (diameter 28 µm) were introduced to prevent the chambers from collapsing.

Top compartment and actuation chamber were realized pouring PDMS on master moulds to get heights of roughly 3 mm and 1.5 mm respectively. PDMS stamps were then detached from the master moulds, cut, and appropriate holes bored for hydrogel inlets (diameter 1mm), culture medium reservoirs (diameter 4 mm) and actuation tube inlet (diameter 1.5 mm).

The bottom compartment layer was realized pouring a small quantity of PDMS (roughly 4 ml) on the master mould and then spinning the master itself so that a thin PDMS layer would cover everything but the top surface of the necking geometry. An optimization procedure was carried out to find the optimal spinning velocity which consisted in a 5 second ramp to 500 rpm held for 15 sec, and followed by 30 seconds at 230 rpm.

The PDMS stamps of the top culture chamber were bonded directly on this spinned thin PDMS layer before removing it from the master mould to ease its handling. Subsequently, the appropriate hydrogel inlets and outlets and culture medium reservoirs holes (3mm) were bored through both layers. A thick PDMS slab was bonded on the unpatterned membrane and a hole (1.5 mm) bored through both layers in correspondence of the actuation inlet of the actuation chamber layer. The increased thickness provided by the slab was required to assure retention of the tube providing pressure for devices mechanical actuation. Actuation chamber and actuation membrane (plus the PDMS slab) were bonded first and united with the top compartment-bottom compartment assembly in a second moment (Supplementary Fig. 1) All procedures were performed treating surfaces to be united with air plasma (Harrick plasma) and bringing them in conformal contact for at least 30 min at 80°C to achieve irreversible bonding.

Single culture microfluidic devices (Fig. 3a) were produced in PDMS similarly to what specified above and as previously described^11^. The same procedure was adopted to produce co-culture horizontal devices (Fig. 4a) in which two 300 µm wide channels, flanked by T-shaped pillars, are separated by a row of hexagonal channels (width of 50 μm, length 90 μm, spacing between two consecutive pillars 50 μm).

### Device geometrical characterization

Devices geometrical features were measured through observation of their cross sections (Fig. 1f). Thin PDMS sections (1-2 mm thick, n >6 for each device version) were cut with a razor blade and observed through a brightfield microscope connected to a digital camera (AmScope MU500); features dimensions were measured from acquired images. Image analysis was performed using Image J. Assembled top and bottom compartment PDMS layers were also imaged through SEM (LEO 1525 Field Emission Scanning Electron Microscope). PDMS devices were coat-sputtered with a few nanometers of gold. Images were acquired adopting an Electron High Voltage (EHV) of 10.00 kV.

### PEG precursor production and PEG hydrogel preparation

PEG hydrogels were produced as previously described.^24^ 1 ml FXIII (200 U ml^−1^; Fibrogammin; CSL Behring) was activated with 100 μl thrombin (20 U ml^−1^; Sigma–Aldrich) for 30 min at 37 °C and the resulting activated FXIII (FXIIIa) was stored in small aliquots at −80 °C. Eight-arm PEG vinylsulfone (molecular weight: 40 kDa; NOF Europe) was functionalized with peptides that contained either an FXIII glutamine acceptor substrate (Gln peptides; NQEQVSPL-ERCG-NH2; Bachem) or an MMP-degradable FXIII lysine donor substrate (Lys-MMPsensitive peptides; Ac-FKGGGPQGIWGQ-ERCG-NH2; Bachem), resulting in 8-PEG-Gln or 8-PEG-MMPsensitive-Lys precursors, respectively. A stoichiometrically balanced solution of 8-PEG-Gln and 8-PEG-MMP sensitive-Lys was mixed for an indicated final dry mass content of hydrogel precursors in Tris buffer (50 mM Tris (pH 7.6) and 50 mM calcium chloride, leaving a spare volume of 10% v/v for the addition of cell culture medium and cells. The hydrogel cross-linking was initiated by adding 10 U ml^−1^ FXIIIa and vigorous mixing. Adopted Hydrogels had a final concentration of 2% PEG precursors and contained cells at a 50 x 10^6^ cells ml^−1^ density.

### Finite Element analyses

An FE model was introduced to (i) confirm the application of two different and defined average strain levels in the constructs hosted in top and bottom compartments with low dependence on the properties of the constructs themselves, (ii) evaluate the strain field in the different versions of the device, namely NDG, LDG, and DG, and (iii) estimate the effective strain field applied when compressing mature cartilaginous constructs.

Cartilaginous constructs/PEG hydrogels were defined as having a BPE constitutive relation which describes the mechanical response of a homogeneous continuous constituted by an elastic solid phase and an incompressible inviscid liquid phase. The BPE model accounts for the strain and time dependent behaviour resulting from phases interaction but neglects the intrinsic viscoelasticity of the solid phase^124^. Notably, eventual inhomogeneities present in the construct are not captured in the model which assumes uniform properties throughout the control volume. As such, the BPE model underestimates short term-reaction forces of cartilage-like constructs and hydrogels upon compression, but allows accurate prediction of their strain field^124,125^.

Abaqus 6.14 (Abaqus FEA: Dassault systems), formerly established to be adequate in the biomechanical description of biphasic tissues^125^, was adopted to perform computations. Given the device repetitive inner geometry, a minimal unit volume constituted by the PDMS region corresponding to two T-shaped posts facing each other and the constructs volume in-between was adopted in simulations, so to reduce the computational cost (Fig. 2a). Cartilaginous constructs/PEG hydrogels volume was divided in two regions to simulate the presence of two different constructs layers (Fig. 2a blue and red areas). The PDMS region was described as comprehensive of two posts in the bottom chamber, the PDMS VBV necking geometry, and two posts (or the alternative geometric features) of the top compartment. All three versions of the device were simulated.

Given the symmetries presented by the chosen volume along the X and Y directions (Fig. 2a), this repetitive unit was further reduced to a ROI constituted by half of a post and a quarter of the hydrogel using cinematic boundary conditions. The final region used in simulations is the one shaded in Fig. 2a. which has a thickness of 165 μm along Y accounting for half a post and half of the hydrogel region between posts (nominally 30 μm). The total height of the PDMS portion was 443 μm, given by 143 μm of the post, 150 μm of the connection region and 150 μm of the top compartment. The hydrogel region was divided as follows: bottom construct (height 150 μm), and VBV necking plus top construct (height 150 μm + 150 μm respectively). The edges of the PDMS structures were rounded with a 9 μm curvature radius in conformity to measurements performed on physical devices.

The PDMS region was represented using either twenty-node quadratic elements with hybrid formulation (C3D20H elements: Abaqus) or ten-node quadratic modified tetrahedral elements with hybrid formulation (C3D10MH elements: Abaqus). Hydrogels were meshed using eight-node linear hexahedral elements with hybrid formulation and trilinear pore pressure (C3D8PH elements: Abaqus), as required for porous materials. The characteristic dimensions of the elements varied across the hydrogel volumes with smaller elements adopted in the lower part of the bottom compartment constructs, subjected to higher strain levels (Supplementary Fig. 5a). PDMS region’s elements dimension varied accordingly, depending on the dimension of the hydrogel elements at the interface. A mesh sensitivity analysis was conducted to assure the consistency of the solutions. (Supplementary Fig. 5b). The ratios between the dimensions of the elements in the different areas were kept constant varying the total number of elements. A final total number of 51282 element was adopted for the hydrogel region adopting elements with average characteristic dimensions of 12, 5, and 3 μm in the different regions of the hydrogel volumes. The total number of elements adopted to describe PDMS parts depended on the particular geometry (29809 elements for the NDG configuration, 32930 elements for the LDG configuration, and 29502 elements for the DGconfiguration).

Interactions between model parts were modelled using a surface-to-surface contact. Hard contact between Cartilaginous constructs/PEG hydrogels and PDMS was modelled assuming tangential perfect lubrication. The top surface of the PDMS compartment was fixed with an Encaster boundary condition to model its continuity with the tick PDMS top compartment layer. In the case of the non-diffusive top geometry, the outermost lateral part of the PDMS region was also fixed simulating continuity with lateral walls.

Displacements along the Z directions were impeded on the top surface of the hydrogel compartment simulating the presence of the PDMS ceiling. A zero pore pressure was assumed on the hydrogel lateral portions not in contact with the PDMS to allow fluid outflow.

PDMS was described as a non-linear elastic material adopting the Mooney-Rivlin strain energy function equation:

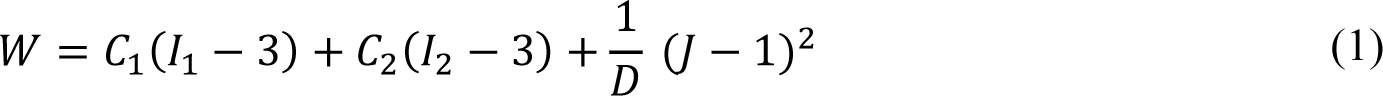

Where I_1_ and I*_2_* are the first and second invariant of the right Cauchy-Green strain tensor C, J is the determinant of the deformation gradient tensor F representing the ratio between the deformed and initial volumes of the object, and C_1_, C_2_, and D are the constitutive parameters of the model. C_1_, and C_2_, were assumed respectively equal to 254kPa and 146 kPa, as reported in the literature for PDMS with a 10:1 base to curing agent ratio^26^. D was set equal to zero assuming a perfectly incompressible material.

As introduced, hydrogel and cell laden constructs were described with a BPE constitutive behaviour. Poisson’s ratio was fixed at 0.33, as previously reported for similar materials^30^, the specific weight of the permeating fluid was set to 9.956 x 10-6 Ncmm-^327^. The description of a BPE model in Abaqus makes use of the specific material permeability defined as 𝐾_𝑠_ = 𝛾_𝑤_𝑘, where 𝛾_𝑤_ is the permeating fluid specific weight and k is the absolute permeability. Ks was set to 3×10^-4^ mm s^-1^ as previously done for similar hydrogel formulations^27^. The initial void ratio e=dVw / dVg, where dVw is the volume of the fluid phase and dVg is the volume of the solid phase, depended on the considered material. An initial value of 45 was adopted for the 2% dry mass PEG-based hydrogel formulation; an e value of 7.5 was used for mature cartilaginous-like constructs, accounting for the presence of both hydrogel initial dry mass and cellular volume fraction^126^. Complete fluid saturation was assumed in all occasions.

PEG based hydrogel Young’s modulus E was assumed equal to 1.9 kPa, as deduced from an estimation based on rheological data,^24,127^ and confirmed by IT-AFM of the gel in the construct. The E modulus of the cartilaginous constructs, estimated through IT-AFM indentation as reported below, was set to 3.66 kPa.

The Abaqus solid consolidation option was adopted to perform a transient analysis implementing an automatic Δ𝑡 incrementation with a minimum time increment of 10^-8^ seconds and a maximum pore pressure incrementation allowed for time increment set to 10^-5^ MPa. The volumetric strain compatibility tolerance for hybrid elements was set to 10^-4^. Compression was applied imposing a 43 μm vertical displacement of the hydrogel bottom surface mimicking the deflection imposed by the membrane in the physical device. Displacement was ramped in a time frame of 0.5 seconds accounting for the 1 Hz frequency adopted in cyclical mechanical stimulation during experiments.

Strain fields were evaluated through the nominal strain components actin in the X, Y, and Z direction and calculated according to Abaqus definition as:

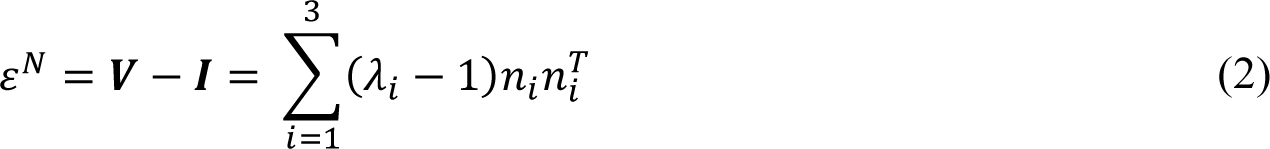

Where 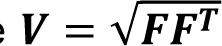 is the left stretch tensor (F being the deviatoric deformation gradient), 𝜆_𝑖_ are the principal stretches, and 𝑛_𝑖_ are the principal stretch directions in the current configuration.

Mean nominal strains in the different compartments, namely bottom and top, were calculated averaging the elements strains calculated in the centroid. To compensate the bias introduced by the usage of higher mesh densities in certain model regions (Supplementary Fig. 5a), the average strains were calculated separately for each of the areas with different mesh dimensions and averaged subsequently weighting for the volumetric fraction of the areas.

### Indentation Type Atomic Force Microscopy (IT-AFM) measurements

AFM indentation was adopted to determine the local Young’s Modulus, E, (i) of cartilaginous constructs, (ii) of the PEG-based hydrogel adopted to encapsulate chondrocytes, and (iii) of the beads laden PEG hydrogels used to assess the effect of substrate mechanical properties on cartilaginous constructs response to loading. After 14 days of static maturation, or immediately after hydrogel polymerization for PEG/ PEG + beads hydrogels, devices were opened to expose the constructs, peeling of the actuation membrane and chamber. Devices were cut with a razor blade to minimize the presence of excessive PDMS and glued onto a plastic culture dish (Sigma-Aldrich, 30mm diameter), with the exposed cellular construct facing upward. Two components epoxy resin (Araldite Rapid, Huntsman corporation) mixed with black dye to increase image contrast was adopted for sample fixation. Measurements were performed at room temperature in degassed PBS.

Measurements on cartilaginous constructs ad empty PEG hydrogels (Fig. 2b,c) were conducted using the ARTIDIS ADO (Automated Device Operation) AFM (ARTIDIS AG, Basel, Switzerland). Indentations were performed using a silicon nitride cantilever (DNP-S10 D, Bruker AFM Probes, Santa Barbara, USA) with a nominal stiffness of 0.06 N m^-1^ and a pyramidal tip of approximately 6 μm of height and a typical radius of the uncoated tip of 10 nm. The exact spring constant of each adopted cantilever was determined through the thermal tune method^128^. Deflection sensitivity was determined directly on the dish as previously reported^34^. Three separate constructs were measured for both experimental conditions, for each construct at least 5 different force maps were acquired at random sample locations. Force maps with dimension of 20 μm x 20 μm, were acquired with a submicron resolution, each force map containing 20 x 20 force curves. Each force curve was recorded at an indentation speed of 16 µm s^-1^. Force displacement curves were corrected for tilt and tip-sample displacement as previously documented^34,129^. Both Forward and Backward force-displacement (F-D) curves were recorded. Backward curves were adopted to extract average samples Young’s modulus adopted in FE simulations making use of the Oliver-Pharr model^130^ as indicated to be suitable for Hydrogels and soft materials^131^. An elastic modulus histogram was derived for each experimental group and mean and standard deviation calculated.

Measurements on beads laden PEG hydrogels (Extended Data Fig. 2c,d) were performed using JPK NanoWizard 4 AFM (Bruker Nano GmbH, Berlin, Germany). Indentations were performed with Sharp Silicon Nitride tips with radii of around 10 nm, a tip height of 2.5-8 μm, mounted on soft triangular shaped Silicon Nitride cantilevers (MSCT-A, Bruker AFM Probes, Camarillo, USA) with nominal spring constants of 0.07 N/m The actual spring constants was determined prior to every experiment by using the Sader method. Force spectroscopy was performed with a load of 800 -1000 pN, with indentation depths ranging of between 500 and 1000 nm. The loading and unloading speed was set to 10 μm/s. Three samples of each condition (0.1% Beads, 5% Beads) were probed recording F-D curves at two different random locations over a scanning area of 40 x 40 μm (32 x 32 force curves) for each sample. E values were calculated from Backward F-D curves with the JPK Data Processing software by using the quadratic pyramid model. An elastic modulus histogram was derived for each experimental group.

### Human cartilages sample collection, chondrocyte isolation and expansion

Cartilage specimens from a total of 8 patients (age: 56 ± 13, 5 males, 3 females) with no clinical history of OA and/or without evident signs of cartilage fibrillation in the harvesting area were collected working under general consent of the hospital after informed consent from the relatives and in accordance with the Ethical Committee of northwest and central Switzerland (EKNZ). Samples were minced and digested enzymatically to isolate human articular chondrocytes (hACs) as previously described^132^.

Briefly, hACs were isolated using 10 ml of a 0.15% (w/v) type II collagenase (300 U mg^−1^; Worthington Biochemical Corporation) solution per g of tissue (37 °C, 22 hours) and resuspended in Dulbecco’s modified Eagle’s medium (DMEM) containing, 4.5 mg ml^−1^ d-glucose, 0.1 mM non-essential amino acids, 1 mM sodium pyruvate, 100 mM HEPES buffer, 100 U ml^−1^ penicillin, 100 μg ml^−1^ streptomycin, 0.29 mg ml^−1^ l-glutamine and 10% foetal bovine serum (FBS), i.e. complete medium.

Isolated hACs were counted using Trypan blue (Thermo-Fisher), plated in culture flasks at a density of 10^4^ cells cm^-2^ and expanded, (in a humidified incubator, 37 °C, 5% CO_2_), in complete medium supplemented with 1 ng ml^−1^ of transforming growth factor-β1 (TGF-β1) and 5 ng ml^−1^ of fibroblast growth factor-2 (FGF-2) which were previously demonstrated to increase chondrocytes proliferation and foster re-differentiation^133^. Passage 3 hACs were used in experiments if not otherwise stated.

Further cartilage samples from 3 patients (age: 64 ± 8, 3 females) undergoing total or unicondylar knee replacement were harvested from preserved areas of the dorsal part of the femoral condyles working under general consent of the hospital after informed consent from the relatives and in accordance with the Ethical Committee of northwest and central Switzerland (EKNZ). Separate samples were collected from cartilage superficial and deep zones. Samples were digested as described above and used for RT-qPCR analyses.

### Human bmMSCs collection, isolation, and expansion

Human MSCs were isolated from bone marrow aspirates and processed as previously described^134^. Marrow aspirates (20 mL volume) were harvested from 5 healthy donors (age: 34 ± 13, 3 males, 2 females) during routine orthopaedic procedures involving exposure of the iliac crest. Bone marrow harvesting was performed after informed consent and in accordance with the local ethics committee (University Hospital Basel, Switzerland). A bone-marrow biopsy needle (Argon medical devices) was inserted through the cortical bone, and the aspirate was immediately transferred into plastic tubes containing 15,000 IU heparin. Isolated bmMSCs were counted using Crystal violet, plated in tissue culture flasks and expanded in a humidified 37°C/5% CO2 incubator in minimum essential medium eagle - Alpha modification (α-MEM), containing 10% FBS, 4.5 mg/mL D-glucose, 0.1 mM nonessential amino acids, 1 mM sodium pyruvate, 100 mM Hepes buffer, 100 Ul ml ^-1^ penicillin, 100 μg ml ^-1^ streptomycin, and 0.29 mg ml -1 L-glutamate, and further supplemented with 5 ng ml ^-1^ of FGF-2 (R&D Systems). Medium was changed twice a week. After approximately 10 days, when they were about 80% confluent, cells were rinsed with phosphate buffered saline (PBS), detached using 0.05% trypsin/0.53 mM EDTA and replated at 5×10^3^cells cm-^2^. Passage 3-4 bmMSCs were used in experiments if not otherwise stated.

### Constructs on-chip maturation

Cartilage and subchondral tissue microconstructs were achieved embedding hACs or MSCs into an enzymatically crosslinkable and degradable PEG based hydrogel^24^ with a final dry mass content of 2% prepared as described above. Biphasic constructs were generated through subsequent injections of appropriate cell laden hydrogels in bottom and top compartments (e.g. with GFP positive and negative hACs seeded respectively in the two compartments, or with MSCs in the top compartment and hACs in the bottom one) in the VBV device or with subsequent injection in the left and right compartments in horizontal co-culture devices.

The polymer precursor solution was mixed with cells so that a final density of 5×10^4^ cells ml^-1^ could be obtained. The cell-polymer precursor solution was then mixed with 10 U ml^-1^ of thrombin activated factor FXIIIa, immediately manually injected into microscale devices, and incubated for 10 minutes (37°C, 5% CO2). The procedure was repeated for the second gel when needed.

Appropriate differentiation medium was injected into the dedicated channels. Chondrogenic medium (CHM) was constituted by DMEM (Sigma–Aldrich) containing, 4.5 mg ml^−1^ D-glucose, 0.1 mM non-essential amino acids, 1 mM sodium pyruvate, 100 mM HEPES buffer,100 U ml^−1^ penicillin, 100 μg ml^−1^ streptomycin, 0.29 mg ml^−1^ l-glutamine, 1.0 mg/ml insulin, 0.55 mg/ml human transferrin, 0.5 μg/ml sodium selenite, 50 mg/ml bovine serum albumin, 470 μg/ml linoleic acid, and 1.25 % of human serum albumin and supplemented with 0.1 mM ascorbic acid 2-phosphate, 10^-4^ mM dexamethasone and 10 ng ml^-1^ TGF-β3^98^. Osteochondral medium (OCM) was obtained by addition of beta-Glycerophosphate 10 mM (β-Gly, Sigma Aldrich), previously demonstrated to induce mineralization^59^, to the CHM.

Culture medium was changed every 2 days. Samples were cultured in static regimen for 14 days, previously demonstrated to be sufficient for achievement of cartilage on chip constructs^11^, before application of cyclic loading or collection for RT-qPCR, immunofluorescence imaging, or IT-AFM analyses. MSCs based constructs cultured in OCM were also harvested after 7 days to assess the evolution of constructs gene expression profile in time. Brightfield images of the constructs were acquired throughout the culture period.

### Application of mechanical compression

After 14 days of maturation, constructs were subjected to mechanical loading for further 7 days. Loading was applied following a previously tested regimen demonstrated to induce OA traits in cartilage on-chip tissues subjected to hyperphysiological compression^11^. Briefly, two 2-hours stimulation cycles (1Hz, 50% duty cycle) were applicated each day with a four hours stop period in-between. A pressure of 0.4 Atm, demonstrated to be sufficient for the actuation membrane to reach the bottom surface of the pillars in the bottom compartment (Supplementary materials, Supplementary Fig. 4) was adopted during cyclical loading. Mechanical stimulation was achieved connecting the devices to a pressure regulator (Comnhas), linked to a custom made electropneumatic controller able to apply the described loading regimen^11^. External connections were realized through tygon tubing (internal diameter = 0.5 mm, Qosina, Ny, USA). Culture medium was changed every second day and collected for analysis. Notably, dexamethasone, that was proved to have anti-inflammatory effects^11^, was removed from the culture medium during the mechanical stimulation period. At the end of the stimulation period, constructs were collected for analyses, possibly after construct digestion and cellular populations sorting as described below. Brightfield mages of the constructs were acquired throughout the culture period.

### Transduction of green fluorescent protein (GFP) in chondrocytes

GFP+ hACs were obtained through lentiviral transduction adopting a previously implemented protocol demonstrated not to affect chondrocytes differentiation capacity^135^. Briefly, first passage hACs were seeded in Petri dishes (Sigma Aldrich; 60 mm x 15 mm polystyrene Petri dishes) at a density of 10^4^ cells cm^-2^ and transduced the following day with 2.8 x 10^6^ TU of GFP lentivirus (Multiplicity of Infection, MOI, of 5) in presence of 5 μg ml^-1^ of protamine sulphate. Cells were detached 3 days post transduction, and GFP+ cells selected through FACS (BD FACSAria III Cell sorter). GFP+ hACs were subsequently re-passaged and used at passage 3.

### Sorting of GFP+ chondrocytes

To determine the effect of the compression in the two compartments, GFP transduced chondrocytes (i.e. GFP+ hACs) were injected in one of the compartments and non-labelled hACs in the remaining one (e.g. GFP+ hACs in the top compartment and GFP-in the bottom compartment, or *vice versa*). The LDG configuration (Extended Data Fig. 1) was adopted in this phase. Constructs were cultured statically for 2 weeks and subjected to mechanical loading for 7 days as previously described. At the end of the culture period, constructs were digested enzymatically. Briefly, constructs were extracted from devices and incubated on an orbital shaker for 1 hour in a solution of 0.15% type II collagenase (Worthington Biochemical Corporation) in DMEM, 300 µl per construct. Cells were then centrifuged, rinsed in a solution of 1mM EDTA and 2% FBS in PBS, incubated for five minutes in 0.05% trypsin/0.53 mM EDTA (37°C 5% CO2), washed twice in 1mM EDTA-2% FBS in PBS and held on ice prior to sorting. GFP positive and negative cellular populations were sorted (BD FACSAria III Cell sorter, Supplementary Fig. 7a) and analysed separately. RT-qPCR for EGFP was performed on the sorted populations and on single GFP+ and GFP-starting populations as controls to assess the efficiency of the process (Supplementary Fig. 7b). Sorted populations were also replated in 2D (IBIDI μ/slide 8 well) and left to adhere for 48 hours in complete medium (using non sorted population as controls) to directly visualize the population purity after FACS. After 48 hours, samples were rinsed with PBS and fixed with a solution of 4% formalin for 24 hours. Populations purity after sorting was confirmed through immunofluorescent imaging (Supplementary Fig. 7c).

Similar procedures were performed seeding bmMSCs in the top compartment and GFP+ hACs in the bottom compartments of VBV devices to determine the effect of the co-culture on the single populations and to assess the response to loading of the two layers of the osteochondral constructs. After 14 days of static maturation, or 14 days of static maturation and 7 days of cyclical loading, GFP+ hACs and bmMSCs populations were sorted and cells analysed separately through RT-qPCR.

### Determination of subchondral layer’s mechanical properties effects on chondrocytes response to loading

The effect of introducing local alterations in the mechanical properties of the subchondral layer during compression was investigated. NDG devices were adopted in this phase. hACs were loaded in an enzymatically crosslinkable and degradable PEG based hydrogel formulation^24^ as described above and injected in the bottom compartment of the device. The top compartment was injected with hydrogels loaded with Polystirene beads (diameter 10 um, Sigma-Aldrich) with volumetric fractions of either 0.1% or 5%. Hydrogels mechanical properties were assessed by IT-AFM as described below. Constructs were cultured statically for 14 days in chondrogenic medium before application of the above-described loading regimen for 7 further days. During the stimulation period the culture medium, which was changed every second day, was collected for analyses biochemical. At the end of the culture, constructs were collected for RT-qPCR.

### Gene expression analysis

Total RNA extraction by TRIzol (Sigma), complementary DNA synthesis, and RT-qPCR were performed according to standard protocols (7300 AB; Applied Biosystems). The following gene of interests were quantified (Applied biosystems): FOS (Hs99999140_m1), PTGS2 (Hs00153133_m1), C-JUN (Hs01103582_s1), MMP13 (Hs00233992_m1), IL8 (Hs00174103_m1), COL2A1 (Hs00264051_m1), COL1A1 (Hs00164004_m1), ACAN (Hs00153936_m1), PRG4 (Hs00981633_m1), FRZB (Hs00173503_m1), DKK1 (Hs00183740_m1), GREM1 (Hs01879841_s1), GDF5 (Hs00167060_m1), GFP (Mr00660654_cn), COL10A1 (Hs00166657_m1), IHH (Hs01081800_m1), BMP2 (Hs00154192_m1), BSP2 (Hs00173720_m1), ALP (Hs01029144_m1), OCN (Hs01587814_g1), and OPN (Hs00959010_m1). Glyceraldehyde 3 phosphate dehydrogenase (GAPDH) was used as housekeeping gene (Hs02758991_g1).

### Biochemical analyses

The concentration of biochemical factors released in the supernatant was measured. Supernatants were collected at day 16, 18, 20, and 21 during the mechanical stimulation phase, and pooled together to measure solutes accumulation during the whole stimulation period. Culture medium from the top and from the bottom compartments were collected and analysed separately to ascribe the release of solutes to one of the two compartments only. IL8 concentration was measured using the Human IL-8 ELISA Set (BD Bioscience) following the manufacturer instructions. Measurements were acquired using a configurable multimode microplate reader (synergy H1, BioTek instruments). IL-1β, TNFα, and IL6 releases were analysed by Luminex® Assay using the Human Premixed Multi-Analyte Kit (R&D Systems) according to manufacturer instructions and acquiring readings with a BioPlex MagPlex Magneti Beads system (Bio-Rad). MMP-13 production was measured using an enzyme-specific fluorescence substrate kit (SensoLyte 520 MMP-13 Assay Kit; AnaSpec), according to the manufacturer’s instructions.

### Immunofluorescence analyses

Immunofluorescence analyses were performed on cellular constructs at day 0, day 14, and day 21 directly within microscale devices. Samples were washed with PBS and fixed in 4% paraformaldehyde overnight at 4°C. Devices were subsequently disassembled removing actuation membrane and actuation chamber layers so to expose the constructs (for top view images), or cut in sections with a thickness of roughly 1 mm (for side view images). Cells were permeabilized with a solution of 0.5% (v/v) solution of Triton X (Sigma-Aldrich); unspecific binding was blocked (1 hour at room temperature) with a solution of 0.3% (v/v) Tween 20 and 3% (v/V) Goat serum in PBS. Samples were incubated overnight at 4°C with primary antibodies for aggrecan (ACAN, Abcam,ab36861, dilution 1:200), collagen type II (COL2A1,Abcam, ab185430, dilution 1:200), osteocalcin (OCN, Millipore, ab1857, dilution 1:200), and alkaline phosphatase (ALP, Abcam, ab54778, dilution 1:200) to assess tissues maturation. After washing with blocking solution (20 minutes at room temperature, repeated twice), samples were incubated (1 hour at room temperature) with antibodies labelled with Alexa Fluor 488, Alexa Fluor546, and Alexa Fluor 647 (Invitrogen, dilution 1:200) as appropriate and subsequently washed with PBS.

Staining with 4′,6-diamidino-2-phenylindole (DAPI) was used to identify the cell nuclei. Deposition of hydroxyapatite (HA) was assessed staining samples with OsteoImage mineralization assay (Lonza) according to the manufacturer’s instructions. Samples were subsequently Imaged with a Nikon, Eclipse Ti2 widefield microscope or a Nikon A1R NALA confocal microscope. GFP and RFP positive cells were imaged directly without prior stainings. Image analysis was performed using Image J and QuPath softwares.

### hACs and bmMSCs culture in 3D aggregates (i.e. pellets)

hACs and bmMSCs isolated and expanded as described above were cultured in 3D pellets for 14 days in either CHM or osteochondral medium OCM. Passage 3 cells were detached using 0.05% trypsin-0.53 mM EDTA (Gibco) and counted with Trypan blue (Thermo-Fisher). 2.5 x 10^5^ cells were adopted for each pellet. Cellular aggregation was obtained via centrifugation (1300 RPM, 3 min), in screw top microtubes (Thermo-fisher). Culture medium, 1 ml for each pellet was changed thrice a week.

### GAG and DNA quantifications

Samples were digested overnight at 56°C in 500 µl of proteinase-K solution (1 mg mL^-1^ proteinase-K in 50 mM Tris with 1 mM EDTA, 1 mM iodoacetamide, and 10 μg mL^-1^ pepstatin-A). GAG amounts were measured spectrophotometrically after reaction with dimethylmethylene blue using chondroitin sulphate as a standard^136^. DNA quantified using the CyQuant cell proliferation assay Kit (Molecular Probes, Eugene, OR) and was measured spectrophotometrically according to manufacturer’s instructions.

### Histological Staining and Immunohistochemistry

Cell pellets were washed with PBS, fixed in 4% paraformaldehyde overnight at 4 °C, and embedded in paraffin. After deparaffinization and re-hydratation sections were stained for Safranin-O (Fluka) and Alizarin red (Sigma) to assess the deposition of GAG and calcium deposits respectively.

Immunohistochemical staining, respectively for COL2A1 (MP Biomedicals 63171, 1:1000 dilution) and COL10A1 (Invitrogen 14-9771-80, 1:200 dilution) were performed with Ventana Discovery Ultra (Roche Diagnostics (Switzerland), SA) automated slide stainer. Tissue sections were deparaffinized and rehydrated. Antigens were retrieved by protease digestion (Protease 3, 760-2020, Ventana) for 20 to 44 minutes at 37°C. Primary antibody were manually applied and incubated for 1 hour at 37°C. After washing, the secondary antibody (anti-mouse polymer horseradish peroxidase (HRP), R&D Mouse IgG (VC001-025, VisUCyte)) was incubated for 1 hour at 37°C. The detection step was performed with the Ventana DISCOVERY ChromoMap DAB (760-159, Ventana) detection kit. Slides were counterstained with hematoxylin II, followed by the bluing reagent (respectively, 790-2208 and 760-2037; Ventana).

Histological and immunohistochemical sections were imaged with a using a Nikon, Eclipse Ti2 microscope with Nikon DS-Ri2 camera.

Safranin-O images were adopted for tissues histological grading through the Modified Bern Score^137^. Grading was performed as previously described^76^ through a deep learning based automated procedure using tiles of 224×224 pixels with a pixel dimension of 0.511 µm.

### Mineral deposition quantification

The effect of cyclical loading on osteochondral tissue mineralization was analysed quantifying the amount of Hydroxyapatite (HA) deposited in biphasic osteochondral constructs, and the amount of calcium salts deposited in the culture medium channels of microscale devices.

Ha deposition was detected staining constructs sections (n≥10 from n≥3 chambers considering two different hACs and two different MSCs donors) with OsteoImage (Lonza) as detailed above. DAPI was used to counterstain nuclei. Images were quantified using QuPath^89^. Briefly, areas corresponding to the subchondral compartment (comprehensive of top compartment and VBV necking area) and to the cartilaginous compartment were traced manually on each image. The total number of cells in each compartment was computed using QuPath cell detection tool. The total HA positive area for each of the tissues was computed though a threshold-based pixel classifier. Subchondral compartment and cartilage compartment were analysed separately. For each compartment the HA positive area was normalized for the number of cells detected in the area.

The amount of calcium salts deposited in the culture medium channels of microscale devices was quantified through an Alizarin Red staining. Briefly devices were opened peeling of actuation membrane and actuation chamber layers and cell constructs removed. Empty devices were stained with Alizarin Red completely immerging the chambers in the solution. Stained calcium salts were then dissolved in a 10% (v/v) acetic acid solution (30 minutes at room temperature with shaking at 300 rpm) which was then collected incubated at 85

°C for 10 minutes, put immediately on ice, and centrifuged at 12000 g for 15 minutes. Supernatants were analysed quantifying the at 405 nm in a Synergy H1 Hybrid Multi-Mode Reader (BioTek Instruments,Winooski, VT, USA).

### Statistical Analysis

Results of FE analyses, GAG/DNA qunatifications, and biochemical analyses were presented ad mean ± standard deviation. RT-qPCR results were presented as mean + standard deviation. Single data were plotted to account for non-normal distributions. Population normality was verified using Shapiro-Wilk and Kolmogorov-Smirnov tests. Non-Paired double comparisons were performed using two tailed t-test for normal populations and Mann-Whitney test for non-Gaussian ones. Paired double comparison were performed using paired t-test and Wilcoxon test respectively. Multiple comparisons were performed using ordinary one-way analysis of variance (ANOVA). Ordinary one-way ANOVA with Tukey’s multiple comparison tests were adopted for normal distributions, Kruskal-Wallis test with Dunn’s multiple comparison tests for non-normal distributions.

### Single-cell RNA-seq

Single cell RNA-seq (scRNA-seq) was performed on hACs after 21 days of culture (i.e. 14 days of static maturation and 7 days of mechanical stimulation, with constructs cultured statically for 21 days used as controls). Single culture hACs and OCU-on-chip constructs were retrieved from microfluidic devices and enzymatically digested on a shaker for 30-45 minutes as described above. Cells were then transferred in a solution of 10% FBS in PBS and live-dead sorted using DAPI. Enzymatic digestion time and sorting strategies led to a preferential selection of hACs over bmMSCs. For each donor and for each condition cells from 4 cellular constructs were pooled together. A total of 4 donors was considered. Biological replicates originating from different donors and corresponding to the same condition were then pulled together into one sample before sequencing preparation.

Single-cell capture and cDNA and library preparation were performed at the Genomics Facility of the University of Basel

Cells from each sample were loaded on a Chromium Single Cell Controller (10x Genomics, Pleasanton, CA, USA). Single-cell capture, and cDNA and library preparation were performed with a Chromium Next GEM Single-Cell 3’ Reagent Kit v3.1 (Dual Index) with Feature Barcode technology for Cell Surface Protein and Cell Multiplexing from 10x Genomics (User Guide CG000390, Rev B) according to the manufacturer’s instructions. Pair end sequencing (with read setup 28-10-10-90) was performed on 1 Lane of a S4 Flowcell with an Illumina NovaSeq 6000 according to 10x specifications.

### Single-cell data analysis

cDNA reads were aligned to the ‘hg38’ genome using Ensembl 104 gene models employing the STARsolo tool (v2.7.10a) with default parameter values exception made for the following ones: soloBarcodeReadLength=0, clipAdapterType=CellRanger4, outFilterType=BySJout, outFilterMultimapNmax=10, outSAMmultNmax=1, soloType=CB_UMI_Simple, outFilterScoreMin=30, soloCBmatchWLtype=1MM_multi_Nbase_pseudocounts, soloUMIfiltering=MultiGeneUMI_CR, soloUMIdedup=1MM_CR, soloCellFilter=None, soloMultiMappers=EM, soloCBstart=1, soloCBlen=16, soloUMIstart=17, soloUMIlen=12.

Merged samples were demultiplexed based on present single nucleotide polymorphisms (SNPs) using the cellsnp-lite^138^ and vireo^139^ tools. BAM files with mapped reads and the SNP database HapMap 3.3 (https://www.sanger.ac.uk/resources/downloads/human/hapmap3.html) were used as input. Resultingly, cells were either assigned to a particular genotype of individual donors or not assigned to any genotype.

Further analysis steps were performed using R (v4.2.0). Cells were clustered based on detected SNPs by building a shared nearest-neighbour (SNN) graph (k=10)^140^ and using the ‘cluster_louvain’ method from the R igraph package^141^. The majority of clusters was composed of cells of the same genotype. A cluster of cells that was characterized by mixed genotypes was assigned as composed of low-quality cells; a cluster of cells with no genotype was assigned as composed of doublets. Cells from these clusters as well as residual bmMSCs cells from bmMSCs donors used for OCU. on-chip constructs were removed from further analyses. The final data set consisted of 24579 cells.

Multiple Bioconductor (v3.15) packages including DropletUtils (v1.16.0), scDblFinder (v1.10.0), scran (v1.24.1), scater (v1.24.0), scuttle (1.6.2) and batchelor (v1.12.3) were applied for the subsequent gene expression analyses of remaining cells. Analyses were mostly performed following the steps of the workflow presented at https://bioconductor.org/books/release/OSCA/. Raw counts were normalized by deconvolution^142^. The function ‘computeDoubletDensity’ did not point at any group of cells, which could be potential doublets, confirming the efficiency of removing doublets while demultiplexing samples.

The most variable genes were derived by applying the function ‘getTopHVGs’ with the following parameters values: fdr.threshold=0.01 and var.threshold=0.1. The expression of the most variable genes was then used to perform principal component analysis. Principal components were corrected by removing the batch effect based on the genotype (i.e. the effect of individual donors) (Extended Data Fig. 5a). Afterwards the ‘runUMAP’ function from the Scater package was applied (with default parameters) to the corrected principal components in order to perform UMAP dimensionality reduction, adopted to visualize single cells on two dimensions (Fig. 6b).

Cells were then re-clustered based on the corrected principal components coordinated by building a shared nearest-neighbour (SNN) graph (k=100) and using the ‘cluster_louvain’ method (Fig. 6c, d). The ‘scoreMarkers’ function of the scran package was applied to find clusters characteristic marker genes. The standardized log-fold change across all pairwise comparisons ‘mean.logFC.cohen’>1 was used as the significance threshold. Marker genes common for two or more clusters were removed from the analysis.

The data set was subjected to cell-type annotations using the Bioconductor package SingleR (v1.10.0). A publicly available annotated scRNA-Seq data set^8^ was used as the reference.

Differential expression analyses in static and mechanically stimulated samples, both considering single culture hACs, and hACs that were co-cultured in OCU-on-chip constructs, were performed aggregating single cells into *in silico* bulk samples^143^. Specifically, pseudo-bulk samples were created by summing up counts per gene across all cells belonging to a particular cluster, condition, and replicate. The function ‘filterByExpr’ excluded lowly expressed genes from the analysis. Combinations containing at least 20 cells underwent analysis with edgeR (3.38.4). Genes were considered as differentially expressed if they had a false discovery rate (FDR)<0.05. A gene ontology (GO) analysis of differentially expressed genes was performed using STRING Database (https://string-db.org/)^95^, Version 11.5.

Gene set enrichment analysis (GSEA, v4.2.3) was applied to differentiate pathways affected by mechanical loading in hACs form single culturesa and hACs from co-cultures in OCU-on-chip constructs^144^. Reported pathways were based on the KEGG database (http://www.kegg.jp). For each contrast, expressed genes were ranked by their log-fold changes as reported by edgeR. For each pathway and each comparison, the FDR accounting for the significance of the enrichment was calculated. A pathway was reported if the corresponding FDR was less than 0.1.

**Reporting summary**. Further information on research design is available in the Nature Research Reporting Summary linked to this article.

**Data availability.** The authors declare that all data supporting the findings of this study are available within the paper and its Supplementary Information. The data that support the findings of this study are available from the corresponding author upon reasonable request.

## Supporting information

Supplementary file 1 Clusters markers

Supplementary file 2 PCA Analysis

Supplementary File 3 DE genes

## Acknowledgements

The authors are grateful to Dr. Alessandro Caimi for his inputs on FE simulations and to Dr. Lorenzo Raeli, Dr. Jelena Markovic-Djuric, and Stella Stevanova for their support with cell sorting. The authors are also grateful to Monica S. Schönenberger for her help with IT-AFM analyses on beads laden hydrogels. Device manufacturing was partially performed at PoliFAB, the micro-and nanofabrication facility of Politecnico di Milano. scRNA-seq was performed at the Genomics Facility Basel in the D-BSSE of the ETH of Zurich in Basel. The authors are specifically grateful to Dr. Christian Beisel and to Mirjam Judith Feldkamp. scRNA-seq data were analysed by the Bioinformatics Core Facility, Department of Biomedicine, University of Basel. This work was partially funded by the Swiss National Science Foundation (number 310030_175660/1, to AB) and by Cariplo Foundation (grant number #2018-0551, to MR)

## Author contributions

AM conceived the VBV design. AM and MR conceived the device. AM realized and functionally validated the device. AM and PO planned biological experiments. AM performed biological experiments and analyses. AM conceived and performed FE simulations. AM, and POE performed IT-AFM. POE and ML performed IT-AFM analyses. ME and LK fabricated and provided the hydrogel. AB, PO and MR conceived the project. AM, PO, IM, MR and AB wrote the manuscript. AM realized the manuscript figures. RI and AB performed single cell RNA-seq analyses. AM and AB discussed scRNA-seq data. All authors discussed the results, commented on the manuscript, and contributed to its final version. AB and MR contributed equally to this work.

## Additional information

Additional information concerning scRNA-seq analyses can be found in Supplementary Files 1 – 3. The gene expression profiles of the samples have been deposited in NCBI’s Gene Expression Omnibus ^145^and are accessible through the GEO Series accession number GSE237786.

