## Extended data for "Modelling Osteoarthritis pathogenesis through Mechanical Loading in an Osteochondral Unit-on-Chip"

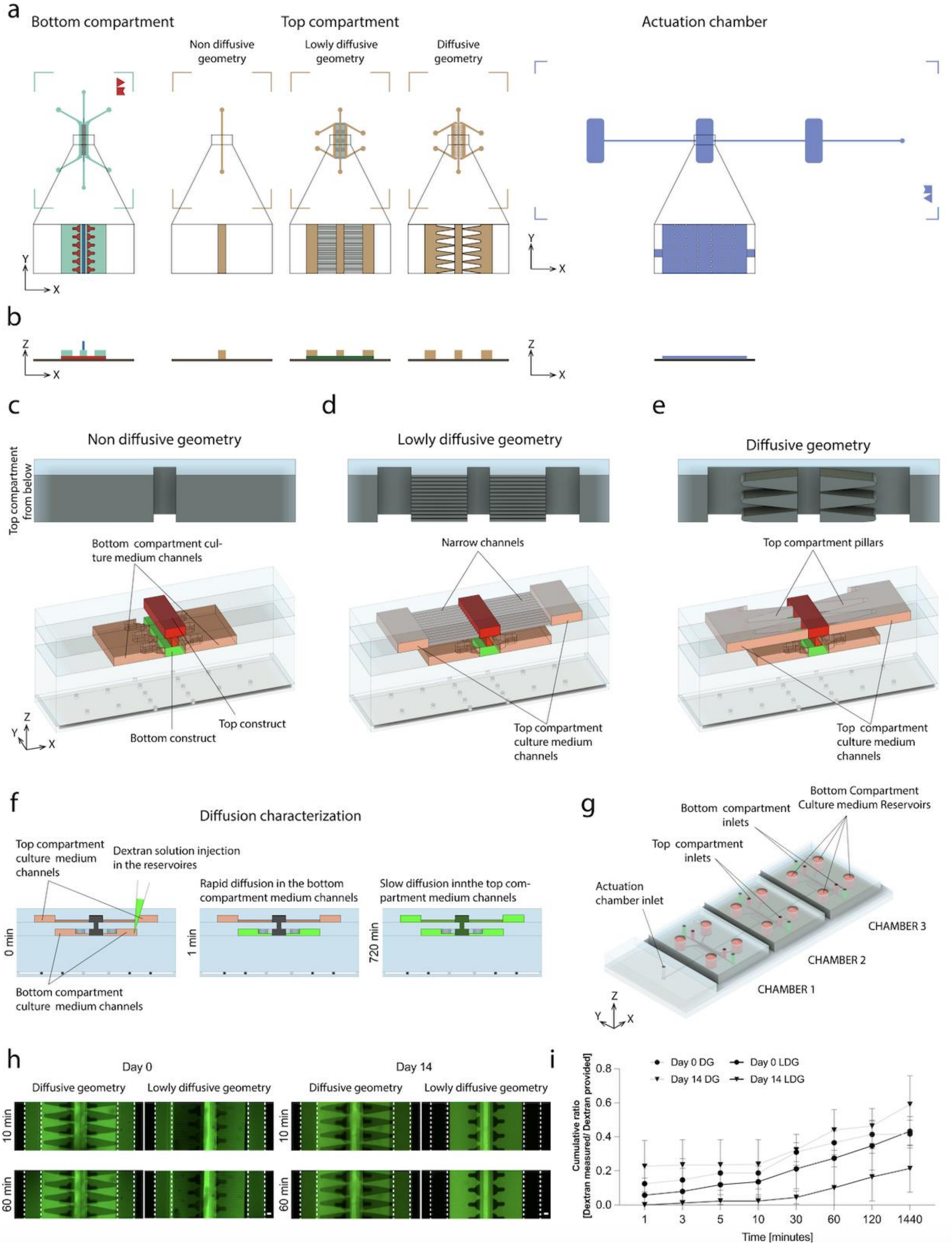


**Extended Data Fig. 1 Geometries of the alternative versions of the device and diffusion kinetic characterization. a, b** Layouts of the master molds used to produce the device; all three versions of the top compartment are represented. The different layers used to obtain the final master molds geometries through multi-layer photolithography are indicated by the colors. The bottom compartment was realized through three layers: gap layer (height 43 μm) indicated in red, pillars layer (height 100 μm) in aqua green, and necking geometry layer (height 150 μm) in blue. Top compartment: Non diffusive geometry and Diffusive geometry have a single layer (height 150 μm) indicated in beige; the Lowly diffusive geometry is constituted by the narrow channels layer (height 50 μm), indicated in dark green, and the top layer (height 100 μm), depicted in beige. The Actuation chamber was realized through a single layer (height 50 μm), portrayed in purple. Six rows of circular pillars (diameter 28µm) were incorporated in the actuation chamber to prevent buckling. **c-e,** Schematics detailing the inner structure of the device in its three versions. In the top area are represented the three geometries of the top compartment as seen from below. The Non-diffusive geometry is constituted by a simple channel; in the Lowly diffusive geometry the central channel is connected with lateral culture medium channels by narrow channels (10 µm width, 50 µm height, 50 µm separation between the channels); in the Diffusive geometry channels are separated by triangular shaped pillar structures (300 µm large at the channel level) with a distance of 30 µm between them. **f,** Schematic of the experimental setup used to determine the diffusion kinetic between bottom and top compartments. Schemes refer to the LDG and depict a section of the device at three different time points during the experiment. The device bottom channel's culture medium reservoirs were filled with a dextran solution (20 kDa, 1 mg $ml^{-1}$) which rapidly diffuse in the culture medium channels, while it takes longer to diffuse in the top compartment's culture medium channels depending on the geometry of the top compartment layer. The dextran concentration in the top compartment culture medium reservoirs was measured at selected time points. The diffusion process was also observed directly though time lapse imaging. The experiment was performed comparing only LDG and DG given that the NDG does not feature the top compartment culture medium channels. hACs were laden in 2% PEG based hydrogels and seeded in both top and bottom compartments' central channels. The diffusion assay was performed either on the same day of hACs seeding (i.e. day 0) or after static maturation in chondrogenic medium (see methods section) for 14 days (i.e. day 14). hACs maturation causes them to deposit extracellular matrix (ECM) thus diminishing solutes diffusivity through the constructs. **g,** Schematization of the PDMS device with the NDG of the top compartment. Each platform is constituted by three culture chambers. Differently from the DG configuration represented in Fig. 1 b, the NDG does not require the presence of culture medium reservoirs for the top compartment. **h**, Representative fluorescence images of FITC dextran diffusion after 10 mins and 60 mins from injection in the bottom compartment culture medium reservoirs (n= 3 biologically independent samples per condition and per time point). Top compartment culture medium channels are highlighted by dotted lines. Scale bar 100 µm. **i**, 20 KDa FITC dextran diffusion rate quantification via measurement of FITC fluorescent intensity in the top compartment culture medium reservoirs. Concentration in time was expressed as the cumulative ratio between the dextran injected in the bottom compartment and the quantity measured in the top compartment at selected time points (which equals 0.5 at the equilibrium) (n= 3 biologically independent samples per condition and per time point). Adoption of the DG permitted fast diffusion of the molecule and resulted in cumulative ratios, after 60 mins, of 0.37 ± 0.07 and 0.44 ± 0.12, respectively, at day 0 and at day 14, with no difference introduced by constructs maturation. A slower diffusion was registered in the LDG when coupled with mature constructs. Cumulative ratios after 60 mins resulted of 0.27 ± 0.17 and 0.1 ± 0.12 for day 0 and day 14, respectively.

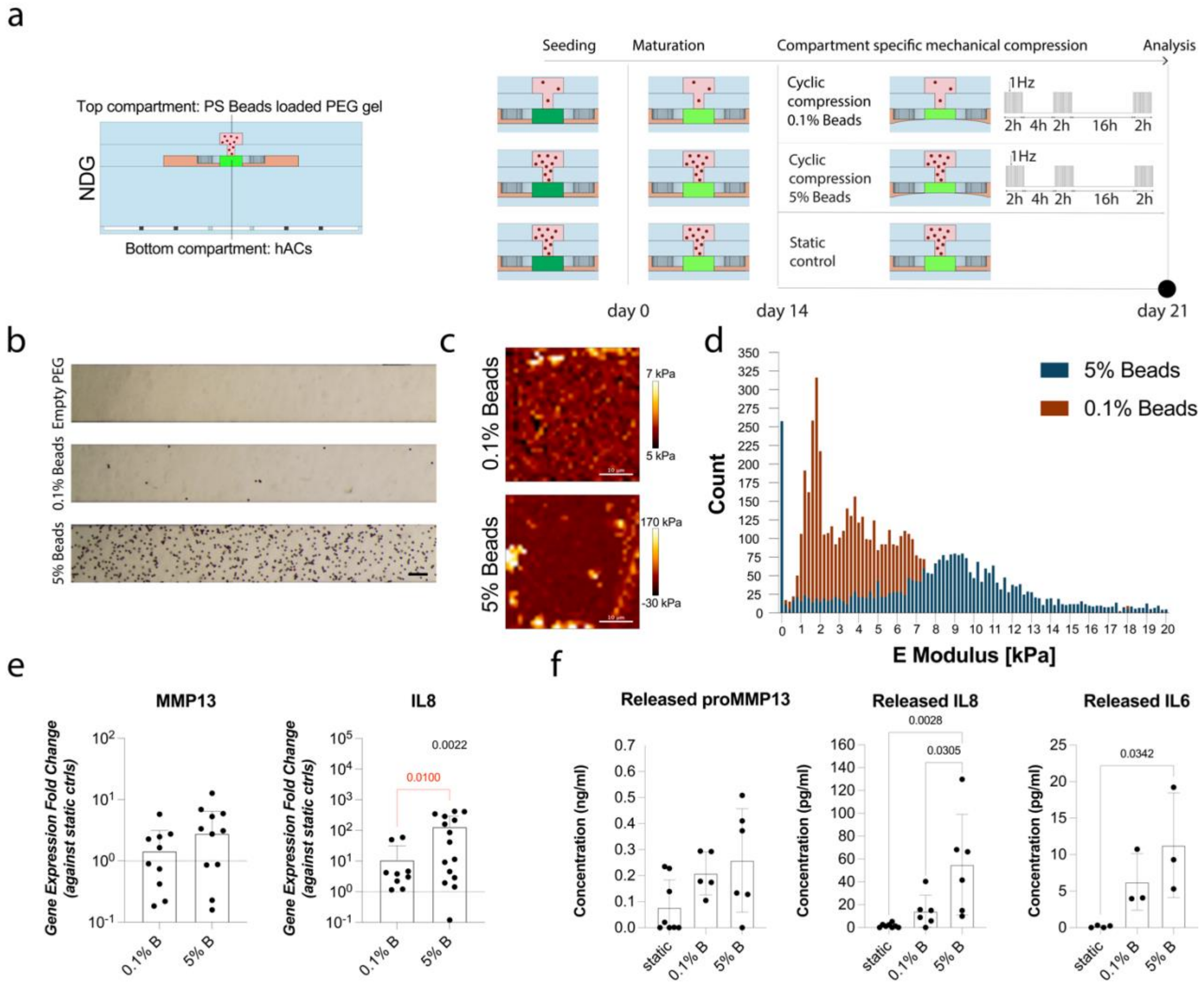

**Extended Data Fig. 2 Subchondral layer's mechanical properties affect chondrocytes response to HPC**. **a,** Experimental setup. Polystyrene beads (either 0.1% or 5% in volume) loaded PEG hydrogels were seeded in the top compartment (NDG configuration), hACs in the bottom compartment. Constructs were cultured statically for 14 days before application of the loading regimen indicated in the panel for further 7 days. Static devices were used as controls. **b,** Brightfield images of the top compartment central channel (NDG configuration) seeded with empty PEG or with PEG hydrogels loaded with different volumetric fractions of polystyrene beads. Scale bar, 100 µm. **c,** Examples of IT-AFM indentation maps. E modulus values are color coded. Scale bar, 10 µm**. d,** Frequency histogram of gels E modulus as assessed through IT-AFM. **e, f,** An increase in the number of stiff inhomogeneities in the subchondral layer enhances chondrocytes response to pathological HPC. **e,** Gene expression quantification through RT-qPCR. (n≥9 biologically independent samples from n=3 donors). Statistical significance was determined by Mann-Whitney test to compare two populations (red), or by Kruskal-Wallis test with Dunn's test for multiple comparisons with respect to static controls (indicated by horizontal black lines)). All genes expression was referred to GAPDH expression and values were normalized for the expression of each donor of static controls. Values are reported as mean + s.d. **f,** Cytokines and degradative enzymes release in the culture medium following HPC. proMMP13 and IL8 concentrations were determined by ELISA (n=6 biologically independent samples from n=2 donors); IL6 concentration by Luminex analysis (n≥3 biologically independent samples from one donor). Statistical significance was determined by one-way ANOVA with Tukey's multiple comparison test for normal populations and by Kruskal-Wallis test with Dunn's multiple comparison test for non-normal distributions. Values are reported as mean ± s.d. For all graphs, populations' normality was assumed if both Shapiro-Wilk and Kolmogorov-Smirnov tests resulted positive. (Adjusted) P values < 0.05 are reported on the graph.

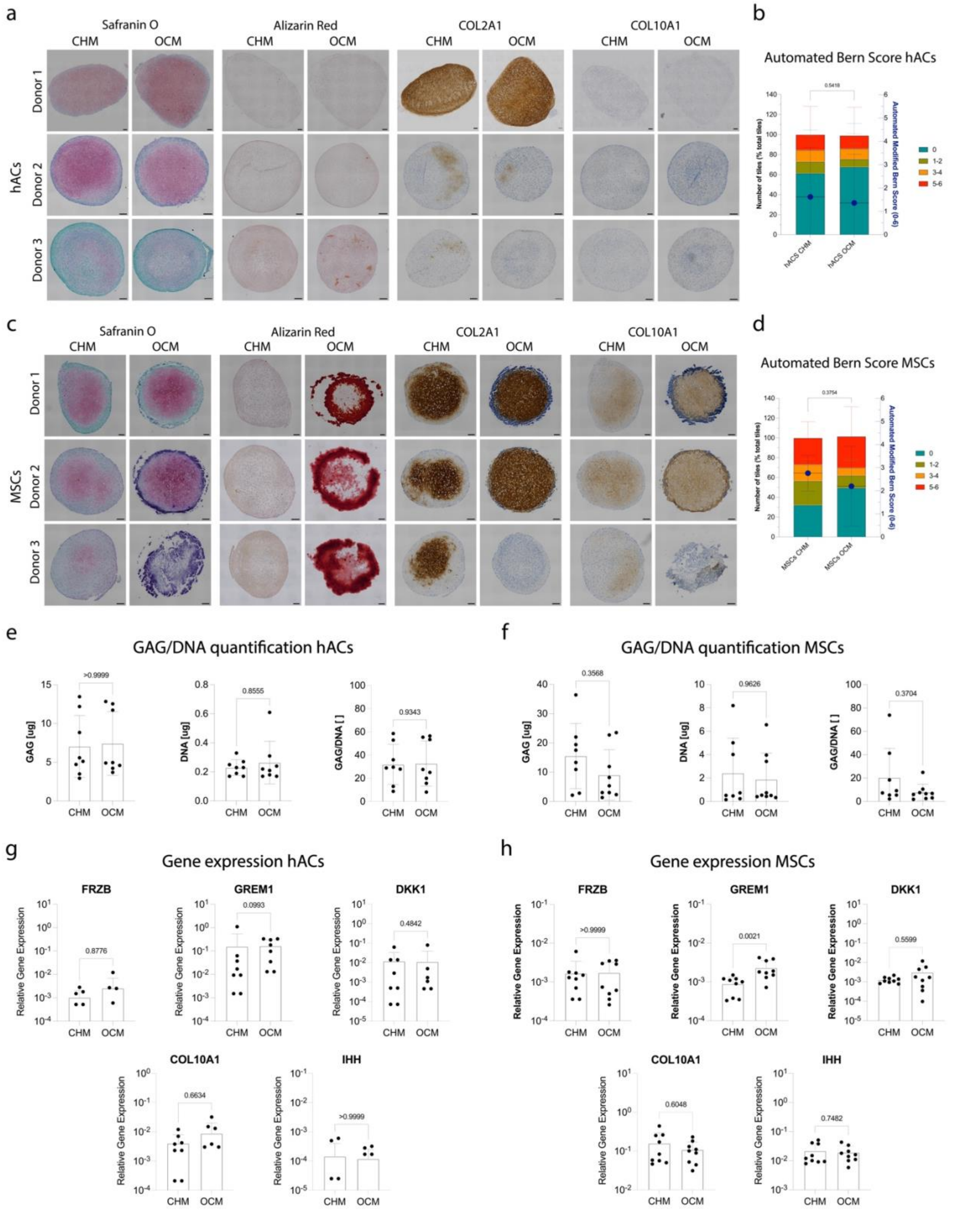


**Extended Data Fig. 3. Beta-Glycerophosphate induces MSCs mineralization without affecting hACs chondrogenicity. a,** Histological stainings and immunohistochemistry images of hACs aggregates (i.e. pellet) after 14 days of culture in CHM or OCM. Safranin-O / fast-green staining was used to indicate GAGs deposition, Alizarin Red to stain calcium deposits. Immunohistochemistry stainings of COL2A1 and COL10A1 were used as markers of hyaline cartilage and hypertrophy respectively. Passing from CHM to OCM had no impact on hACs phenotype (n≥2 biologically independent samples from n=6 hACs donors were evaluated). Scale bar 100 µm. **b,** Grading of hACs pellets Safranin O / fast-green staining images based on the Automated Modified Bern score performed as described in [1] where images are divided in squared tiles of a given dimension, each tile is scored from 0 to 6 (according to positivity for Safranin-O staining and cell morphology and an average score computed). Automated scoring was performed adopting tiles of 224x224 pixels with a pixel dimension of 0.511 µm. Results are reported as mean ± s.d. Tiles percentages are reported in colors according to the score, average values are indicated by the blue dots (n≥2 biologically independent samples from n=6 hACs donors were evaluated). Statistical significance was determined by Mann-Whitney test. **c,** Histological stainings and immunohistochemistry images of MSCs pellets after 14 days of culture in CHM or OCM. (n≥2 biologically independent samples from n=5 MSCs donors were evaluated) Scale bar 100 µm. Passing to the OCM formulation induces MSCs to deposit a Calcium Phosphate rich extracellular matrix. Grading of MSCs pellets Safranin O / fast-green staining as reported in **b**. Results are reported as mean ± s.d. Tiles percentages are reported in colors according to the score, average values are indicated by the blue dots. (n≥2 biologically independent samples from n=5 MSCs donors were evaluated). Statistical significance was determined by Mann-Whitney test. **e, f,** GAG quantifications in hACs (**e**) and MSCs (**f**) pellets after 14 days of culture in CHM or OCM indicating no significant differences between the two experimental conditions (n≥1 biologically independent sample from n=4 donors were evaluated for each cell type and experimental condition). Statistical significance was determined by Mann-Whitney test. Results are reported as mean ± s.d. **g, h,** Gene expression quantification through RT-qPCR of hACs (**g**) and MSCs (**h**) pellets after 14 days of culture in CHM or OCM (n=3 biologically independent sample from n=3 donors). Statistical significance was determined by Two-tailed Student's t-test for normal distributions and by Mann-Whitney test for non-Gaussian ones. All genes expression was referred to GAPDH expression. Gene expression values are reported as mean + s.d. For all graphs, populations' normality was assumed if both Shapiro-Wilk and Kolmogorov-Smirnov tests resulted positive. (Adjusted) P values are reported on the graph. The only statistically significant difference (P< 0.05) being constituted by GREM1 gene expression in MSCs gene expression analyses confirmed that OCM does not affect hACs chondrogenic potential in pellet culture.

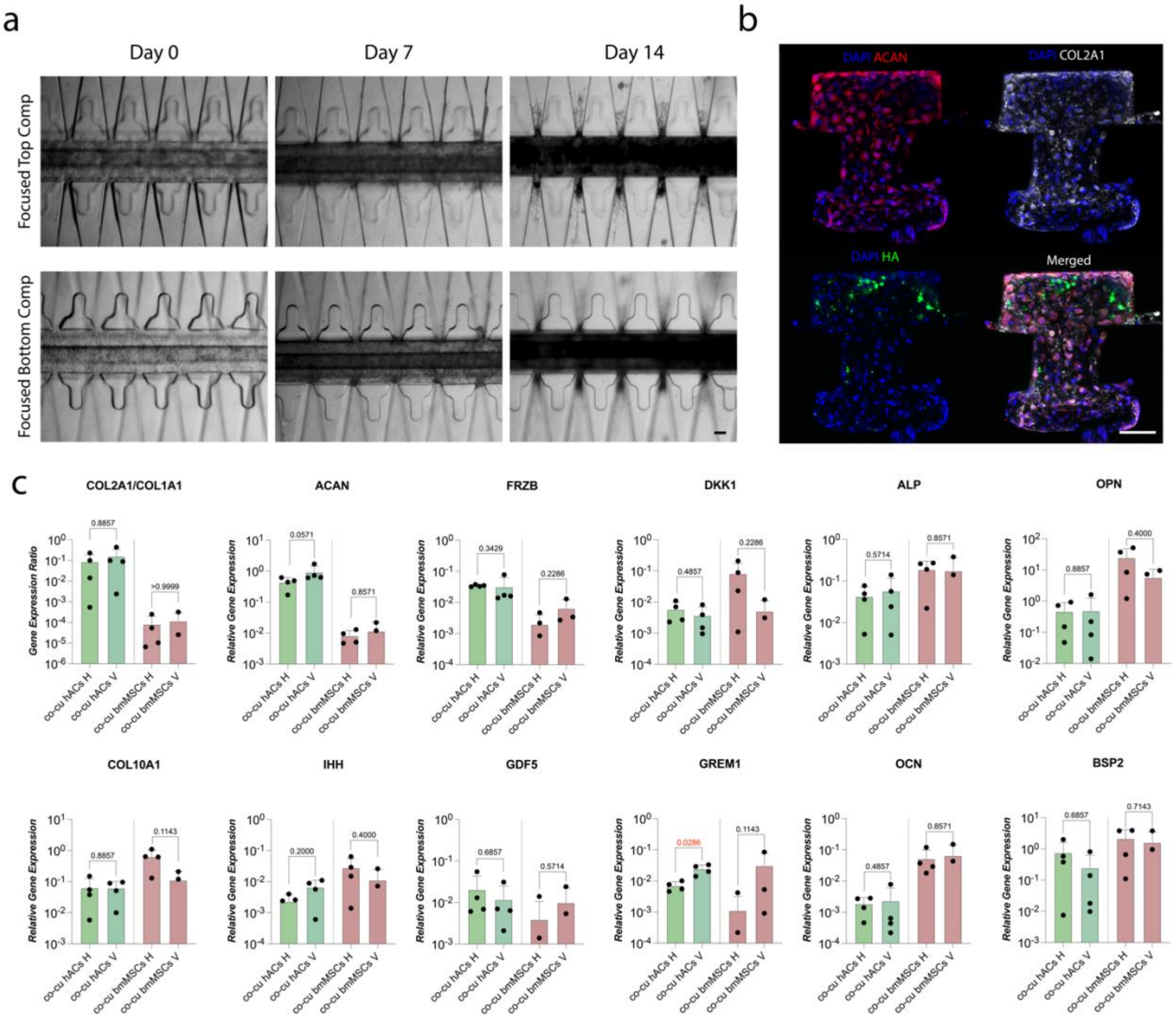


**Extended Data Fig. 4. Osteochondral construct can be achieved in VBV devices. a,** Brightfield pictures of osteochondral tissues in time (n≥3 independent samples from n≥3 donors/experiments were considered). The DG was adopted for the top compartment layer. Constructs appear darker in time during the static maturation phase, consistently with the deposition of a mineralized matrix as observed in osteochondral tissues with a horizontal disposition. Scale bar, 100 µm. **b**, Immunofluorescence images of osteochondral constructs after 14 days of static maturation. hACs were seeded in the bottom compartment, MSCs in the top compartment (n≥.2 biologically independent samples from n≥3 donors/experiments were considered in analyses). Hyaline cartilage markers ACAN and COL2A1 are represented in red and white respectively. HA is represented in green. Both constructs were positive for cartilage markers while HA was confined to the subchondral construct. As observed in horizontal devices HA positive cells appeared also in the hACs compartment. In all images DAPI (represented in blue) was used as nuclear counterstaining. Scale bar, 100 µm. **c**, Comparison of hACs and MSCs gene expression when co-cultured with a horizontal (H) or a vertical (V) disposition. Constructs were co-cultured for 14 days in static conditions, digested and sorted. GFP hACs were adopted to distinguish the two populations. Gene expression was quantified through RT-qPCR. (n=4 biologically independent samples from n=2 MSCs donors and n=1 hACs donor, i.e. the GFP expressing donor were analysed). Statistical significance was determined by Mann-Whitney test comparing only cells from the same compartment co-cultured with H and V dispositions. Populations normality was assessed though Shapiro-Wilk and Kolmogorov-Smirnov tests. P values are reported on the graph. All genes expression was referred to GAPDH expression. Values are reported as mean + s.d. No statistical differences between the two devices were detected in both populations except for GREM1 expression which increased in hACs co-cultured with the vertical disposition.

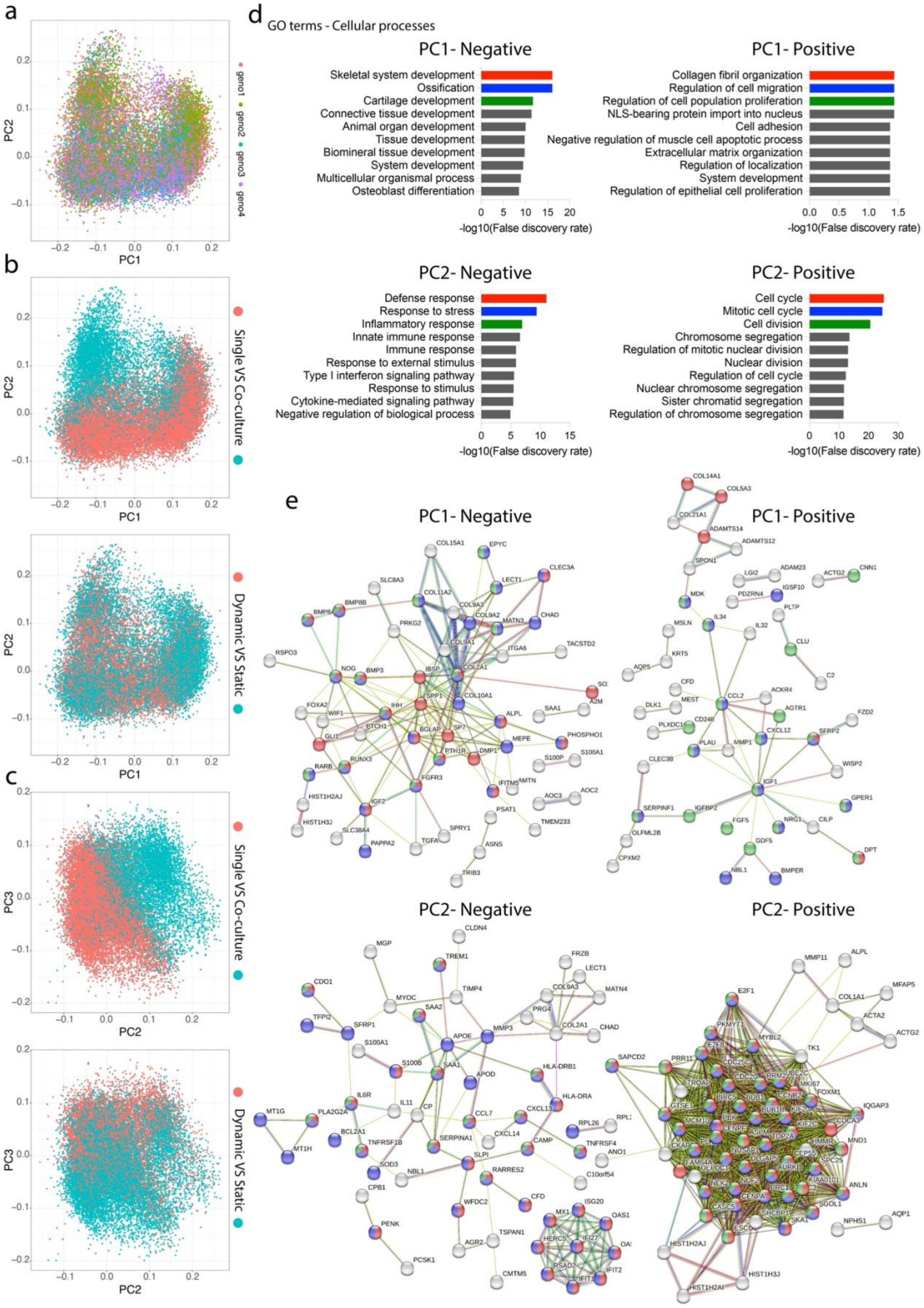
a
PC1
PC2
geno1
geno2
geno3
geno4
b
Single VS Co-culture
Dynamic VS Static
c
PC3
d
GO terms - Cellular processes
PC1- Negative
Skeletal system development
Ossification
Cartilage development
Connective tissue development
Animal organ development
Tissue development
Biomineral tissue development
System development
Multicellular organismal process
Osteoblast differentiation
-log10(False discovery rate)
PC1- Positive
Collagen fibril organization
Regulation of cell migration
Regulation of cell population proliferation
NLS-bearing protein import into nucleus
Cell adhesion
Negative regulation of muscle cell apoptotic process
Extracellular matrix organization
Regulation of localization
System development
Regulation of epithelial cell proliferation
-log10(False discovery rate)
PC2- Negative
Defense response
Response to stress
Inflammatory response
Innate immune response
Immune response
Response to external stimulus
Type I interferon signaling pathway
Response to stimulus
Cytokine-mediated signaling pathway
Negative regulation of biological process
-log10(False discovery rate)
PC2- Positive
Cell cycle
Mitotic cell cycle
Cell division
Chromosome segregation
Regulation of mitotic nuclear division
Nuclear division
Regulation of cell cycle
Nuclear chromosome segregation
Sister chromatid segregation
Regulation of chromosome segregation
-log10(False discovery rate)
e
PC1- Negative
PC1- Positive
PC2- Negative
PC2- Positive

**Extended Data Fig. 5. Principal Component Analysis (PCA) reveals the effect of compression and co-culture on hACs. a**, PCA plot of single-cell transcriptomes coloured according to the cell donor. **b**, PCA plots (PC1 vs PC2) of single-cells transcriptomes coloured according to hACs stimulation state (bottom) or culture conditions (top). **c** PCA plots (PC2 vs PC3) of single-cells transcriptomes coloured according to hACs stimulation state (bottom) or culture conditions (top). **d**, Enriched GO terms (cellular processes,) obtained using the 100 most positively correlated and the 100 most negatively correlated genes along PC1 and PC2 respectively. GO enrichment analysis was performed using STRING. Graphs report the first 10 GO terms as ordered according to lowest False Discovery Rate. **e**, STRING-based protein-protein interaction network obtained considering the 100 most positively correlated and the 100 most negatively correlated genes along PC1 and PC2 respectively. Nodes are either grey or coloured according to their relation to the GO terms with the lowest false discovery rate as reported in respective bar graphs above (i.e. in panel e). Non connected network nodes are not reported in the images.

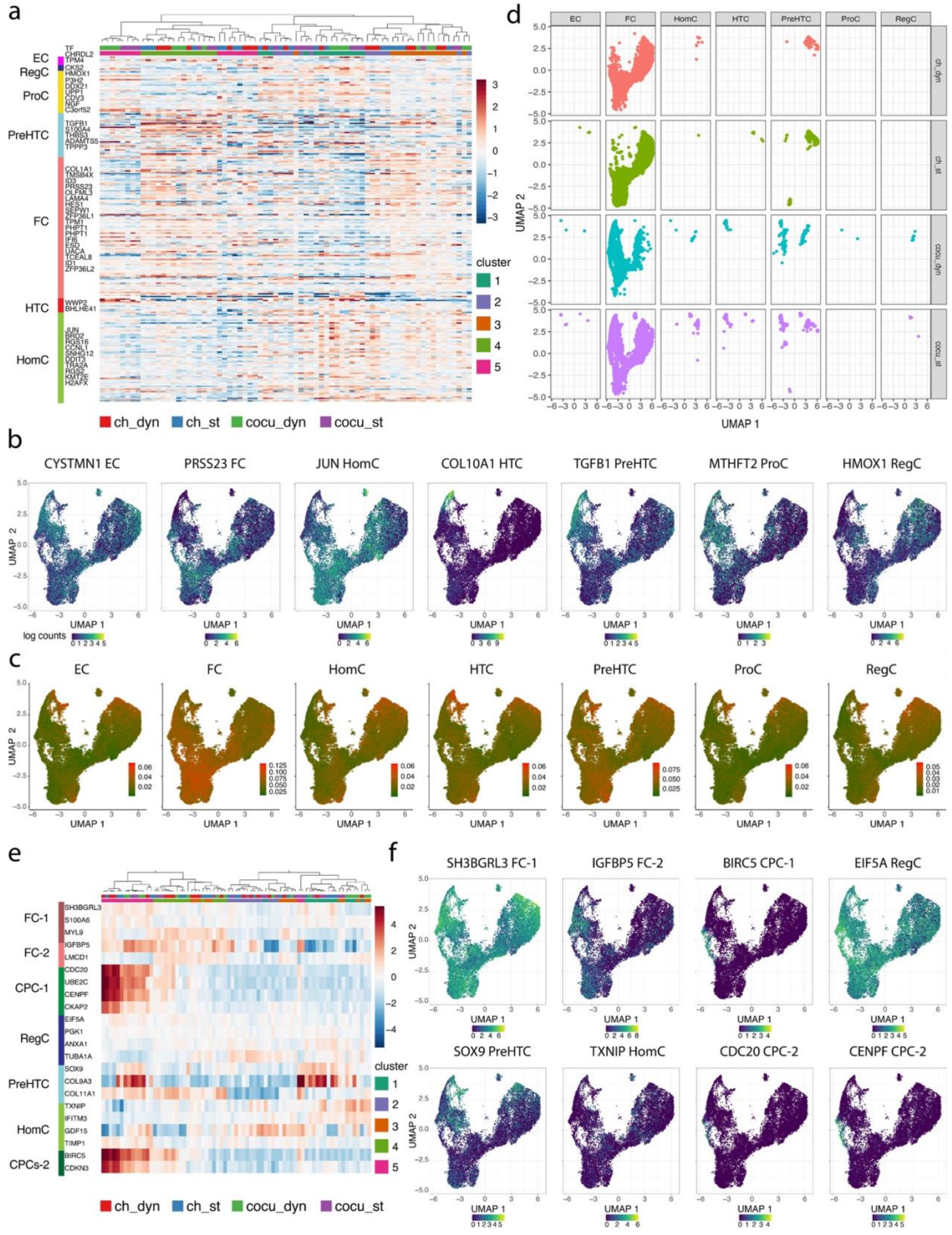


**Extended data Fig. 6 Phenotyping of hACs from cartilaginous and OCU-on-chip constructs**. **a**, Hierarchical clustering of samples based on the expression of known OA hACs subpopulations marker genes[2]. Samples culture conditions, stimulation state, and assigned cluster are represented at the top. Selected marker genes and the corresponding cell type are indicated on the left. **b**, Feature plots of selected cell types markers genes as determined from OA patients' cartilage samples[2]. **c**, UMAP representation of scRNA-seq data. Cells are coloured according to a score that defines the fitting of its assignment to a specific chondrocytes subpopulation based on the expression of known markers[2] **d**, scRNA-seq data UMAP representation according to culture conditions and divided by cell type. **e**, Hierarchical clustering of samples based on the expression of previously reported hACs subpopulations in healthy individuals[3]. Samples culture conditions, stimulation state, and assigned cluster are represented at the top. Selected marker genes and the corresponding cell type are indicated on the left. **f**, Feature plots of selected healthy chondrocytes cell type gene markers[3].

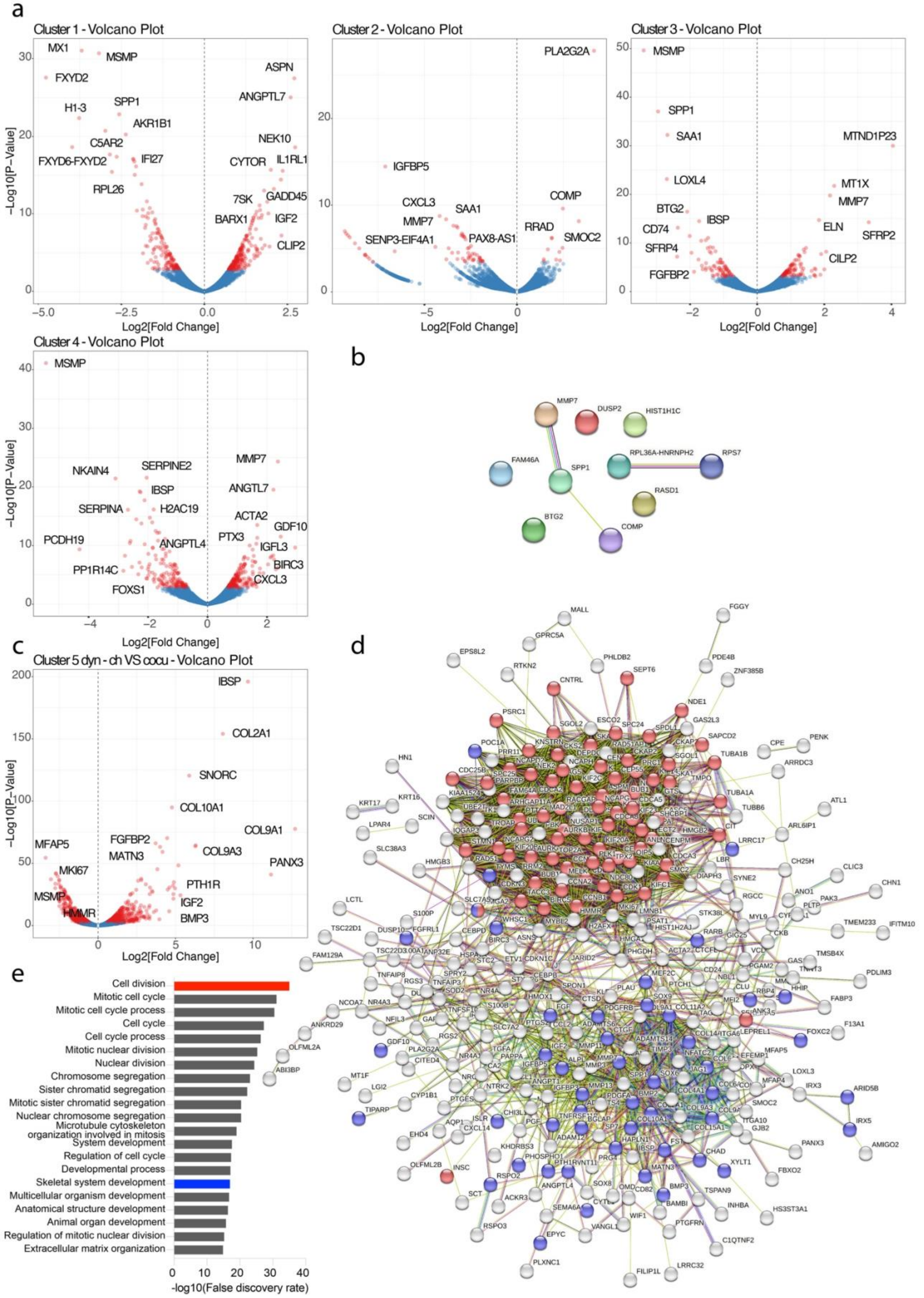

**Extended data Fig. 7 a,** Volcano plot of DE genes obtained comparing static samples and samples subjected to HPC. DE expression was acquired considering *in silico* bulk samples obtained summing the expression of all cells belonging to a given cluster and retrieved from co-cultures. DE genes with adj. P value < 0.05 are indicated in red. Selected genes are labelled on the graphs. **b**, Protein-protein interaction network relative to DE genes that were common to all clusters. The network were obtained through STRING, non-connected nodes were included in the representation. Branches in the network are coloured according to the interaction evidence **c,** Volcano plot of DE genes obtained comparing single culture and co-culture samples, both subjected to HPC. DE genes with adj. P value < 0.05 are indicated in red. Selected genes are labelled on the graphs. **d**, STRING protein-protein interaction network relative to DE genes determined as described in panel c. Nodes relative to selected GO terms highlighted in panel e are represented with the same colour code. Non connected nodes were excluded from visualization. Branches in the network are coloured according to the interaction evidence **e**, GO terms (cellular processes) related to DE genes determined comparing single culture and co-culture samples, both subjected to HPC. GO terms were obtained through STRING.
