## Supplementary information for "Modelling Osteoarthritis pathogenesis through Mechanical Loading in an Osteochondral Unit-on-Chip"

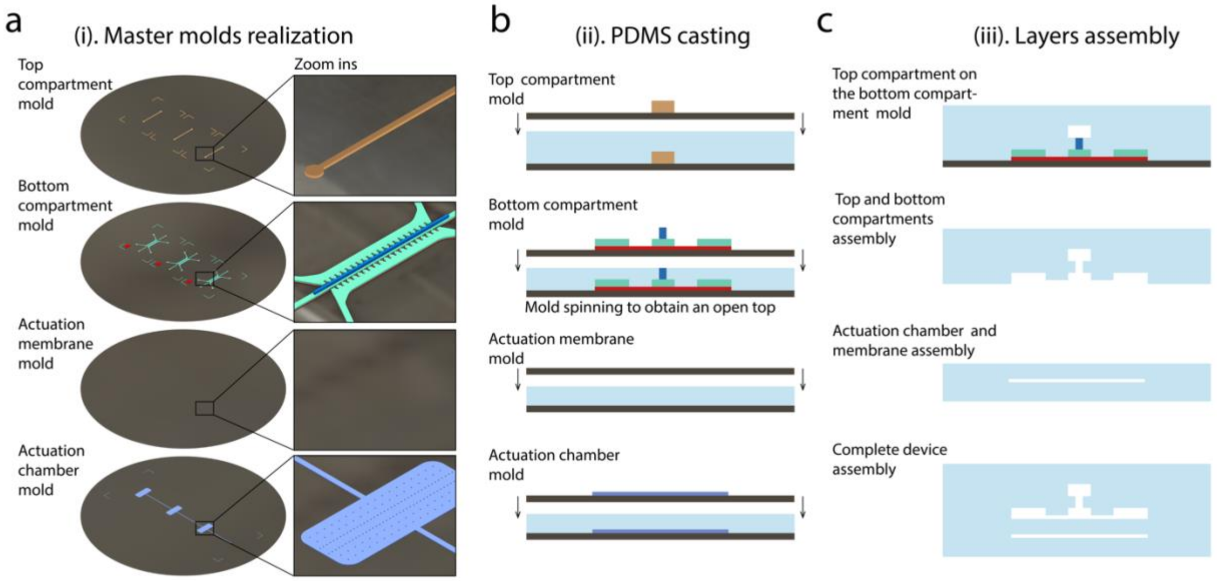

**Supplementary Fig. 1. Device Fabrication protocol. a,** Schematics of the master moulds required to produce the four PDMS device layers. The top compartment mould refers to the NDG. The insets highlight the structures of the different photoresist layers required to obtain the final moulds. The actuation membrane is constituted by a simple PDMS layer produced from a plain silicon wafer. **b,** PDMS casting on the different moulds. The bottom compartment layer is obtained spinning the mould with a spin coater after having poured the PDMS in order to obtain an open top in correspondence of the VBV necking. **c,** Layer assembly. The top compartment layer is peeled of the mould and bonded directly on the bottom compartment layer while this is still attached to its mould. The top compartment – bottom compartment assembly is then peeled of the mould altogether, thus obtaining the VBV tilted-H geometry, and bonded with the actuation chamber-membrane assembly.

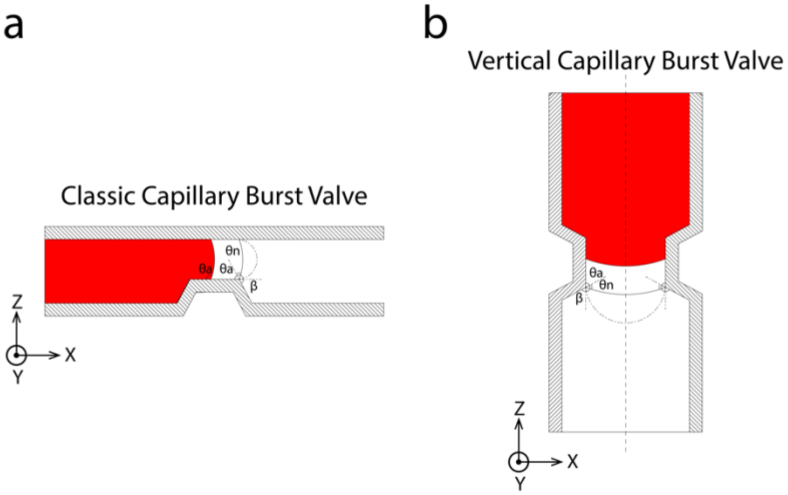

**Supplementary Fig. 2. Classic CBV and VBV working principle.** Assuming a liquid is moving in a narrow channel along the Y direction (i.e. perpendicularly to the represented XZ plane) it can be prevented from invading a neighbouring channel through a striction, or a necking, followed by a sudden aperture which creates an increase in the pressure required for the fluid to proceed in that direction according to the Young-Laplace Equation[1]. This geometry is usually labelled a CBV. The walls of the channel are indicated by the etching, the advancing liquid is portrayed in red. The advancing contact angle is indicated with θa, the angle change caused by the valve with β, and the contact angle that the advancing liquid interfaces assumes with the new wall after the striction with θn = θa – β. The interface required for the liquid to have a contact angle θa with the new interface and resuming advancement is indicated by the dashed line. Given the very limited contribution of gravitational forces at the microscale with respect to the surface tension, responsible for the fluid motion in the channel, it was possible to translate the CBV to a vertical geometry, which we named VBV, and which functions according to the same principle.

***Chambers alignment optimization***

The achievement of monolithic closed channels or partially free-standing structures (as the VBV necking), through the use of photoresists such as SU-8 requires the use of complex and error prone methodologies. These include sacrificial materials that support the structural photoresists until their removal[2], or multi-layer photolithographic processes necessitating a finely tailored selective cross-linking of spatially confined photoresist layers[3]. Achievement of the VBV structures with the abovementioned processes was attempted and abandoned due to the unshapely geometry of obtained features (data not shown).
Given the necessity to align top and bottom compartments of each device, the process precision, accuracy, and efficiency were assessed and optimized. Manually positioning the two compartments might result, in fact, in partially aligned chambers (Supplementary Fig. 3a).
Top compartments with a central channel width of 300 µm, 400 µm, and 500 µm were realized and coupled with bottom compartments with a fixed central channel measuring 300 µm. While a wider top compartment central channel allows an easier centring, minimising the risk of "not covering" the VBV necking area (constituting the contact interface between the compartments), it decreases the portion of the top compartment construct which has a direct interface with the bottom compartment whose dimensions, together with the 100 µm VBV necking, are fixed.
The alignment precision was assess acquiring top view images of assembled devices (Supplementary Fig. 3b). The portion of each central channel which was aligned with the corresponding counterpart was calculated as:

$$Overlapping\ area\ ratio\% = \frac{Overlapping\ area}{Total\ layer\ area} * 100 \qquad (1)$$

Where the *Overlapping area* was defined as the area between the two innermost pillar rows in top view images (regardless of which layer they belonged), while the *Total layer area* was measured as the distance between the pillar rows of each layer (which might slightly differ from the nominal values due to the imaging angle). The product of the *Overlapping area ratio%* of the two layers was adopted as a final criterion to evaluate which configuration maximized the production success.
Values resulted respectively, 82 ± 8 %, 75 ± 11% and 77 ± 6% for the 300 µm, 400 µm, or 500 µm top culture chambers (Supplementary Fig. 3c), being statistically significantly higher in the narrower device version. The top chamber with the narrower central channel (i.e. 300 µm) was therefore adopted in the final design. The assessment was performed with n=6 devices for each configuration. Devices in which the VBV necking area did not overlap with the top compartment central channel were excluded from the analysis. Considering n=30 devices less then 10% of devices were discarded proving the practicability of the fabrication procedure described in Supplementary Fig. 1.

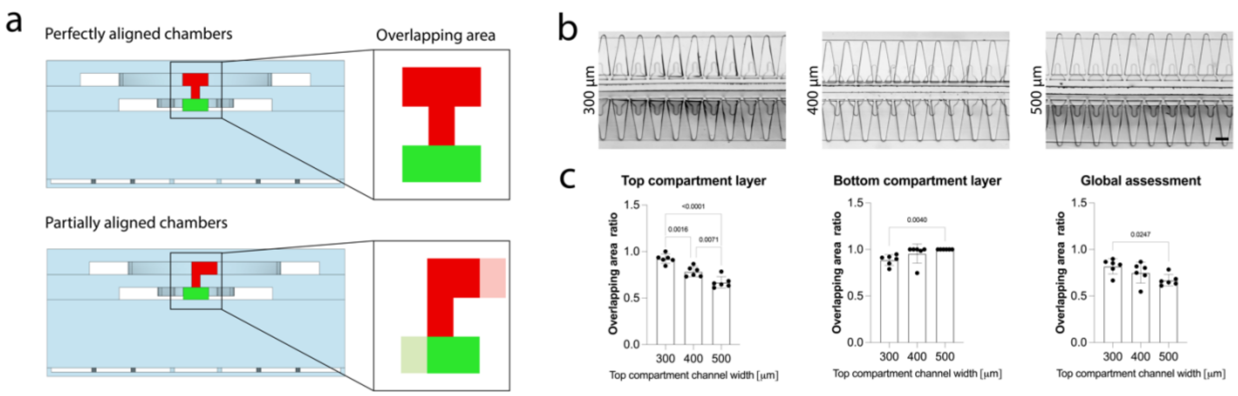

**Supplementary Fig. 3. Top compartment's alignment precision assessment and optimization. a,** Side view schematization of perfectly aligned (top) and partially aligned (bottom) top and bottom compartment . The portion of the aligned central channels is represented in bold colours in the insets while non overlapping areas are depicted in a fainter colour. **b,** Top view brightfield images of aligned top and bottom compartments; DG. For each top compartment geometry three versions were realized, respectively with the central channel of 300 µm, 400 µm, and 500 µm (while the compression chamber gel channel remained of 300 µm) to determine if a wider top channel was necessary to achieve overlapping of top and bottom compartments. Scale bar, 300 µm. **c,** Ratio of the overlapping area with respect to the chamber area calculated as the percentage of one chamber aligned with the other one divided by the total area of the chamber. The Global ratio was calculated as the product of the ratios of top and bottom compartment layers. n=6 devices for each version were considered. Populations' normality was assumed if both Shapiro-Wilk and Kolmogorov-Smirnov tests resulted positive. Statistics by Ordinary One-Way Anova with Tukey's multiple comparison test for normal populations and Kruskal-Wallis test with Dunn's multiple comparison test for non-normal populations. Results are reported as mean ± s.d.

***Determination of devices actuation pressure.***

The actuation pressure, i.e. the pressure necessary to obtain contact of the actuation membrane with the pillars in the bottom compartment was determined with a previously described methodology[4,5].

Actuation chambers were slowly filled with PBS applying a mild continuous positive pressure (i.e. 0.2 Atm) until all air bubbles were removed. Subsequently, top and bottom compartments were filled with blue dye, both the central channel and lateral medium channels. As a result of the blue dye filling the gap between bottom compartment pillars and the actuation membrane, these appear blue at atmospheric pressure. Increasing the pressure in the actuation chamber causes the membrane to deflect upward reducing the gap space until the membrane abuts against the pillars' bottom surface and they appear as white. A correlation between the Mean Grey Intensity measured in correspondence of the pillars and the provided pressure could be established. The actuation pressure was defined as the pressure that at the beginning of the plateau in the Mean Grey intensity vs Pressure curve (i.e. 0.4 Atm, Supplementary Fig. 4a). At least three chambers were considered for each version of the device (Supplementary Fig. 4b). For each chamber the Mean Grey Intensity of three different pillars was measured and results averaged. Images were analysed with ImageJ.

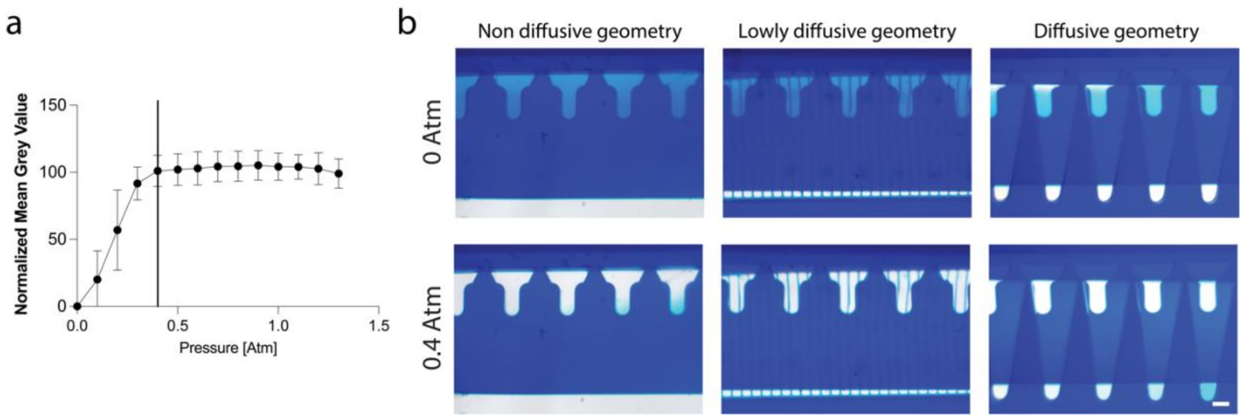

**Supplementary Fig. 4. Determination of the actuation pressure. a,** T-shaped pillars Normalized Mean Grey Intensity versus actuation chamber pressure. The actuation pressure, defined as the pressure necessary to achieve contact between the top surface of the actuation membrane and the bottom surface of the pillars in bottom compartment, was determine with a previously described methodology[4]. Briefly, devices were filled with a blue coloured dye. At atmospheric pressure pillars in the bottom compartment appear blue as a result of the colour filling the gap between pillars and actuation membrane. When the membrane bends upward as a result of the increased pressure in the actuation chamber the gap narrows and pillars start appearing whiter, that is to say the Mean Grey Intensity value increases. Once contact is reached a further increase in pressure does not correspond anymore to an increase in Mean Grey Intensity and a plateau is reached. The actuation pressure (i.e. the onset of the plateau) is indicated by the black vertical line. Results are reported as mean ± s.d. **b,** Examples of top view images of the three device geometries at atmospheric pressure (0 Atm) and at the actuation pressure (0.4 Atm). Scale bar 100 µm.

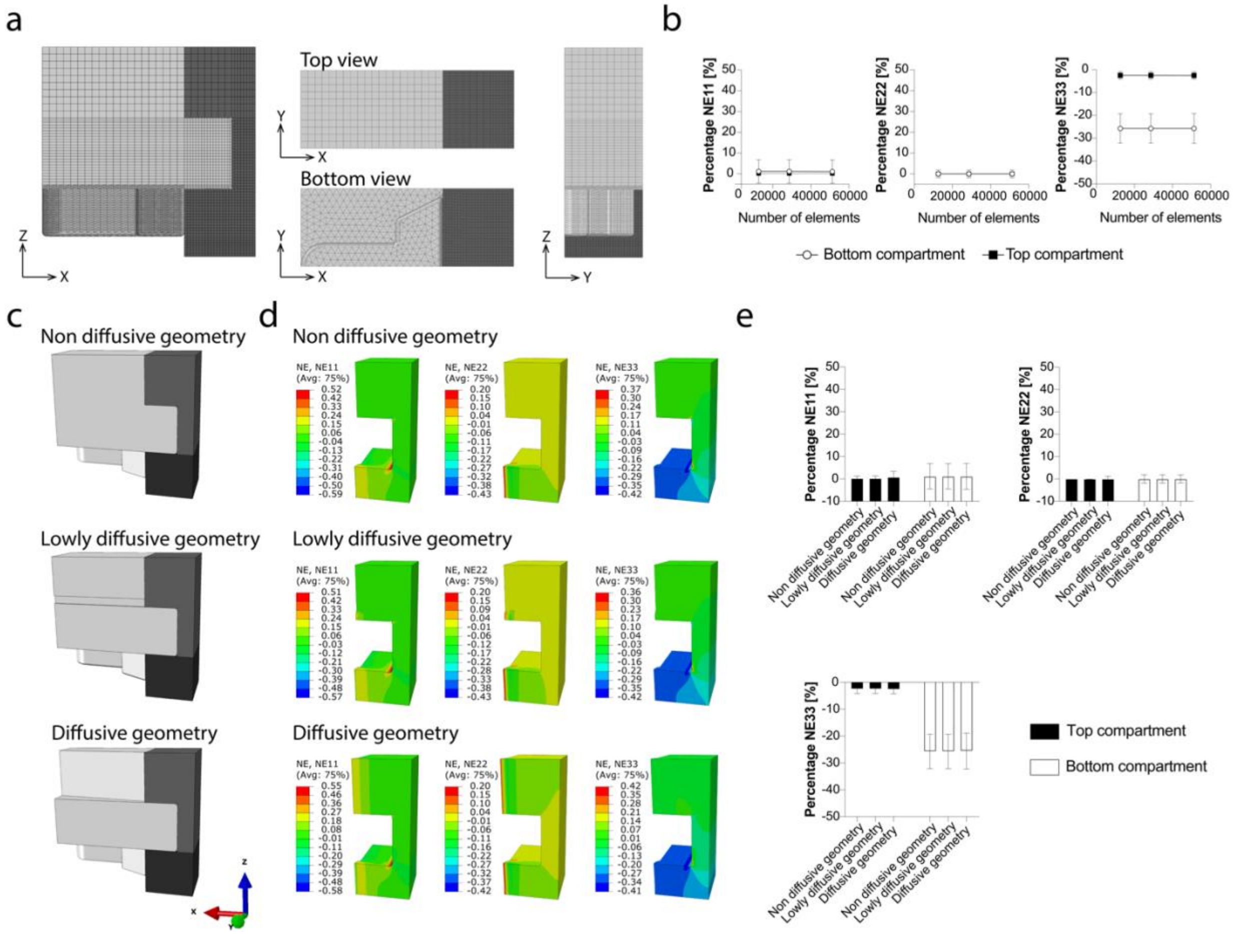

**Supplementary Fig. 5. FE model setup. a,** Orthogonal plane views of the 3D geometry adopted in computations. The adopted elements' mesh is represented. PDMS regions are depicted in light grey, the top construct is represented in grey and the bottom constructs in black. **b,** Mesh sensitivity analysis: nominal strains along the principal directions, NE11, NE22, and NE 33 were evaluated in the top (comprehensive of the necking area) and bottom 3D constructs as a function of the number of elements of the constructs. FE models of devices with all three top compartment configurations, namely Non diffusive geometry, Lowly diffusive geometry, and Diffusive geometry were considered. Values, extrapolated from the elements' centroids are reported as mean ± s.d. **c,** Geometries adopted in computations complete of constructs (in dark grey and black) and PDMS structures (in light grey). **d,** Contour plots of the nominal strains along the principal directions for the three device versions. No noticeable differences could be detected in the strain field in the three cases with the exception of a locally increased lateral expansion in the top compartment for the Diffusive geometry configuration. **e,** Average NE11, NE22, and NE33 in the different geometrical configurations. Values, extrapolated from the elements' centroids are reported as mean ± s.d.

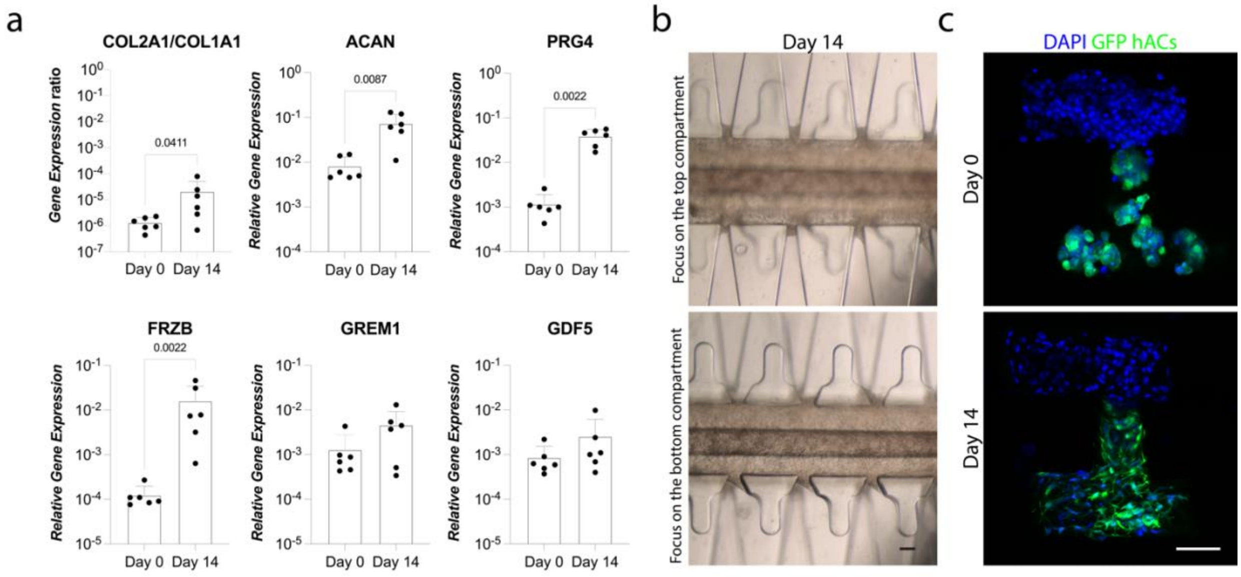

**Supplementary Fig. 6. Establishment of a bi-layer mature human cartilaginous construct on-chip. a,** The achievement of mature cartilaginous constructs after 14 days of static culture in the device was confirmed through the gene expression of known chondrogenic markers[4]. Gene expression was quantified through RT-qPCR. (n≥6 biologically independent samples from n=2 donors). Statistical significance was determined by Mann-Whitney test. Populations' normality was assumed if both Shapiro-Wilk and Kolmogorov-Smirnov tests resulted positive. P values < 0.05 are reported on the graph. All genes expression values were normalized for GAPDH expression, values are reported as mean + s.d. **b,** Representative brightfield pictures of hACs statically cultured in devices for 14 days (DG configuration of the top chamber). After 2 weeks of culture cells are still confined in the central channels. Similar results were obtained with more than 60 devices. Scale bar, 100 µm. **c,** Immunofluorescence images of constructs sections immediately after seeding (Day 0) or after 14 days of static culture (Day 14). Constructs' layers have a direct interface in correspondence of the VBV necking and maintain a stratified structure throughput the culture period. hACs were seeded in the top compartment, GFP hACS in the bottom one. DAPI, 4′,6-diamidino-2-phenylindole, is represented in blue, GFP in green (Images from n=3 biologically independent devices from one donor for each compartment were considered in the analyses). Scale bar, 100 µm.

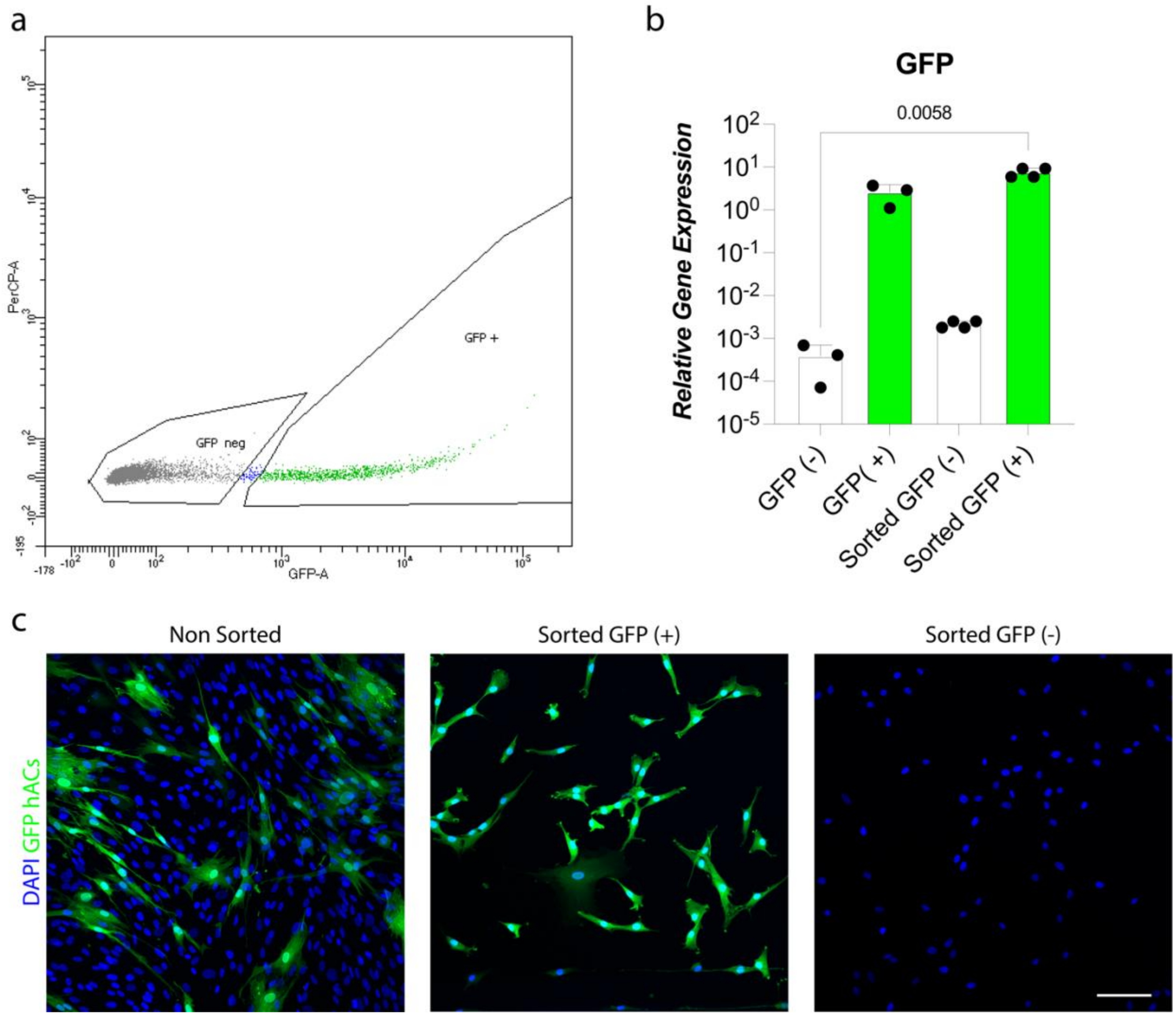

**Supplementary Fig. 7. hACs from top and bottom compartments can be separated based on GFP expression. a,** Example of populations selection during sorting. **b,** The GFP gene expression of sorted populations is similar to that of pure populations. GFP expression was quantified by RT-qPCR. Pure populations i.e. populations which were never in contact with each other are indicated as GFP (+) and GFP (-), populations sorted after culture and constructs enzymatic digestion are indicated as Sorted GFP (-) and Sorted GFP (+), (n≥3 biologically independent samples from n=1 donor for each population). Statistics by Kruskal-Wallis test with Dunn's multiple comparison test. Populations' normality was assumed if both Shapiro-Wilk and Kolmogorov-Smirnov tests resulted positive. P values < 0.0

5 are reported on the graph. All genes expression values were normalized for GAPDH expression, values are reported as mean + s.d. **c,** Representative immunofluorescence images of cellular populations obtained after constructs digestion demonstrating successful population separation based on GFP (At least n=3 independent constructs, from n=1 donor for each population were considered for each condition). Constructs were digested and sorted (or not) and cells plated in 2D on an IBIDI 8 well µ-plate. Cells were left to adhere for 48 hours then fixed and imaged. DAPI is represented in blue, GFP in green. Scale bar, 100 µm.

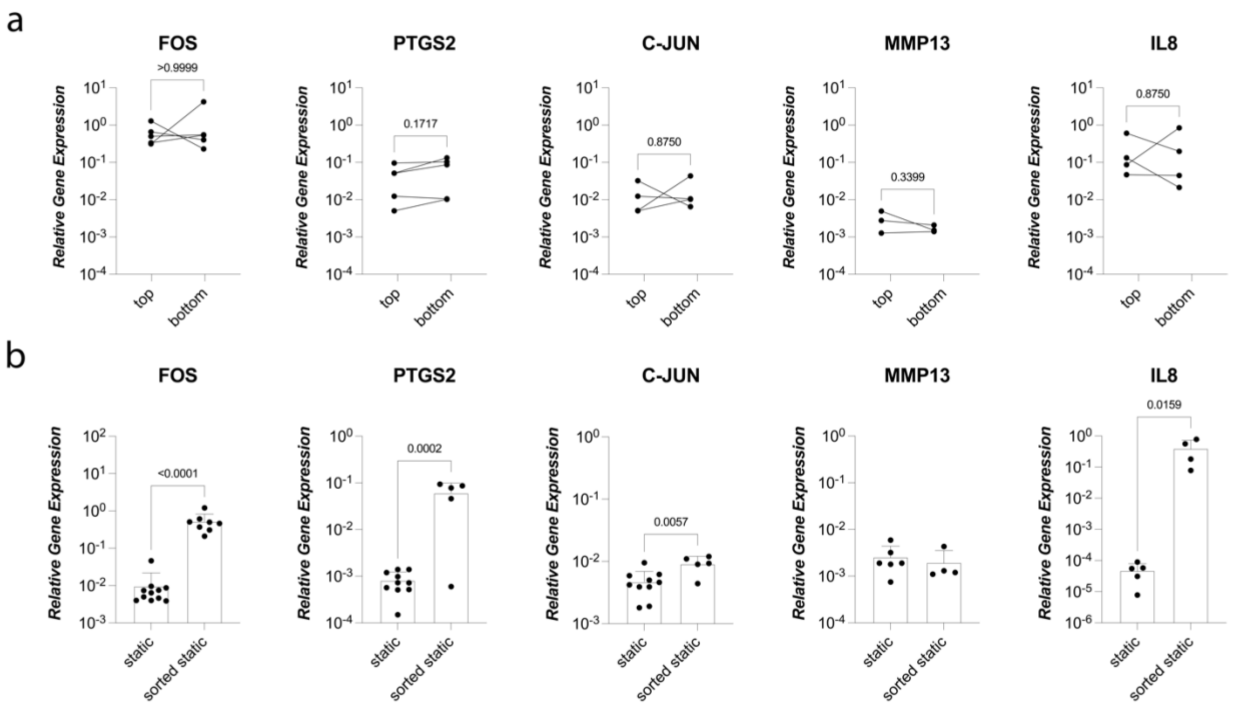

**Supplementary Fig. 8. Cartilaginous tissues gene expression baselines in top and bottom compartments and sorting effect on gene expression. a,** Cartilaginous tissues cultured statically in top and bottom compartment for 21 days do not show differences in gene expression as quantified by RT-qPCR. (n≥4 biologically independent samples from n≥2 donors; samples cultured in the same device were comparted directly). Statistical significance was determined by paired t-test for normal populations and Wilcoxon test for non-gaussian populations respectively. P values are reported on the graph, no statistically significant differences (i.e. P value < 0.05) were detected. Gene expression levels referring to top and bottom constructs coming from the same chamber are connected by black lines. **b,** Constructs' enzymatic digestion and sorting affect the gene expression of all genes of interest except for MMP13 as quantified by RT-qPCR (n≥4 biologically independent samples from n≥2 donors). Statistics by unpaired two tailed t-test for normal populations and Mann-Whitney test for non-Gaussian ones. P values < 0.05 are reported on the graph. All genes expression values were normalized for GAPDH expression, values are reported as mean + s.d. Populations' normality was assumed if both Shapiro-Wilk and Kolmogorov-Smirnov tests resulted positive.

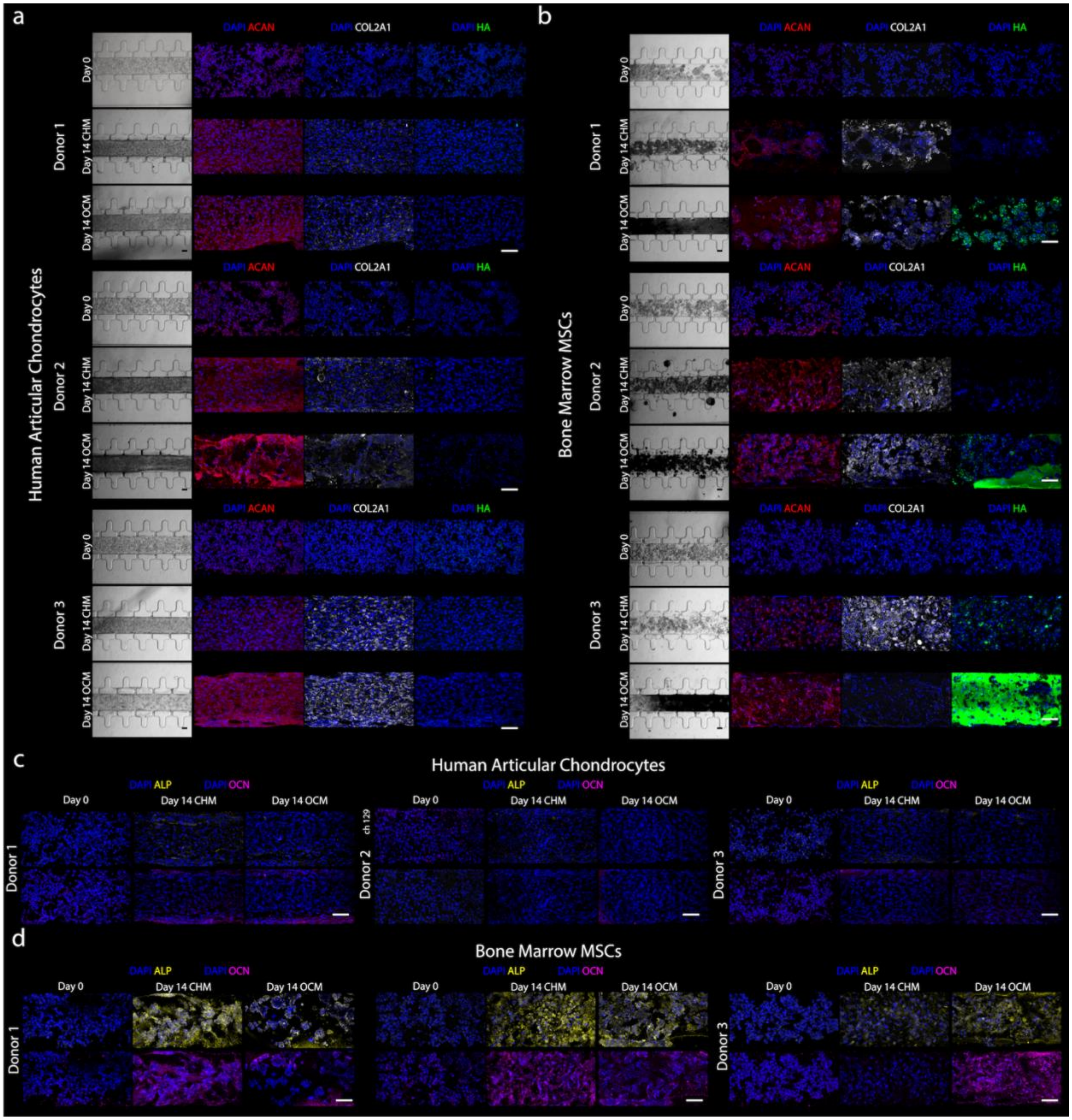

**Supplementary Fig. 9. Characterization of hACs and MSCs-based constructs upon CHM and OCM exposure.** **a**, **b**, Brightfield (left) and immunofluorescence (right) images of hACs (**a**) and MSCs (**b**) constructs cultured in CHM or OCM for 14 days (n≥2 biologically independent samples from n=5 hACs donors and n=3 MSCs were considered in analyses, images from 3 donors are reported) Switching to the OCM formulation does not alter hACs chondrogenic potential (which remains subjected to high donor to donor variability) but consistently leads to MSCs depositing hydroxyapatite (HA). Hyaline cartilage markers ACAN and COL2A1 are represented in red and white respectively. HA is represented in green. Day 0 constructs were adopted as negative controls. **c**, **d**, Immunofluorescence images of hACs (**c**) and MSCS (**d**) constructs cultured in CHM or OCM for 14 days (n≥.1 biologically independent samples from n=3 donors were considered in analyses, images from 3 donors are reported). ALP is represented in yellow, OCN in magenta. In all images DAPI (represented in blue) was used as nuclear counterstaining. Scale bar, 100 µm.

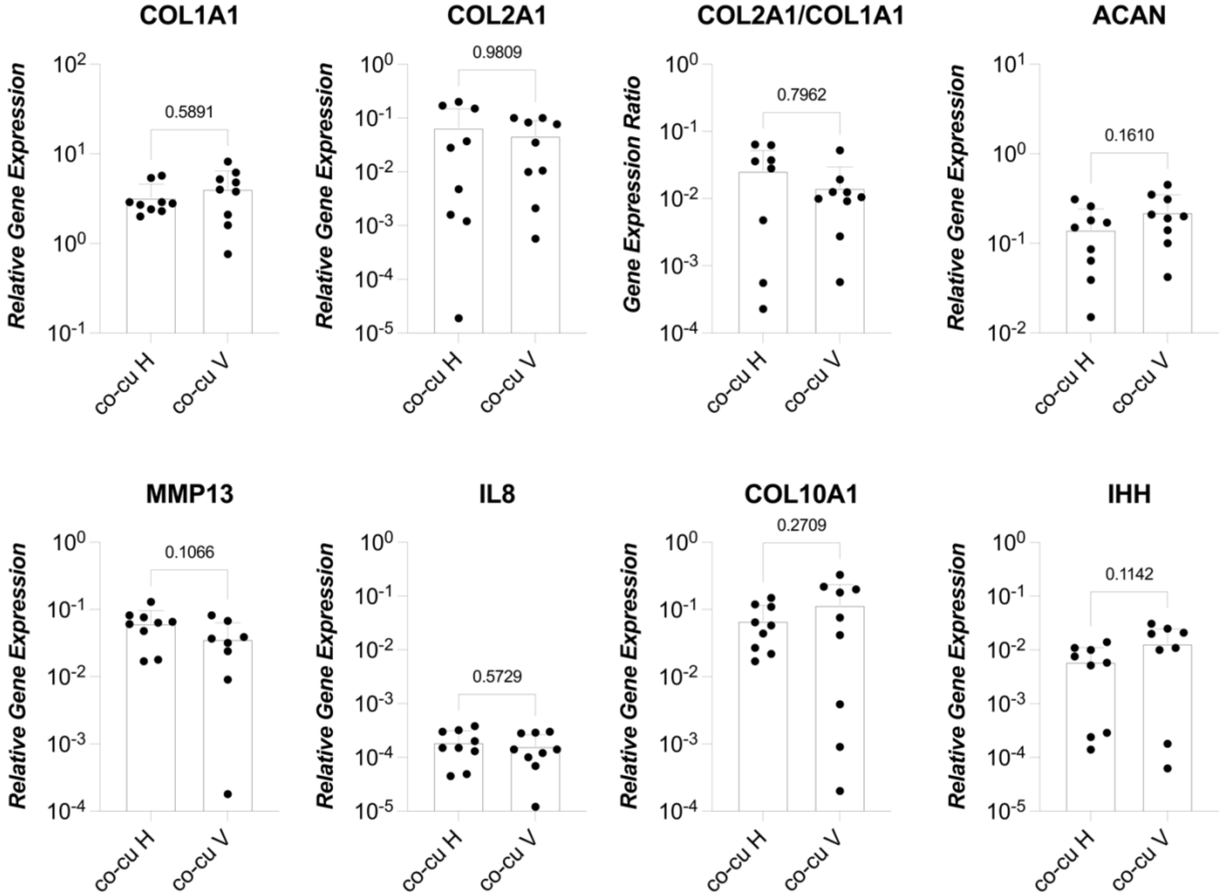

**Supplementary Fig. 10. Comparison of the gene expression in Horizontal (H) and vertical (V) osteochondral constructs.** The gene expression of osteochondral construct does not depend on the cartilaginous and mineralized tissues disposition. While our newly introduced VBV featuring platform is suitable to provide cartilage and subchondral tissues with physio-pathologically relevant mechanical stimuli a horizontal disposition facilitates the observation of the interface between the two constructs. Gene expression was quantified by RT-qPCR. (n=3 biologically independent samples from n=3 independent donors/experiments were considered in analyses, gene expression reefers to whole osteochondral tissues comprehensive of hACs and MSCs in co-culture analyzed together). Statistical significance was determined by paired t-test for normal populations and Mann-Whitney test for non-gaussian populations respectively. P values are reported on the graph, no statistically significant differences (i.e. P value < 0.05) were detected. All genes expression values were normalized for GAPDH expression, values are reported as mean + s.d. Populations' normality was assumed if both Shapiro-Wilk and Kolmogorov-Smirnov tests resulted positive.

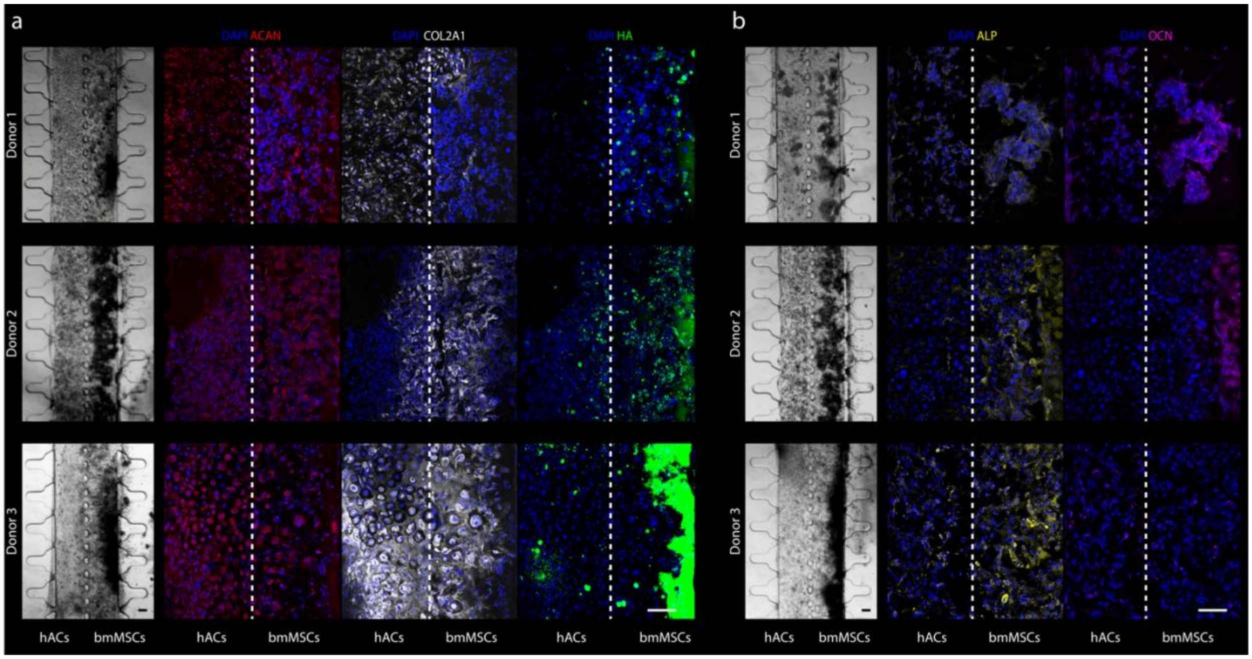

**Supplementary Fig. 11. Maturation of hACs and MSCs co-cultures into osteochondral constructs.**
**a**, Brightfield (left) and immunofluorescence (right) images of hACs and MSCs co-cultured for 14 days in OCM. (n≥2 biologically independent samples from n=6 hACs donors and n=4 MSCs were considered in analyses, images from 3 independent experiments are reported). Hyaline cartilage markers ACAN and COL2A1 are represented in red and white respectively. HA is represented in green. Despite a high donor to donor variability biphasic osteochondral-like constructs with a mineralized layer and a cartilaginous layer are obtained could be obtained consistently. Differently from single cultures cells positive for HA are present also in the hACs compartment. MSCs compartment mineralization appears not to be homogeneous but to increase further away from hACs. **b**, Brightfield (left) and immunofluorescence (right) images of hACs and MSCs co-cultured for 14 days in OCM. (n≥1 biologically independent samples from n=3 hACs donors and n=3 MSCs were considered in analyses, images from 3 independent experiments are reported). ALP represented in yellow, and OCN portrayed in magenta arrear confined to the MSCs compartment. In all images DAPI (represented in blue) was used as nuclear counterstaining. Scale bar, 100 µm.

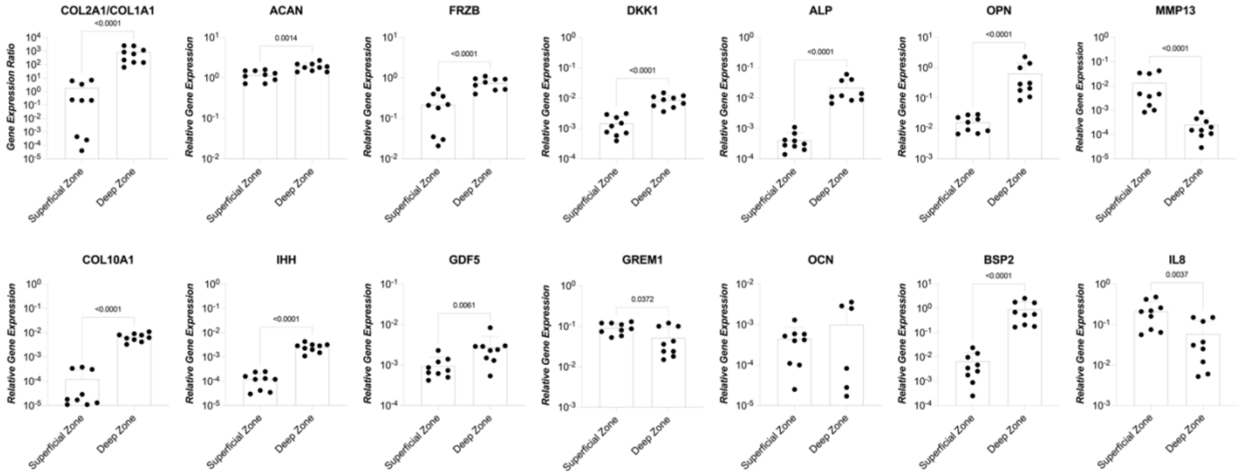

**Supplementary Fig. 12. Genes expression in superficial and deep zones of human knee cartilage.** The expression of genes used as markers of chondrogenesis, hypertrophy, Wnt and BMP activation, or bone maturation was assessed in ex-vivo knee cartilage from human donors. Cartilage samples were harvested from the dorsal portion of the femoral condyle of patients undergoing knee replacement for OA, from areas subjected to minimal load bearing and without signs of cartilage degradation or fibrillation of the surface. Cartilage from the superficial zone and from the deep zone were collected and analyzed separately (samples from n=3 patients were considered, for each patient n=3 separate areas were harvested). Tissues were digested enzymatically; gene expression was quantified through RT-qPCR. Statistical significance was determined by paired t-test for normal populations and Wilcoxon test for non-gaussian populations respectively. P values are reported on the graph. Differences in gene expression were detected between superficial and deep zones. All genes expression values were normalized for GAPDH expression, values are reported as mean + s.d. Populations' normality was assumed if both Shapiro-Wilk and Kolmogorov-Smirnov tests resulted positive.

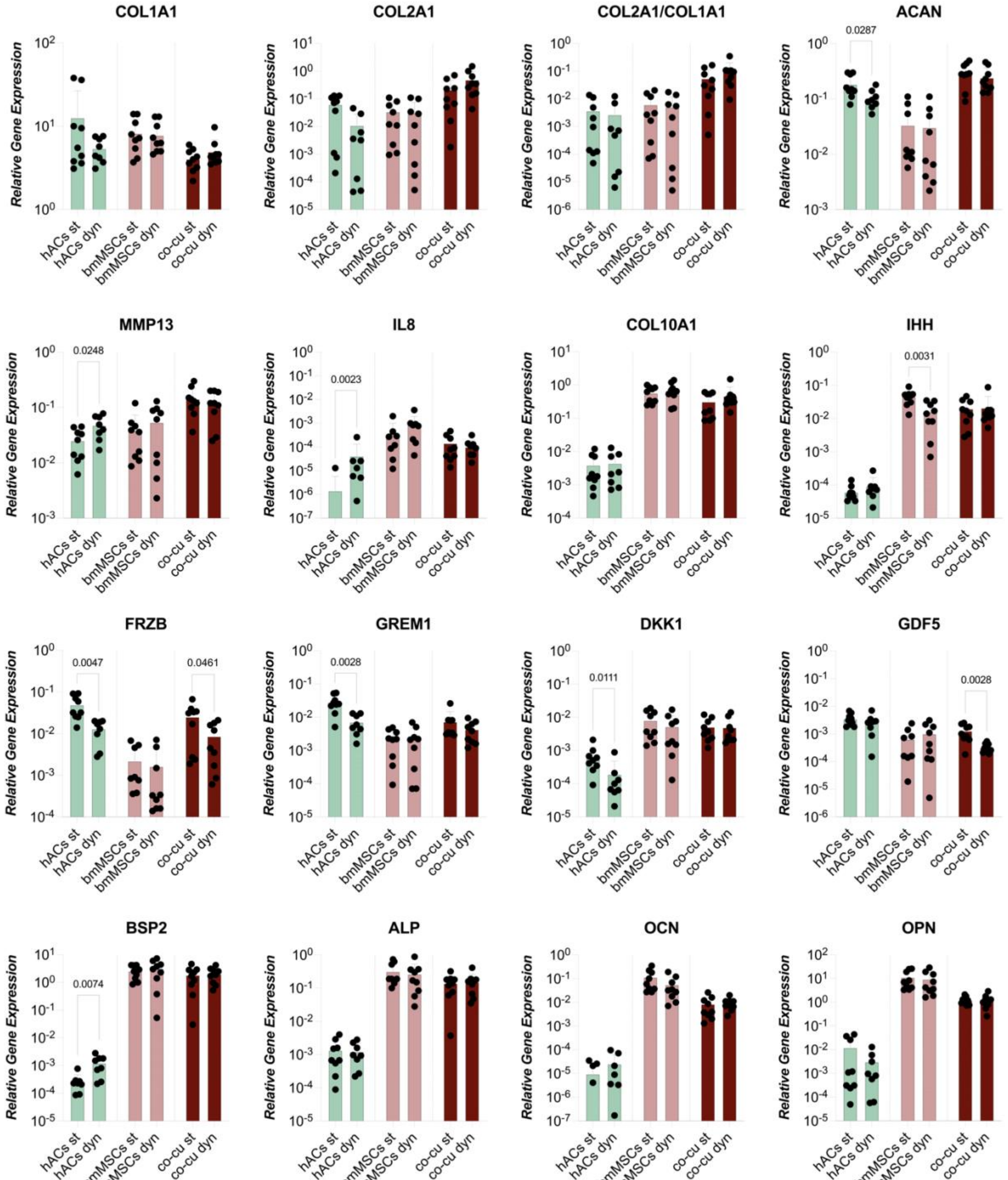

**Supplementary Fig. 13. Effect of mechanical cyclical loading on single culture hACs and MScs and on osteochondral constructs bulk gene expression.** When cultured in OCM cyclical mechanical loading has a catabolic, inflammatory, and hypertrophic effect (in terms of gene expression) on hACs based cartilaginous constructs while it has very limited influence of bmMSCs based mineralized tissues. The effect of hyperphysiological cyclical loading was also assessed in whole osteochondral constructs where, however, only the cartilaginous tissues are exposed to hyperphysiological compression. Notably osteochondral constructs anabolic and catabolic markers expression is higher than that of single cultures indicating a greater degree of remodeling, possibly due to inter tissues crosstalk. Gene expression was quantified by RT-qPCR. (n=9 biologically independent samples from n=3 independent donors/experiments for each condition were considered in analyses, co-cu gene expression reefers to whole osteochondral tissues comprehensive of hACs and MSCs in co-culture analyzed together). Statistical significance was determined by paired t-test for normal populations and Wilcoxon test for non-gaussian populations respectively. P values $P \leq 0.05$ are reported on the graphs. All genes expression values were normalized for GAPDH expression, values are reported as mean + s.d. Populations' normality was assumed if both Shapiro-Wilk and Kolmogorov-Smirnov tests resulted positive.

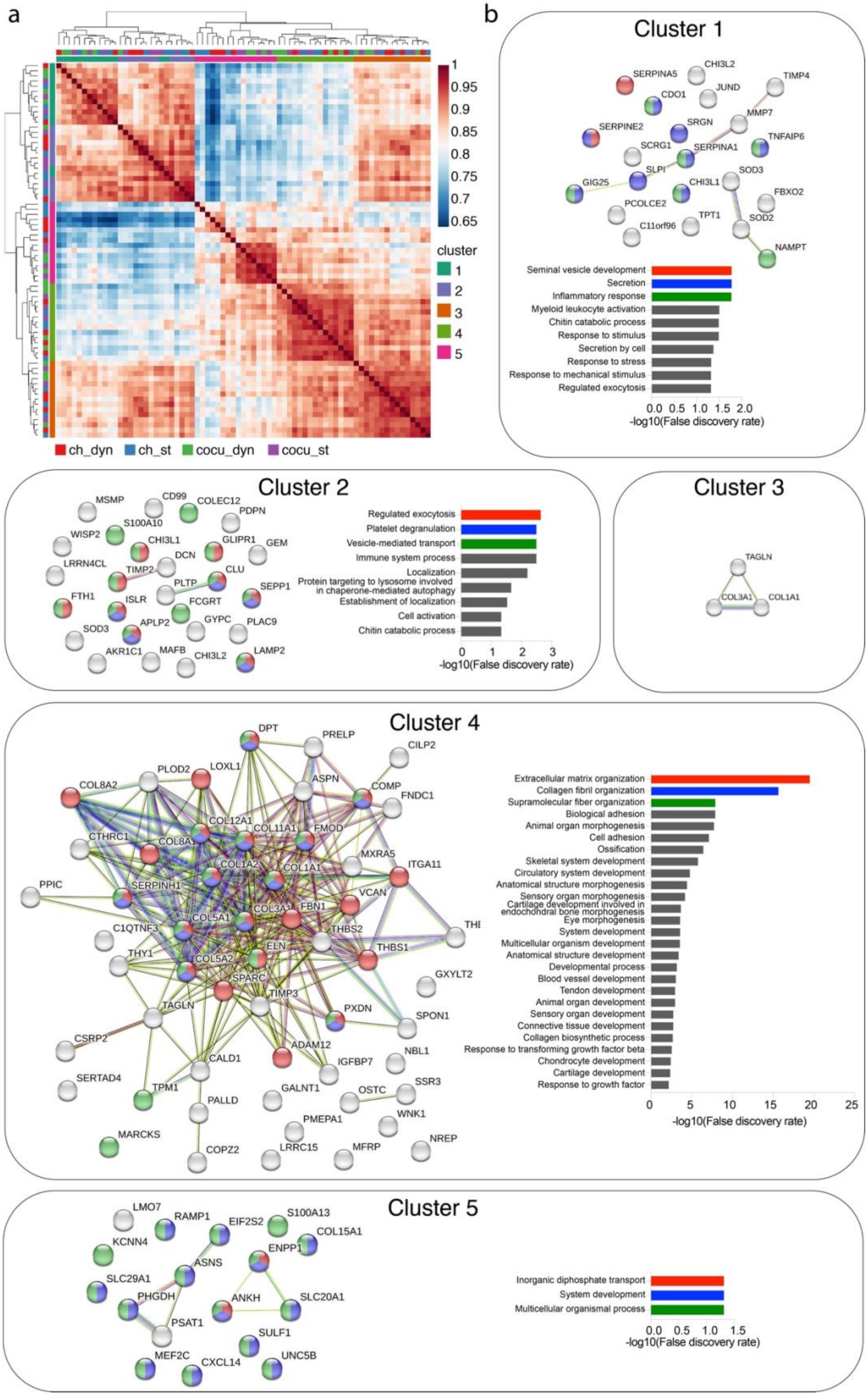
a
1
0.95
0.9
0.85
0.8
0.75
0.7
0.65
cluster
1
2
3
4
5
ch_dyn
ch_st
cocu_dyn
cocu_st
b
Cluster 1
SERPINA5
CHI3L2
TIMP4
JUND
CDO1
SRGN
MMP7
SERPINE2
TNFAIP6
SCRG1
SERPINA1
SOD3
SLPI
GIG25
CHI3L1
FBXO2
PCOLCE2
TPT1
SOD2
C11orf96
NAMPT
Seminal vesicle development
Secretion
Inflammatory response
Myeloid leukocyte activation
Chitin catabolic process
Response to stimulus
Secretion by cell
Response to stress
Response to mechanical stimulus
Regulated exocytosis
0.0 0.5 1.0 1.5 2.0
-log10(False discovery rate)
Cluster 2
MSMP
CD99
COLEC12
PDPN
S100A10
WISP2
CHI3L1
GLIPR1
GEM
DCN
LRRN4CL
TIMP2
CLU
PLTP
SEPP1
FTH1
ISLR
FCGRT
APLP2
GYPC
PLAC9
SOD3
AKR1C1
MAFB
CHI3L2
LAMP2
Regulated exocytosis
Platelet degranulation
Vesicle-mediated transport
Immune system process
Localization
Protein targeting to lysosome involved in chaperone-mediated autophagy
Establishment of localization
Cell activation
Chitin catabolic process
0 1 2 3
-log10(False discovery rate)
Cluster 3
TAGLN
COL3A1
COL1A1
Cluster 4
DPT
PRELP
CILP2
PLOD2
LOXL1
ASPN
COMP
COL8A2
FNDC1
COL12A1
COL11A1
FMOD
CTHRC1
COL8A1
COL1A2
COL1A1
MXRA5
ITGA11
PPIC
SERPINH1
VCAN
COL3A1
FBN1
C1QTNF3
COL5A1
THBS2
THBS1
THY1
ELN
COL5A2
GXYLT2
SPARC
TIMP3
TAGLN
PXDN
SPON1
CSRP2
ADAM12
CALD1
IGFBP7
NBL1
SERTAD4
SSR3
TPM1
GALNT1
OSTC
PALLD
WNK1
PMEPA1
MARCKS
NREP
LRRC15
MFRP
COPZ2
Extracellular matrix organization
Collagen fibril organization
Supramolecular fiber organization
Biological adhesion
Animal organ morphogenesis
Cell adhesion
Ossification
Skeletal system development
Circulatory system development
Anatomical structure morphogenesis
Sensory organ morphogenesis
Cartilage development involved in endochondral bone morphogenesis
Eye morphogenesis
System development
Multicellular organism development
Anatomical structure development
Developmental process
Blood vessel development
Tendon development
Animal organ development
Sensory organ development
Connective tissue development
Collagen biosynthetic process
Response to transforming growth factor beta
Chondrocyte development
Cartilage development
Response to growth factor
0 5 10 15 20 25
-log10(False discovery rate)
Cluster 5
LMO7
RAMP1
EIF2S2
S100A13
COL15A1
KCNN4
ENPP1
ASNS
SLC29A1
PHGDH
ANKH
SLC20A1
PSAT1
SULF1
MEF2C
CXCL14
UNC5B
Inorganic diphosphate transport
System development
Multicellular organismal process
0.0 0.5 1.0 1.5
-log10(False discovery rate)

**Supplementary Fig. 14. Characterization of identified cell clusters. a**, Hierarchical clustering of gene expression correlation profiles. Assigned clusters and samples experimental conditions are reported on the edges according to the colour codes indicated in the legend. **b**, Protein-protein interaction network and enriched GO terms of molecular processes for the differentially expressed genes of each cluster as determined with STRING. Branches in the network are coloured according to the interaction evidence. Nodes are either grey or coloured according to their relation to the GO terms with the lowest false discovery rate as reported in respective bar graphs. Non connected network nodes are not reported in the images. No statistically significant enriched GO terms relative to cellular processes could be found for Cluster 3.

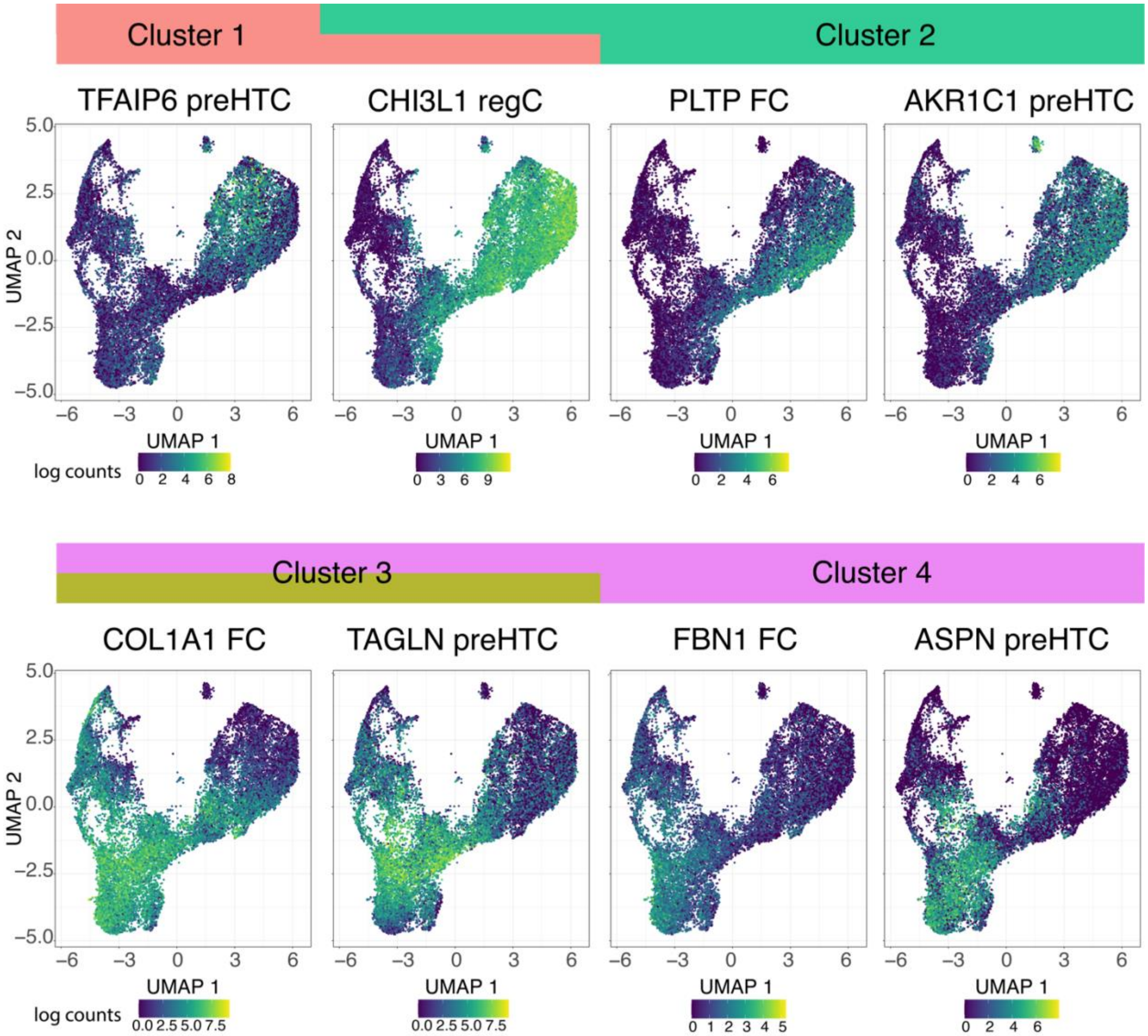

**Supplementary Fig. 15. Correspondence of clusters and chondrocytes subpopulation markers.** Feature plots of selected marker genes for clusters 1-4 that were previously defined as hACs sub-population markers[6,7]. CHI3L1 was expressed both by cell from cluster 1 and 2. Cluster 3 had no cluster specific markers but represented cells with an intermediate state between cluster 2 and 4.

, Supplementary Fig. 15.

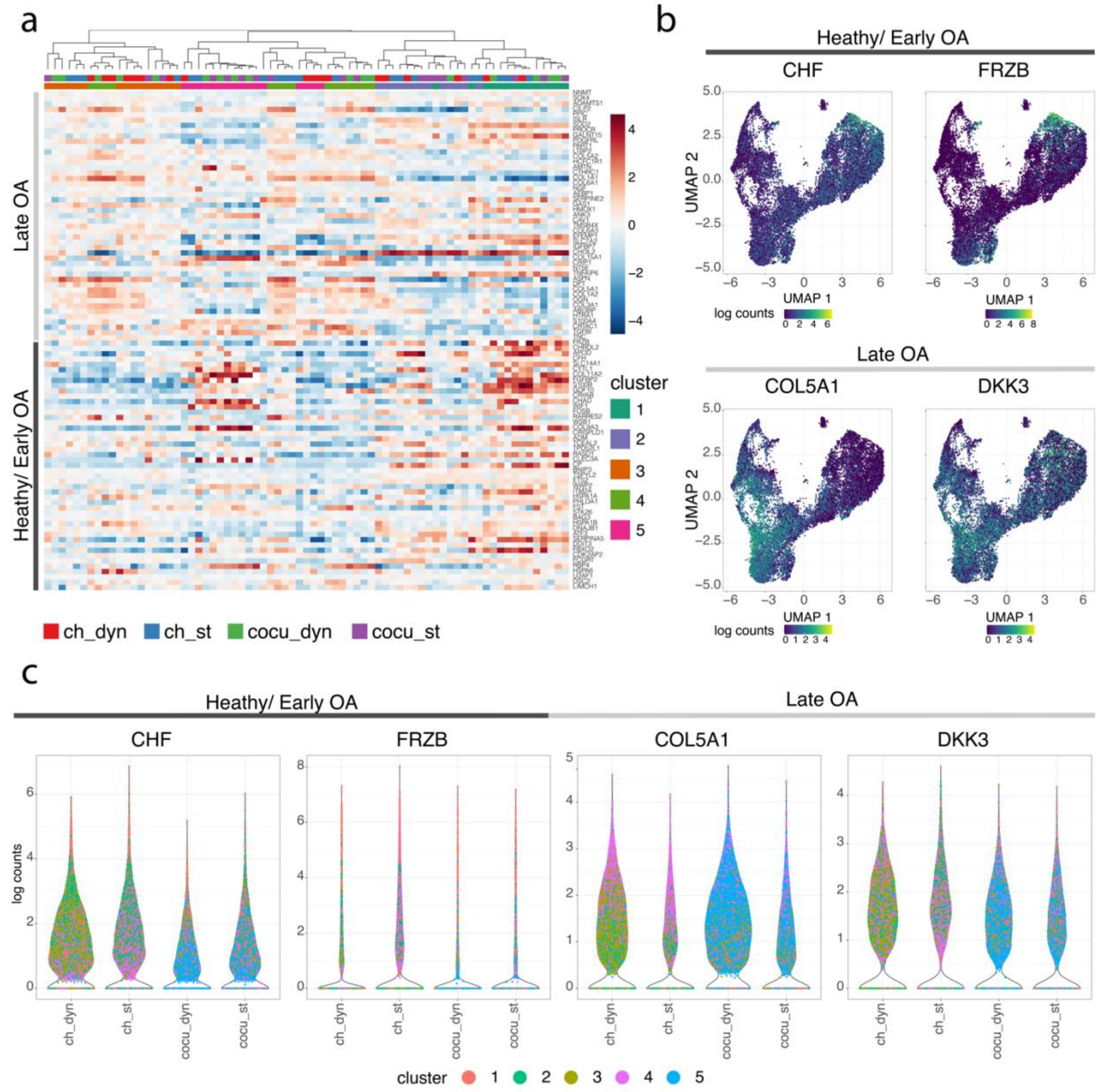

**Supplementary Fig. 16. Clusters correlation with early stage and late-stage OA markers. a**, Hierarchical clustering based on the 50 top negatively correlated (i.e. associated to early disease stages) and the 50 top positively correlated (i.e. associated with advanced disease stages) genes along PC2 in a previously performed PCA analysis of the transcriptomes of chondrocytes from OA cartilage samples with various disease stages[6]. Culture conditions and clusters of belonging are colour coded at the top of the heatmap **b**, feature plots and **c**, Violin plots and of selected early OA markers (i.e. CHF and FRZB), and late OA markers (i.e.COL5A1 and DKK3). Single points in violin plots are coloured according to the cluster to which the specific cell was assigned.

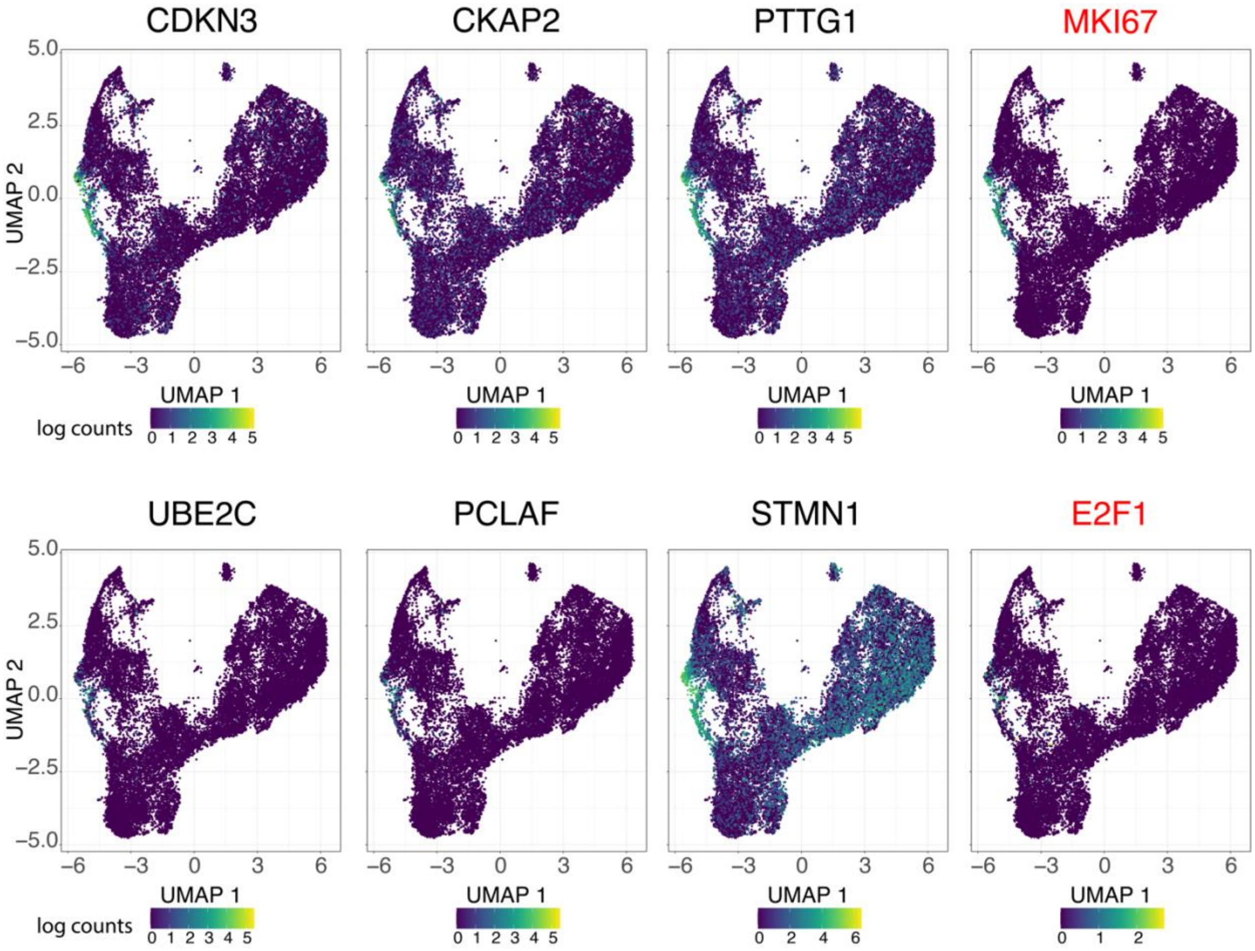

**Supplementary Fig. 17. Expression of CPCs markers.** Feature plots of previously identified CPCs markers[7] and of the proliferation markers MKI67 and E2F1[8] which are reported in red.

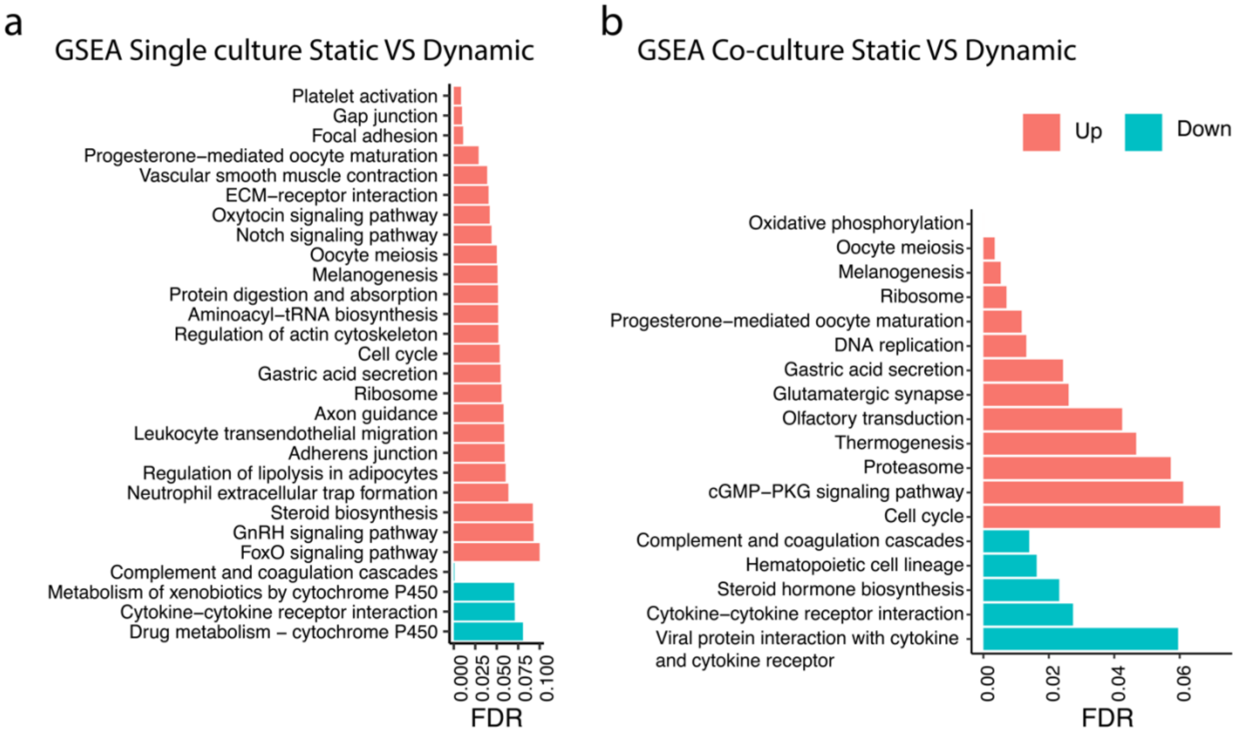

**Supplementary Fig. 18 KEGG pathways gene sets enrichment analyses from in silico bulk samples. a,** Single culture. **b,** Co-culture

### ***References***

1. Cho, H., Kim, H. Y., Kang, J. Y. & Kim, T. S. How the capillary burst microvalve works. *J. Colloid Interface Sci.* **306**, 379–385 (2007).
2. Yun, K. S. & Yoon, E. Fabrication of complex multilevel microchannels in PDMS by using three-dimensional photoresist masters. *Lab Chip* **8**, 245–250 (2008).
3. Moser, Y., Forti, R., Jiguet, S., Lehnert, T. & Gijs, M. A. M. Suspended SU-8 structures for monolithic microfluidic channels. *Microfluid. Nanofluidics* **10**, 219–224 (2011).
4. Occhetta, P. *et al.* Hyperphysiological compression of articular cartilage induces an osteoarthritic phenotype in a cartilage-on-a-chip model. *Nat. Biomed. Eng.* (2019). doi:10.1038/s41551-019-0406-3
5. Marsano, A. *et al.* Beating heart on a chip: a novel microfluidic platform to generate functional 3D cardiac microtissues. *Lab Chip* **16**, 599–610 (2016).
6. Ji, Q. *et al.* Single-cell RNA-seq analysis reveals the progression of human osteoarthritis. *Ann. Rheum. Dis.* **78**, 100–110 (2019).
7. Wang, X. *et al.* Comparison of the major cell populations among osteoarthritis, Kashin-Beck disease and healthy chondrocytes by single-cell RNA-seq analysis. doi:10.1038/s41419-021-03832-3
8. Whitfield, M. L., George, L. K., Grant, G. D. & Perou, C. M. Common markers of proliferation. (2006). doi:10.1038/nrc1802
